# A pan-genome method to determine core regions of the *Bacillus subtilis* and *Escherichia coli* genomes

**DOI:** 10.1101/2020.06.11.147629

**Authors:** Granger Sutton, Gary B. Fogel, Bradley Abramson, Lauren Brinkac, Todd Michael, Enoch S. Liu, Sterling Thomas

**Affiliations:** J. Craig Venter Institute; Natural Selection, Inc.; Noblis, Inc.

## Abstract

Synthetic engineering of bacteria to produce industrial products is a burgeoning field of research and application. In order to optimize genome design, designers need to understand which genes are essential, which are optimal for growth, and locations in the genome that will be tolerated by the organism when inserting engineered cassettes. We present a pan-genome based method for the identification of core regions in a genome that are strongly conserved at the species level. We show that these core regions are very likely to contain all or almost all essential genes. We assert that synthetic engineers should avoid deleting or inserting into these core regions unless they understand and are manipulating the function of the genes in that region. Similarly, if the designer wishes to streamline the genome, non-core regions and in particular low penetrance genes would be good targets for deletion. Care should be taken to remove entire cassettes with similar penetrance of the genes within cassettes as they may harbor toxin/antitoxin genes which need to be removed in tandem. The bioinformatic approach introduced here saves considerable time and effort relative to knockout studies on single isolates of a given species and captures a broad understanding of the conservation of genes that are core to a species.

**Importance:** The pan-genome approach presented in this paper can be used to determine core regions of a genome and has many possible applications. Synthetic engineering design can be informed by which genes/regions are more conserved (core) versus less conserved. The level of conservation of adjacent non-core genes tends to define cassettes of genes which may be part of a pathway or system that can inform researchers about possible functional significance. The pattern of gene presence across the different genomes of a species can inform the understanding of evolution and horizontal gene acquisition. The approach saves considerable time and effort relative to laboratory methods used to identify essential genes in species.

## Introduction

Over the last decade, considerable interest has been directed towards the determination of a minimal bacterial cell, making use of a short genome consisting of only essential genes for viability. The *Mycoplasma mycoides* JCVI-syn3.0 is a case example of synthetic engineering to design and build a genome that contains a streamlined gene set essential for cell viability and cell replication (1). However, the identification of “essential” genes - those genes that are critical for cell viability and replication - takes considerable time and effort in a laboratory setting and is usually determined with respect to one reference genome under one set of specific growth conditions. For instance, Kobayashi et al. (2) and Koo et al. (3) experimentally and computationally determined the minimal gene set in the gram-positive bacterium *Bacillus subtilis*. Of the ∼4,100 genes of the type strain, a total of 271 genes for Kobayashi and 257 genes for Koo were shown to be essential. These essential genes were further categorized in terms of cell metabolism and enzymatic capability. Additionally for the ∼4400 genes in the gram-negative bacteria *Escherichia coli*, Goodall et al. (4), Baba et al. (5), and Yamazaki et al. (6), it was determined that 414 genes were essential to strain K-12.

Experimental studies to determine such essential genes are time consuming and often restricted to a single environmental condition using a single strain of the species. In addition, these approaches also knock out one gene at a time. As such, genes with multiple copies with redundant functions are often not considered as essential following knockout, as their additional copy is able to maintain cellular function. In other words, a viable organism would not result from deleting all but the experimentally determined essential genes from the genome. Another peculiarity of the single knockout essential genes is that pairs or cassettes of genes which can be removed and still have a viable organism are labeled essential because removal of just one gene is lethal. For example, removing the methylation gene(s) without removing the restriction digestion enzyme genes from the restriction mechanism results in cell death but the cell survives if the entire system is removed. This is likewise true for toxin/antitoxin systems.

While it is possible to define “essential” genes relative to viability, another larger question remains; which genes define a species? While specific phenotypes can vary across strains, in general a species seems to require some minimal set of genes to not only survive in the laboratory but to thrive in its natural environment. In contrast, some strains may have retained or acquired some genes which improve survival for specific niches. Pan-genome analysis of large sets of diverse strains from the same species provide the structure for determining what “core” genes are present in all or almost all strains versus what genes are only present in a limited subset of genomes. These core genes should determine the baseline phenotype (capabilities and traits) of a species. We assert that truly essential genes should typically be a subset of core genes given that core genes presumably confer fitness advantages that are not necessarily essential but have been retained for a reason over evolutionary time. Previous pan-genome studies (7) have shown that species tend to only tolerate the placement of noncore genes between what we call “core regions” and not within those core regions. Here we define a pan-genome “core gene” as a gene which is present in at least 95% of the strains of a species. A “core region” on the genome of a given strain is the minimal region containing a set of adjacent core genes (and no noncore genes) which are adjacent in all or almost all strains in a species. The reason an organism might constrain a core region rather than just core genes is that the region may include regulatory mechanisms such as operons, which allows for co-expression of multiple functionally associated genes, or regulons which would be disrupted with the insertion of other genes. We believe that conservation of core regions in species indicates resistance to insertion of genes through evolution or through human-mediated genetic engineering.

Here we present a pan-genome based calculation of core regions for *B. subtilis* spp. *subtilis* and for *E. coli*. These core regions are compared to previous experimentally determined essential genes from the literature. Our bioinformatics approach saves considerable time and effort in determining core regions and can broaden our understanding of core genomic loci for these specific species. The pan-genome approach outlined here provides a method to determine a set of core regions and their associated metabolic functions which help define a bacterial species/subspecies. This goes beyond determining which genes are essential for survival in a defined laboratory setting and addresses what genes are conserved for a specific species. We would expect that all truly essential genes for the species/subspecies would be a subset of the core genes/regions since core genes would encompass genes responsible for providing a fitness advantage in environmental conditions as well as being essential for viability. This approach automates computational prediction of core genes/regions as an alternative to laborious experimental methods to determine core genes through knockout studies. The proper identification of core genes/regions will greatly improve the effectiveness of synthetic biology and improve our understanding of prokaryotic diversity and phylogeny.

## Results

For *B. subtlis*, both Kobayashi et al. (2) and Koo et al. (3) used similar single knockout methods to determine “essential” genes when grown in LB at 37°C. Koo identified 257 essential genes while Kobayashi identified 271 essential genes. The union of these two sets results in 305 essential genes (Table 1). The 305 genes were mapped to the pan genome graph (PGG) clusters using the RefSeq genome NC_000964.3 (BioSample SAMEA3138188). Interestingly through this mapping, 16 of the essential genes were not identified as core genes. We believe only 9 of these 16 genes are truly essential. Gene *wapI/yxxG* (cluster 4769 present in 39 of 108 genomes) is an antitoxin for the *wapA* toxin gene which is adjacent to it (present in 85 of 108 genomes) (8). Gene *rttF/yqcF* (cluster 4590 present in 46 of 108 genomes) and gene *rtbE/yxxD* (cluster 4772 present in 53 of 108 genomes) are also the antitoxin of a cognate toxin-antitoxin pair (9). Gene *yezG* (cluster 4411 present in 43 of 108 genomes) is also the toxin for a cognate toxin–antitoxin pair (10). Gene *sknR/yqaE* (cluster 4643 present in 34 of 108 genomes) is part of a phage-like region which, if removed would still allow *B. subtilis* to remain viable (11) possibly because it is another antitoxin or similar mechanism. Genes *bsuMA/ydiO* (cluster 4838 present in 24 of 108 genomes) and *bsuMB/ydiP* (cluster 4839 present in 24 of 108 genomes) are part of a prophage region of about 15 genes in 48 genomes which includes *ydiR* and *ydiS* which are type-2 restriction enzymes. These are not essential genes but they are essential if the restriction enzymes are present (12).

**Table 1.**
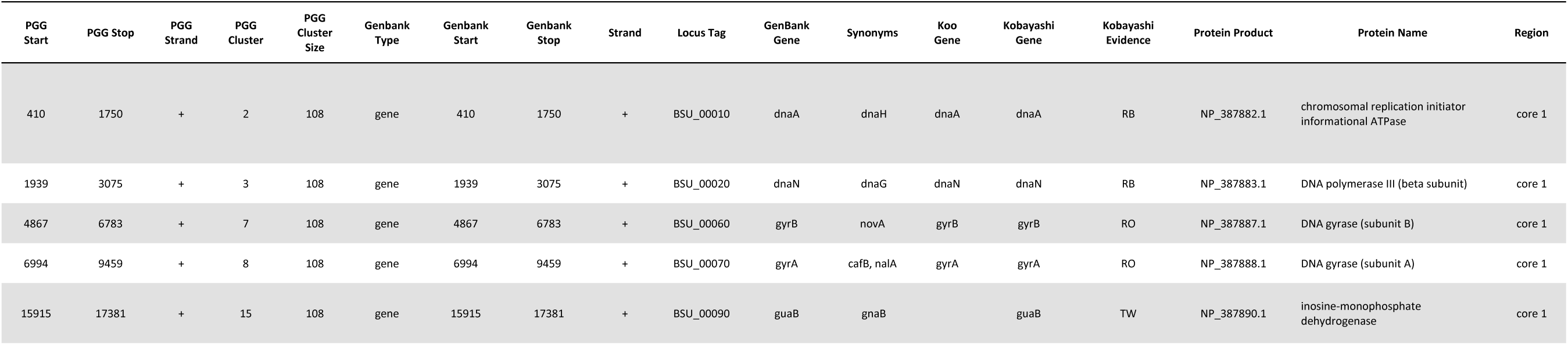
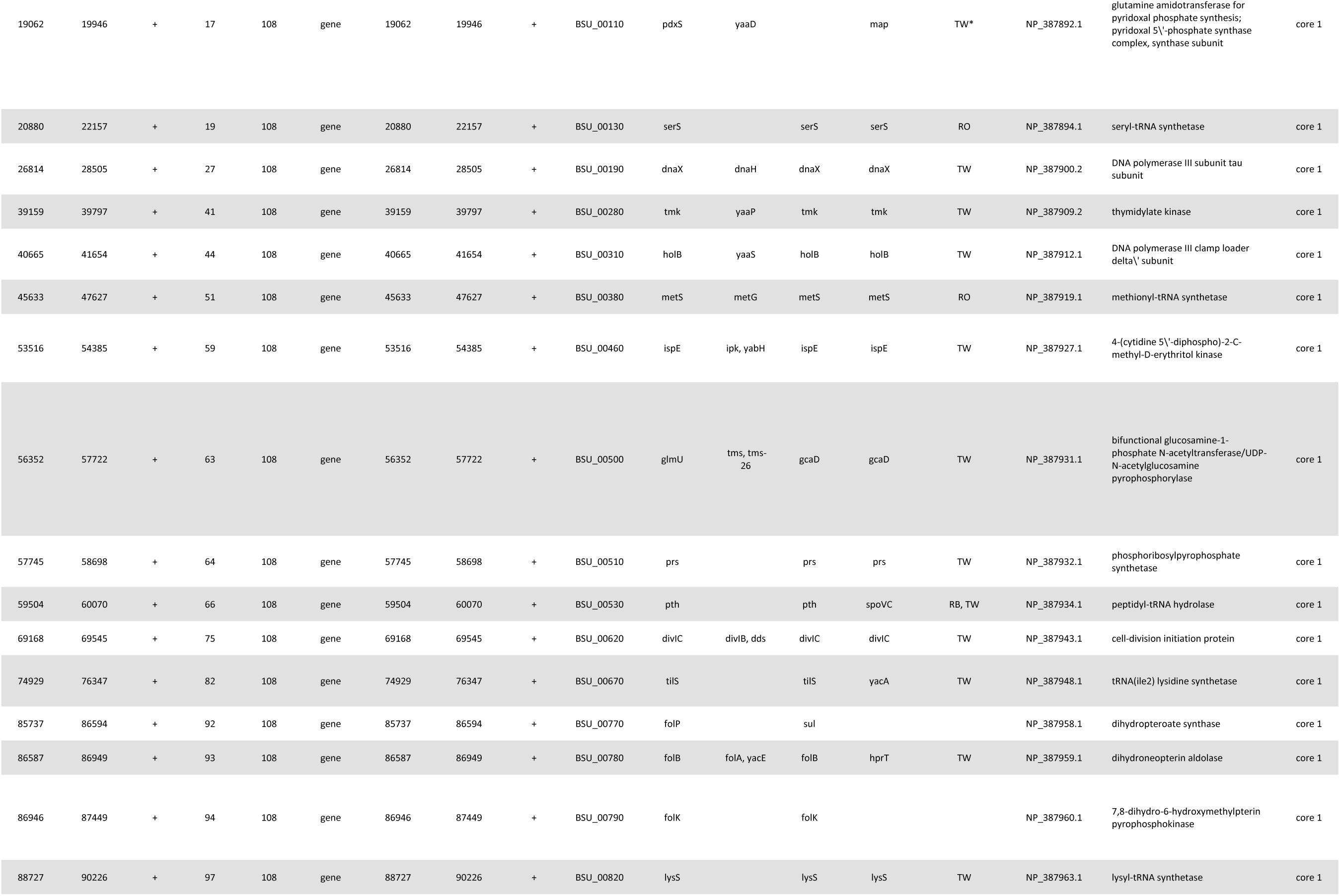
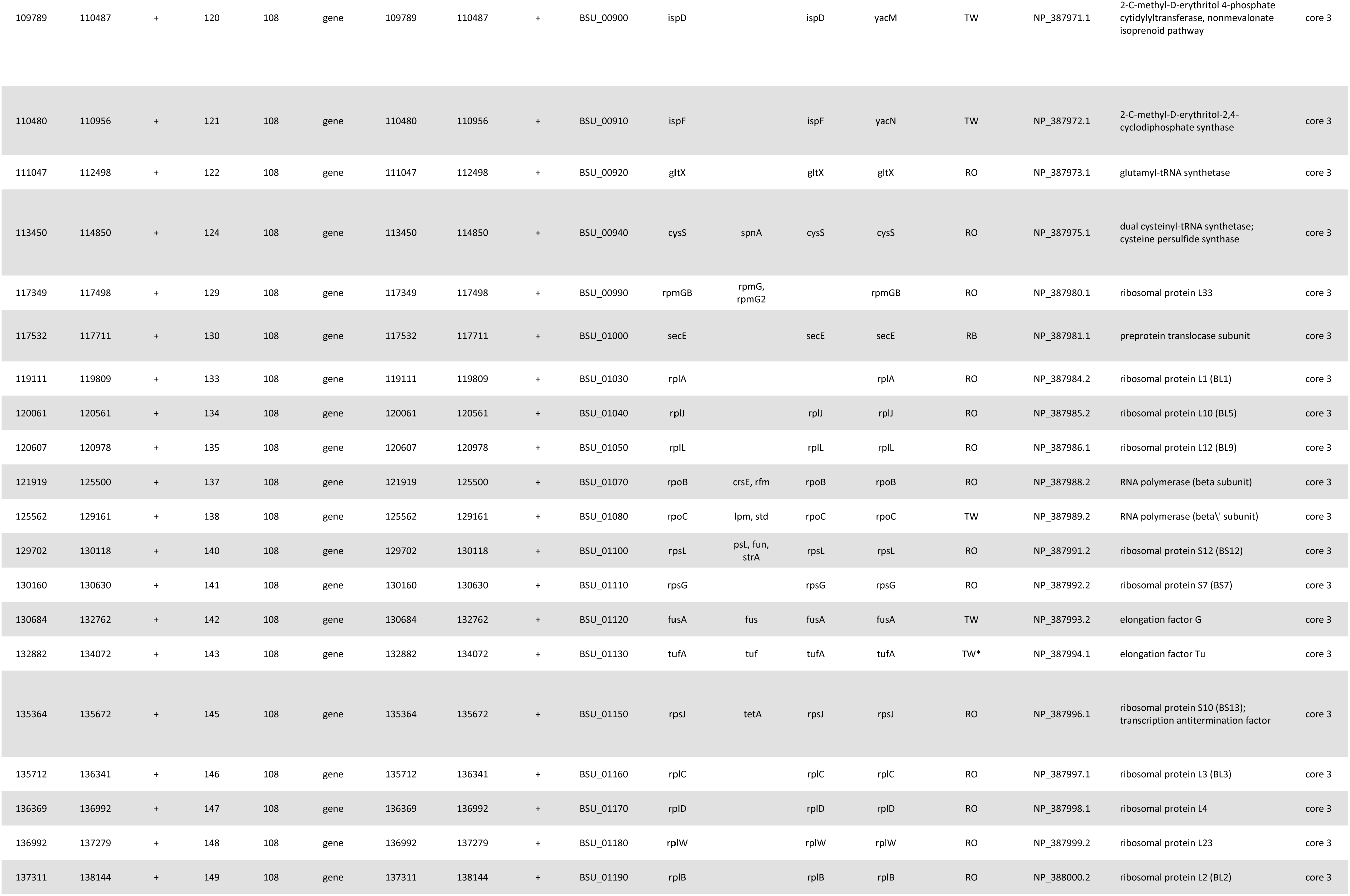
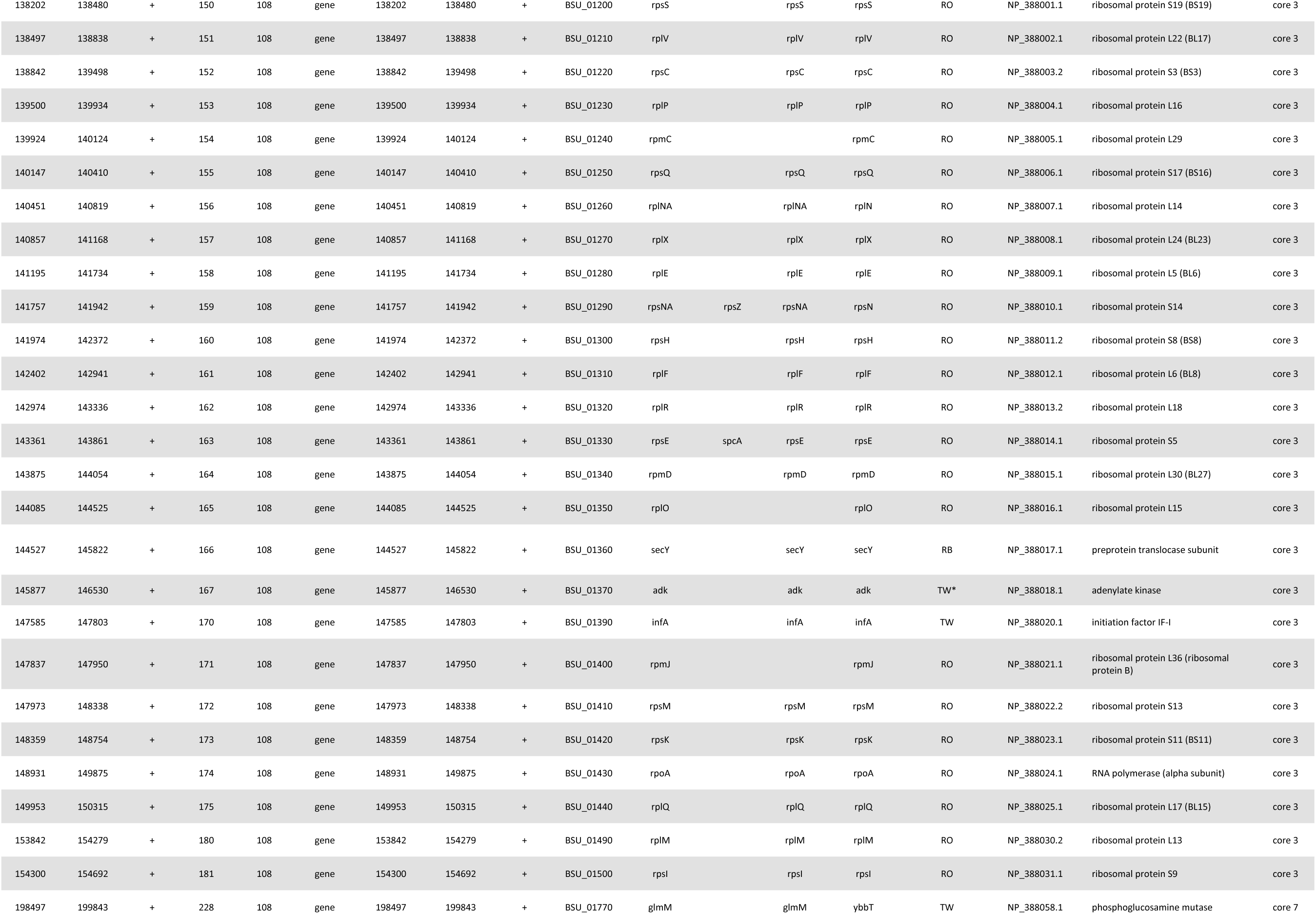
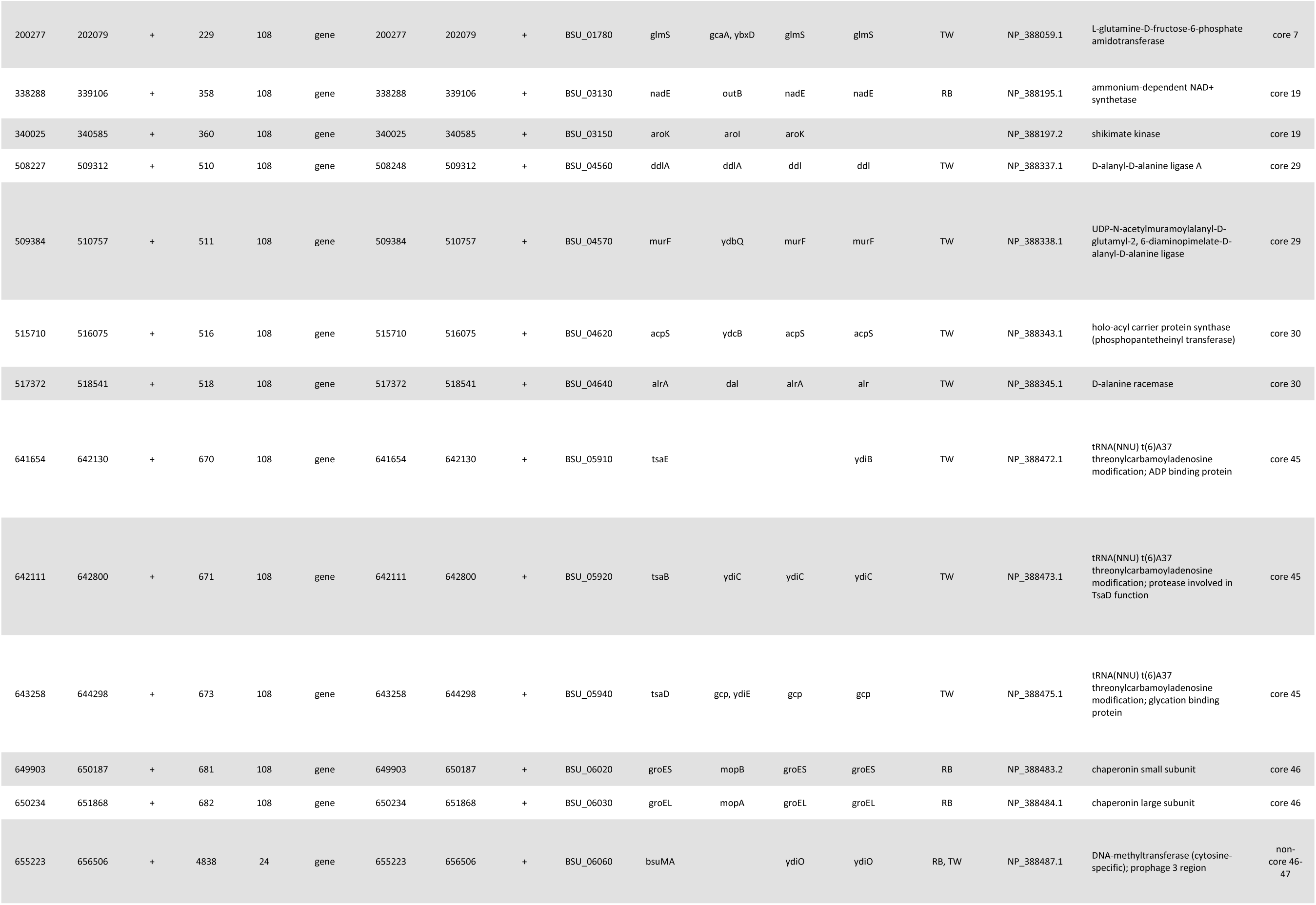
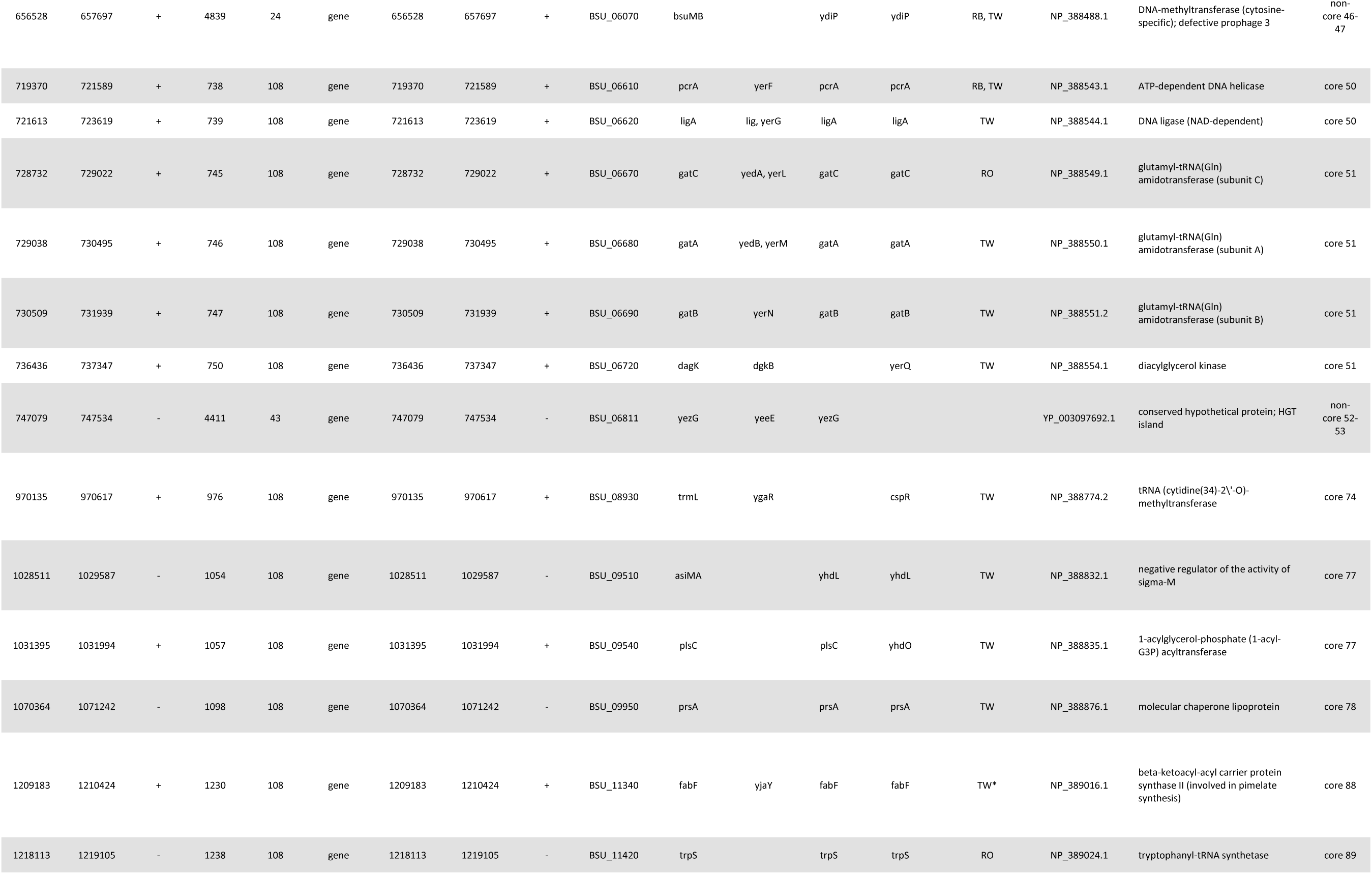
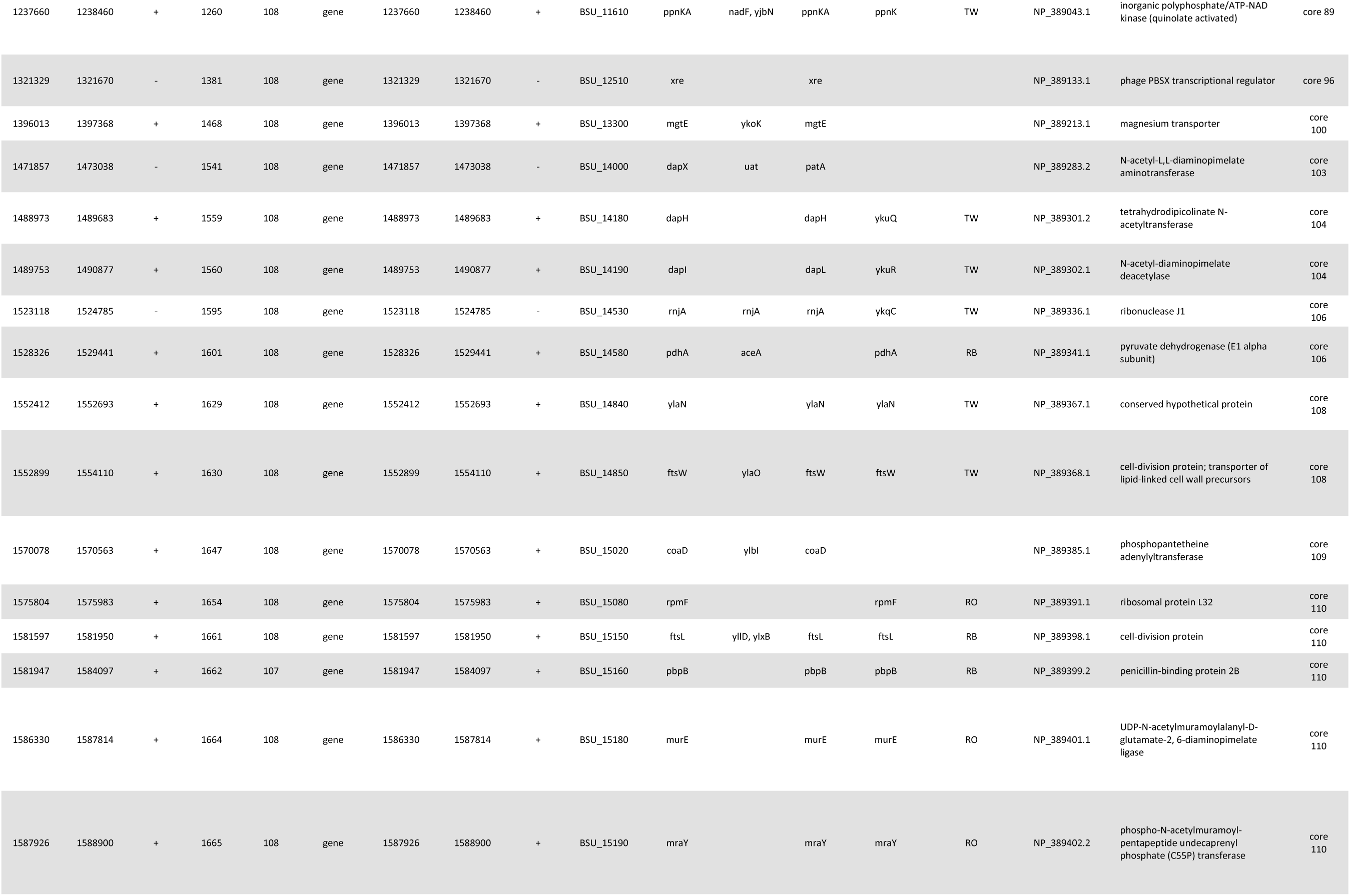
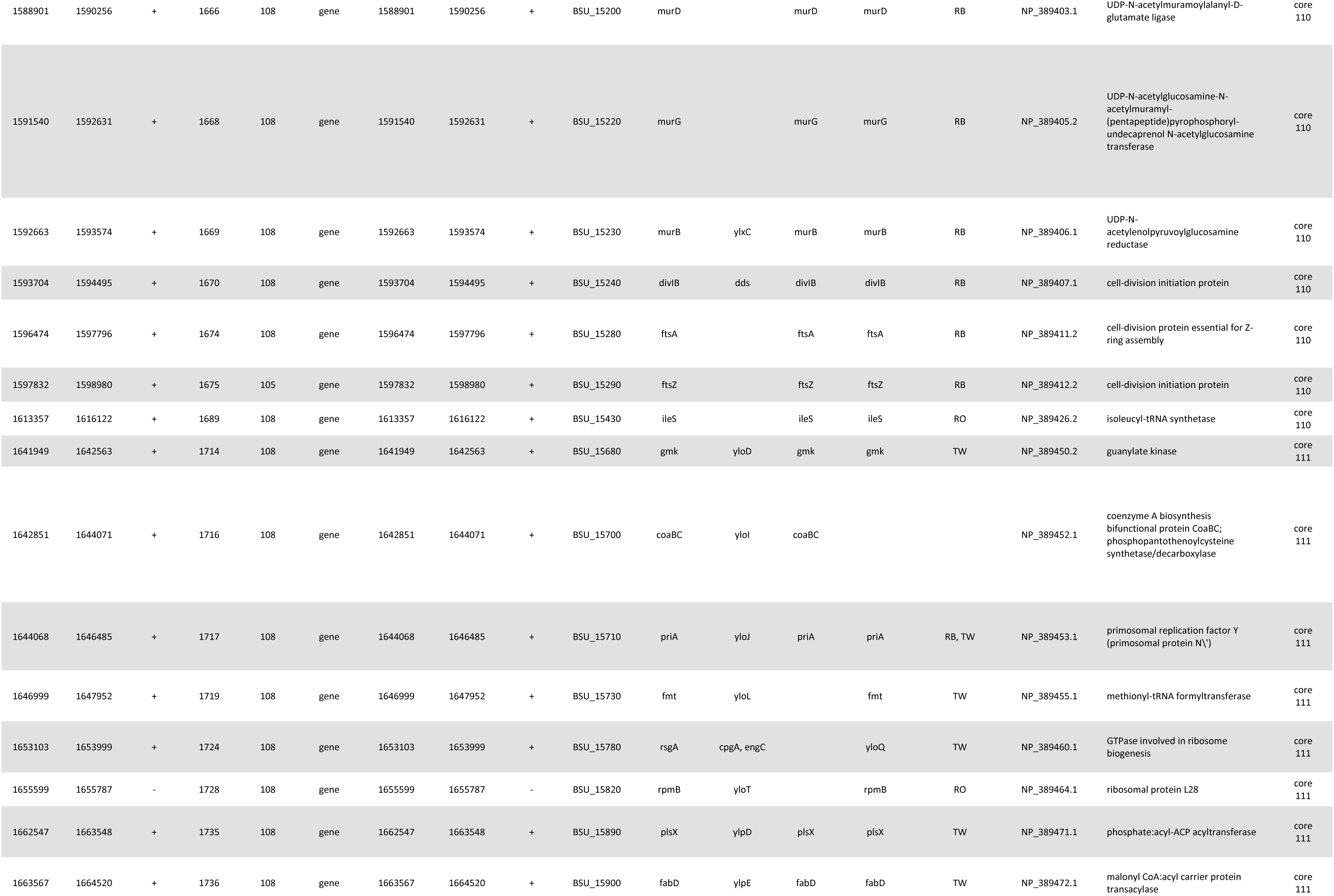
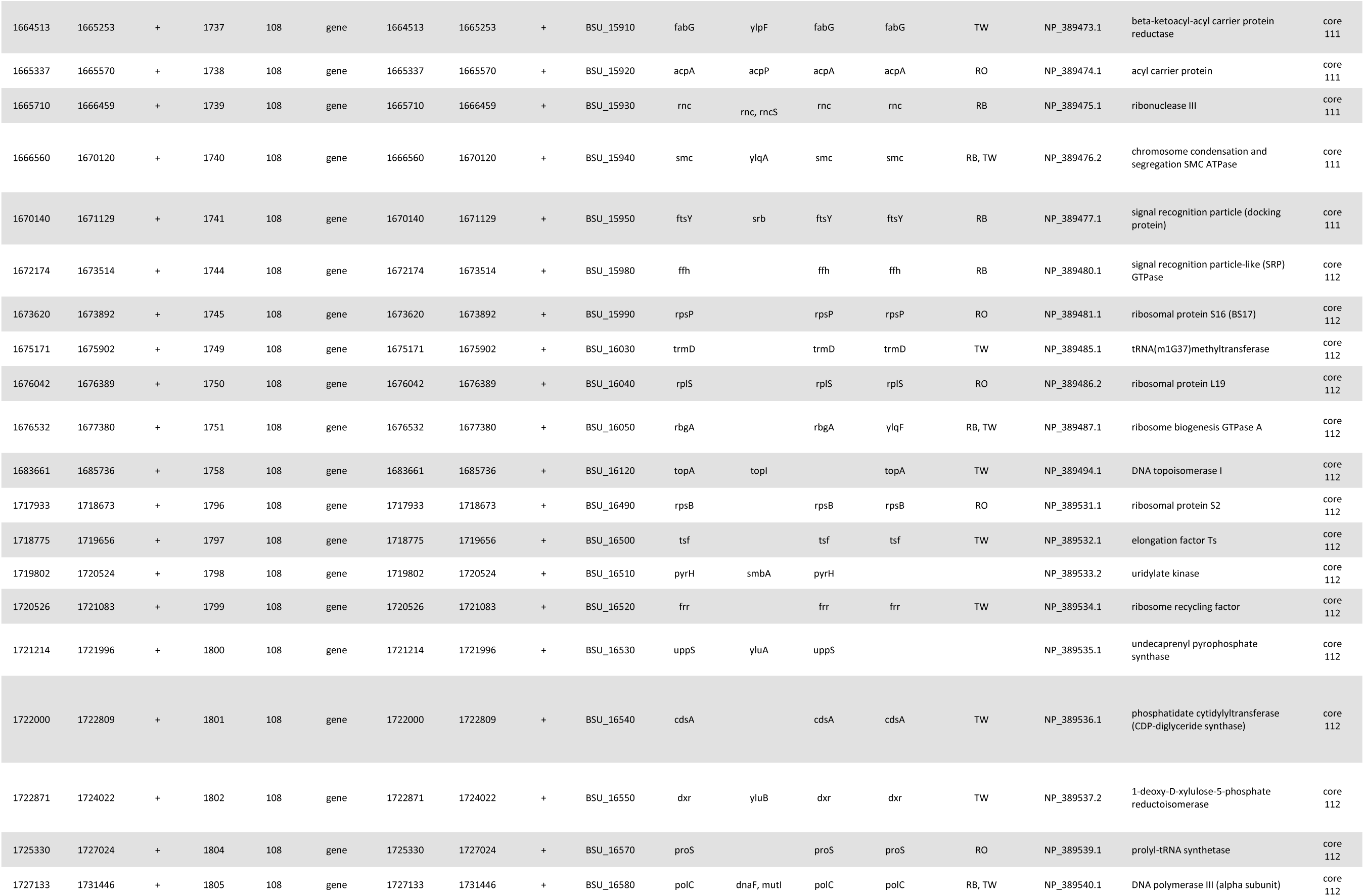
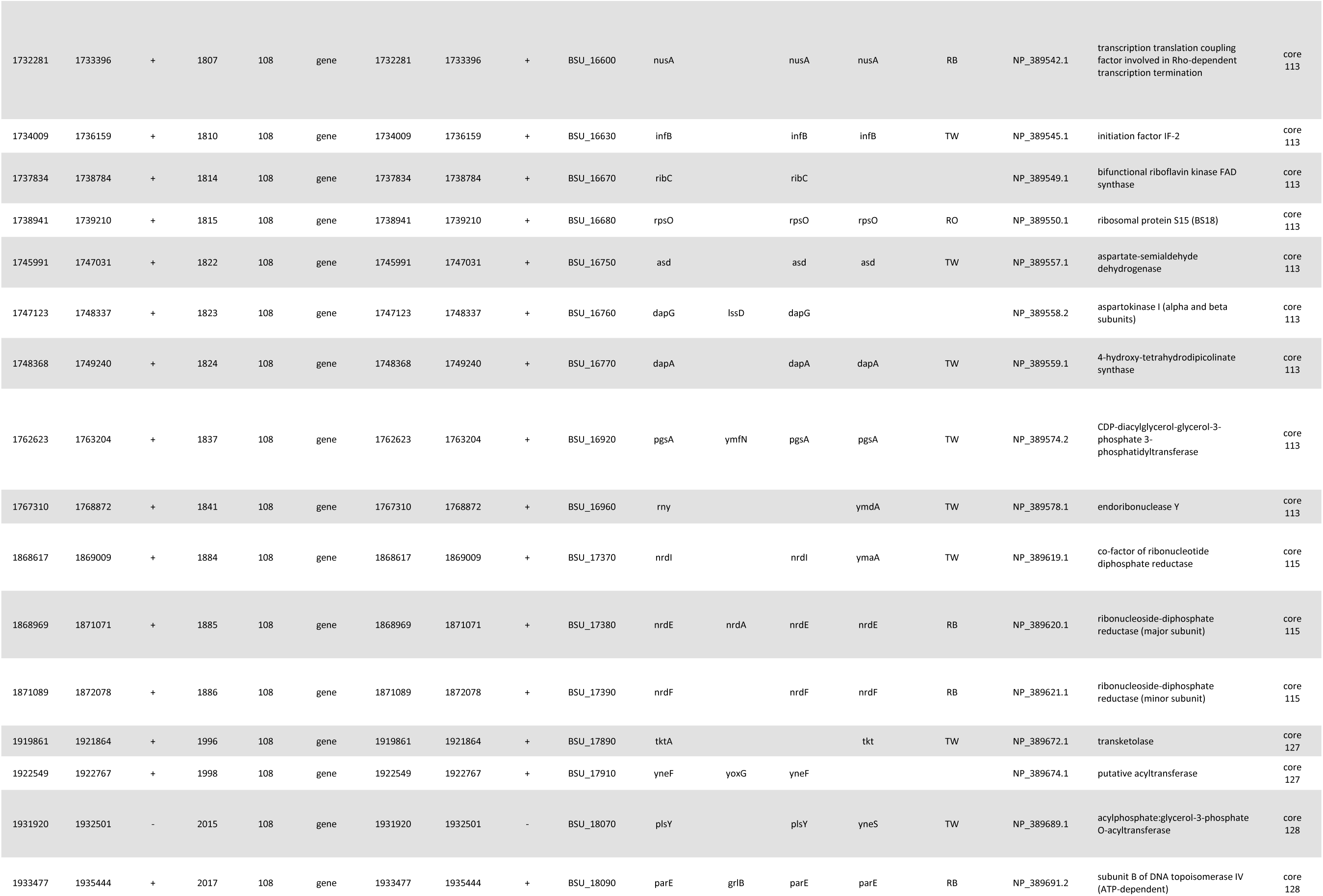
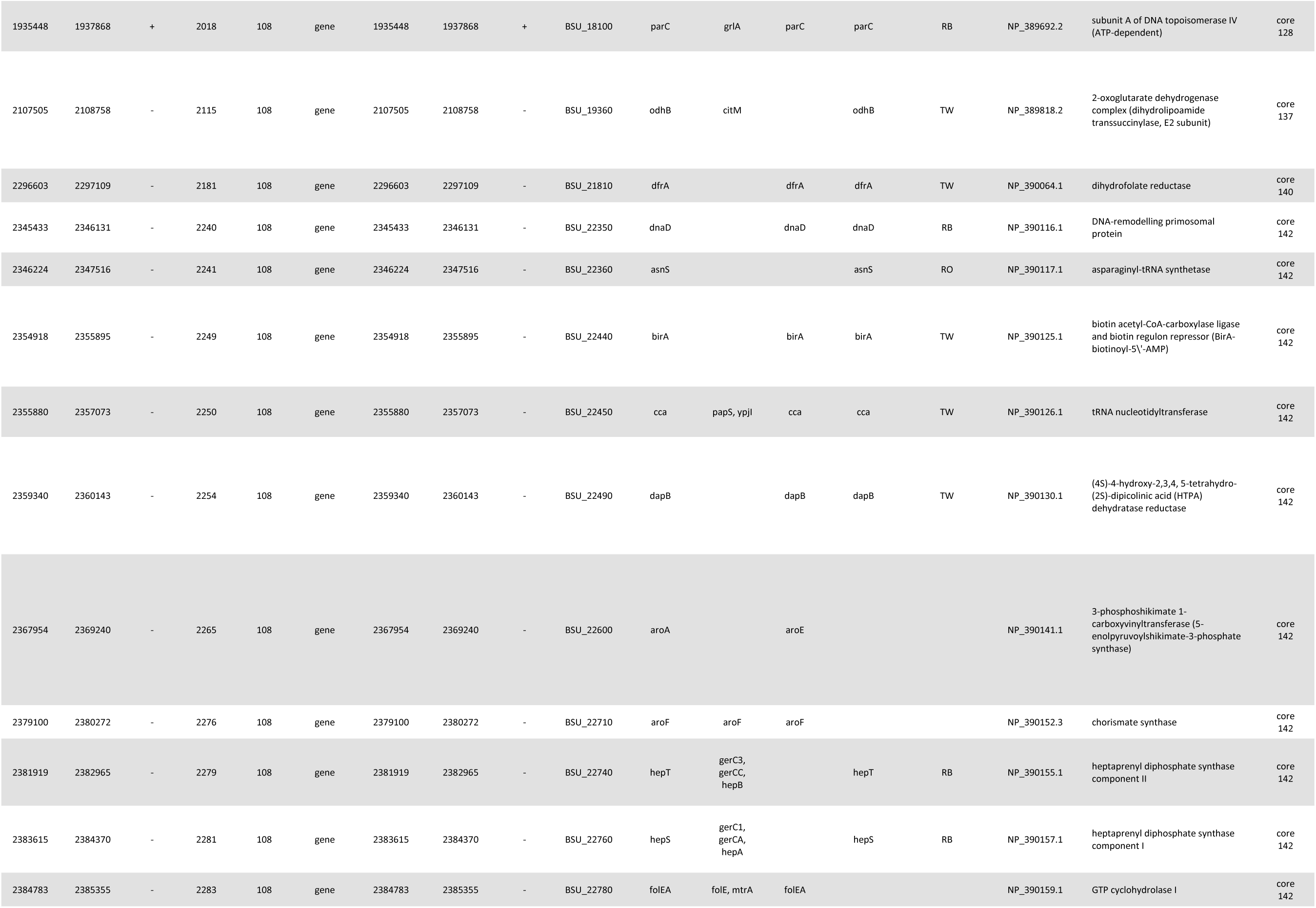
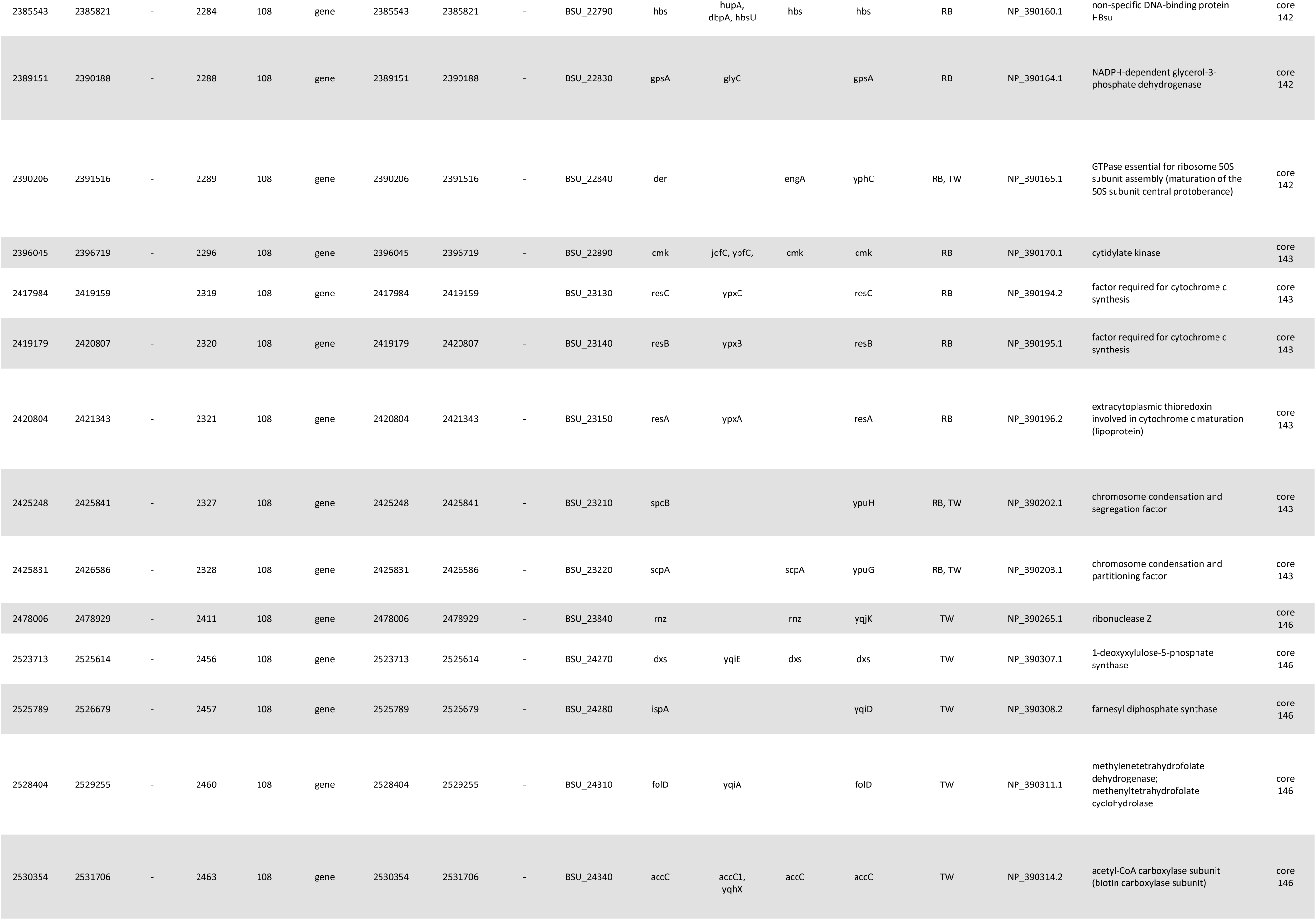
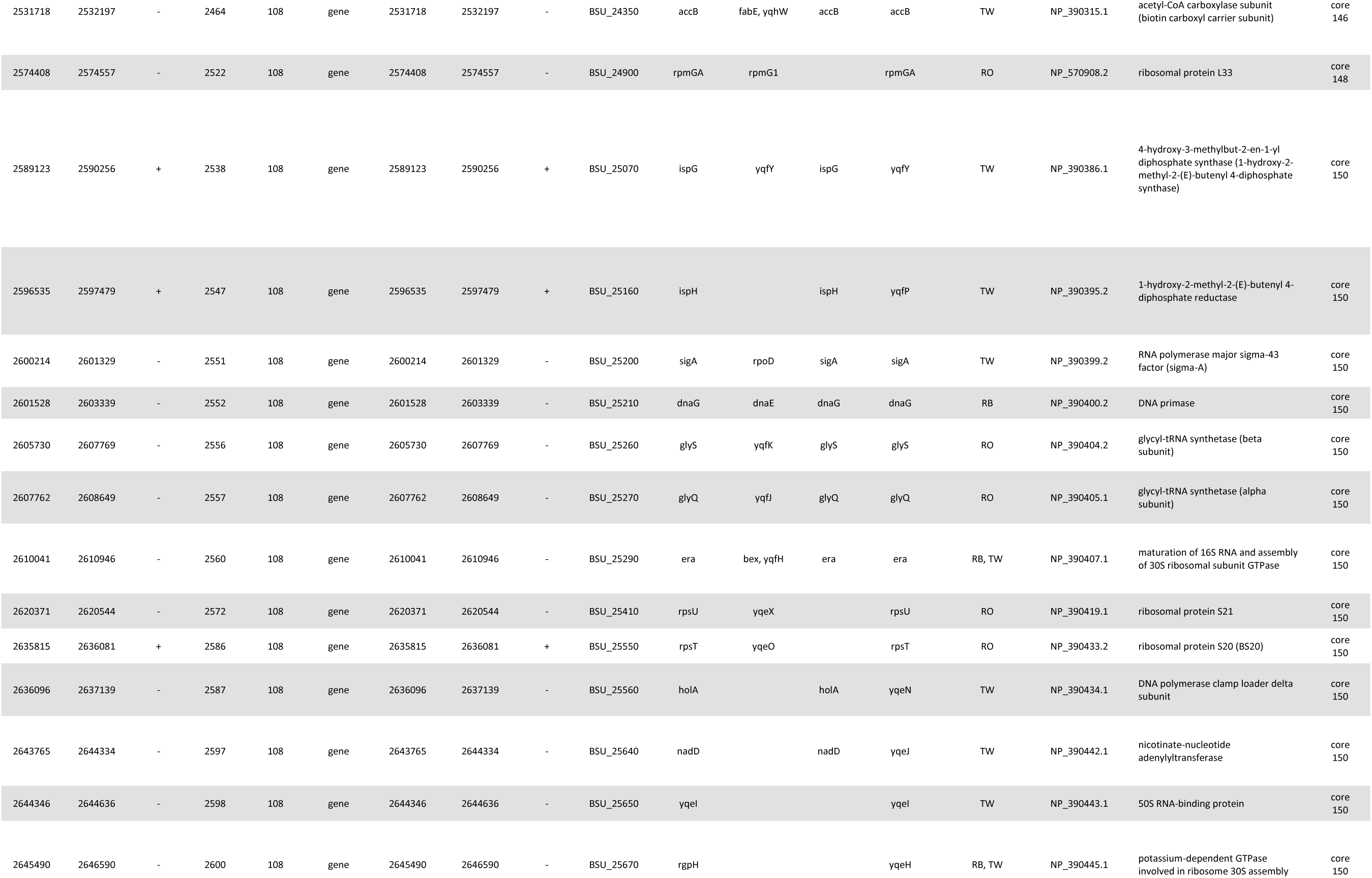
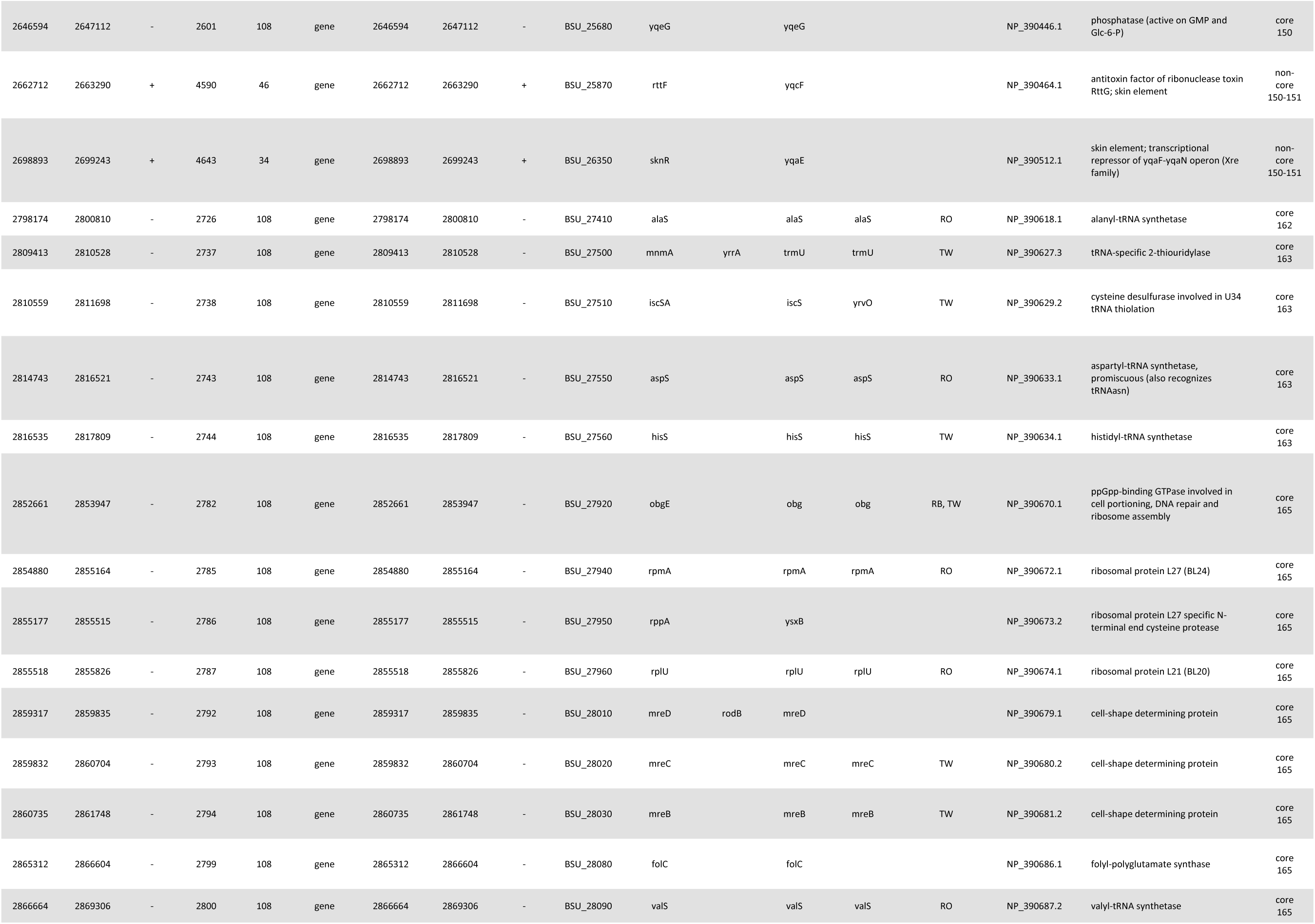
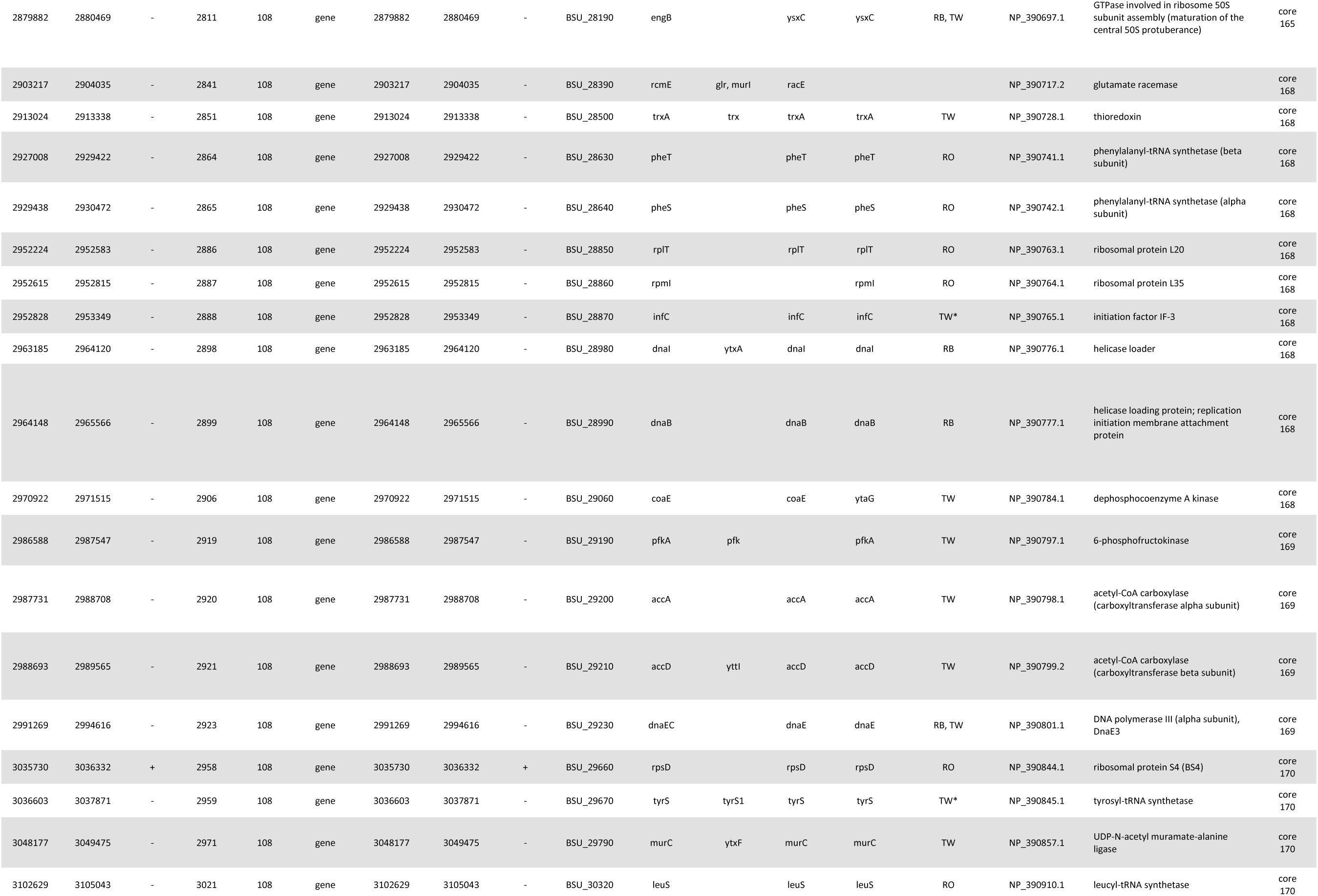
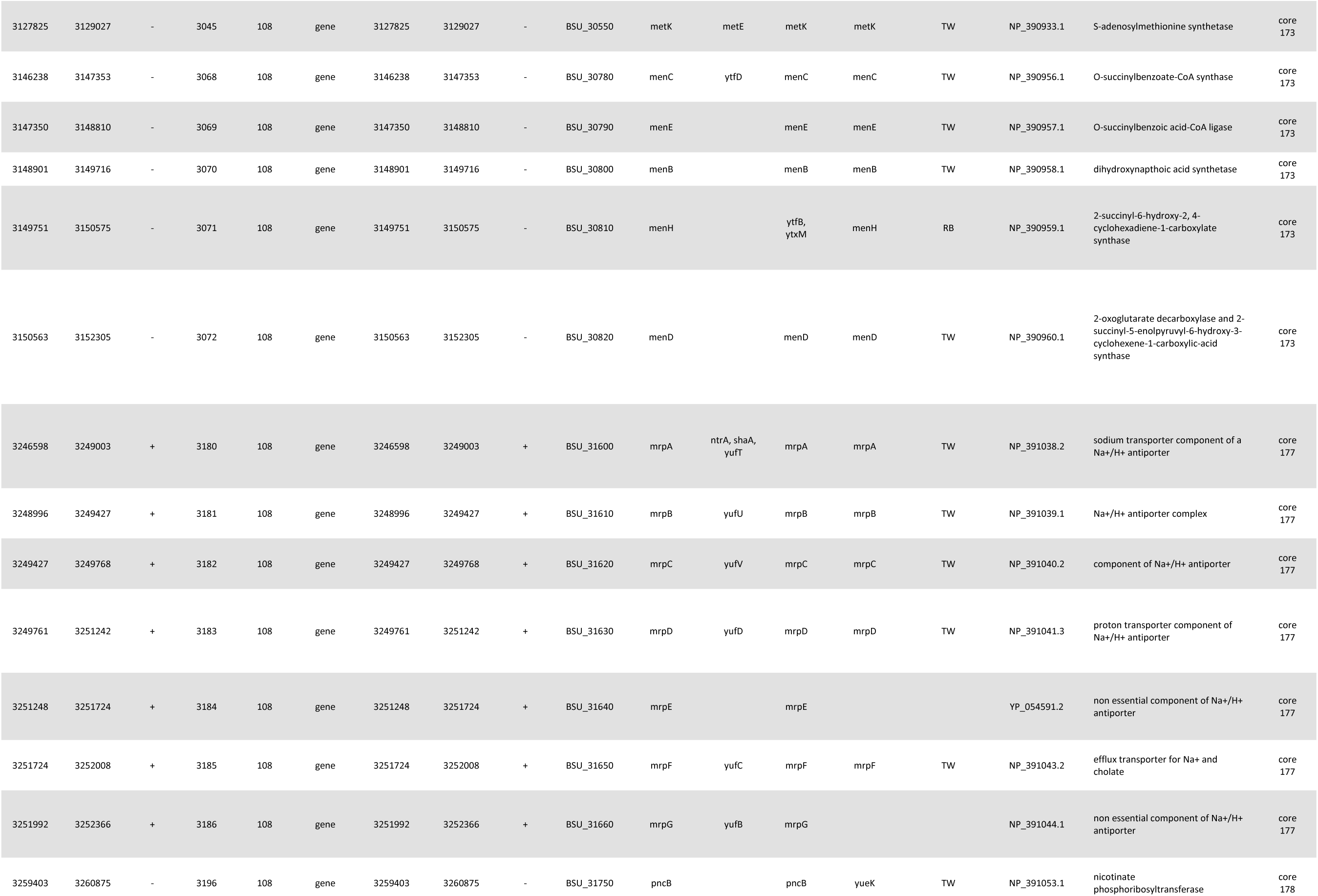
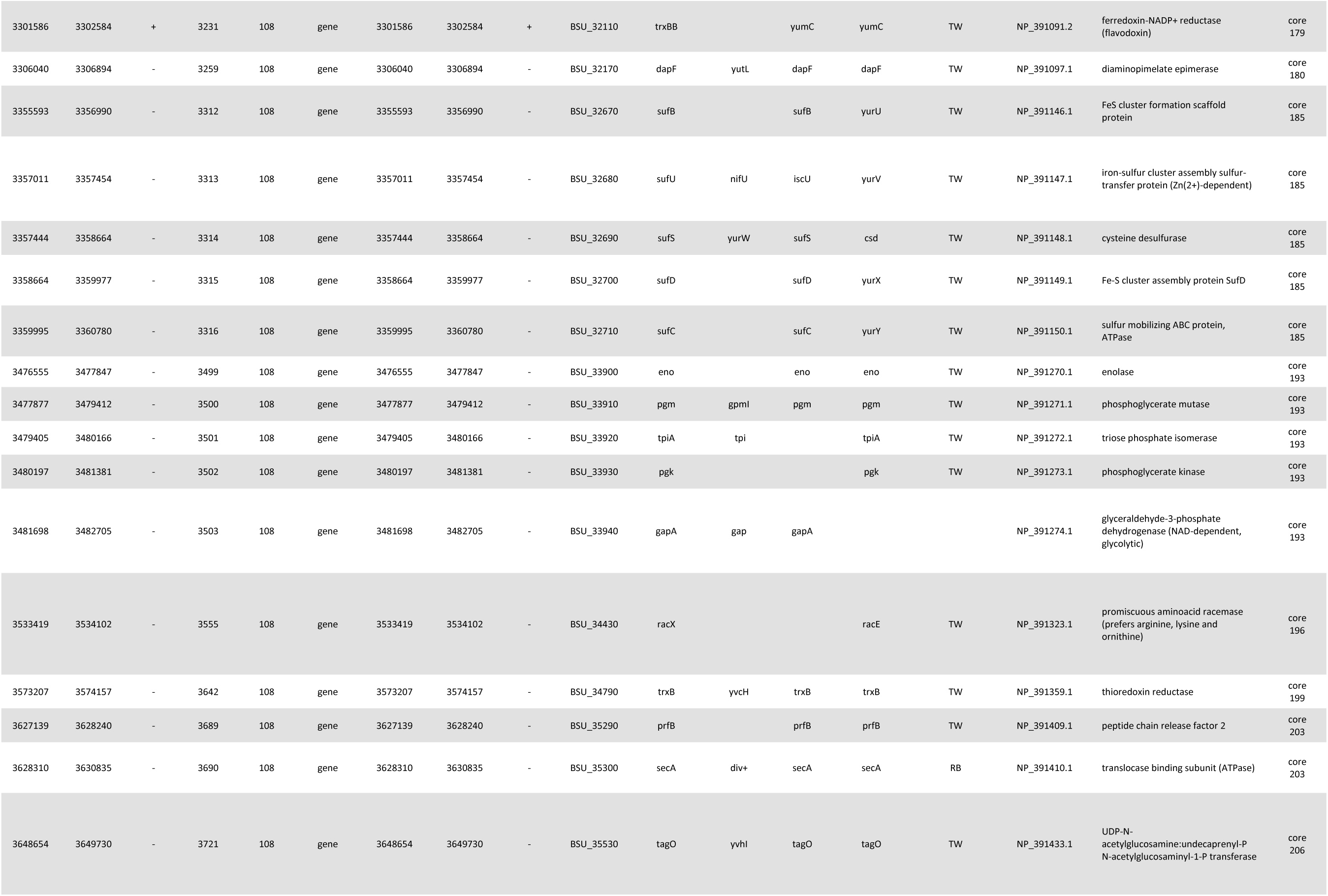
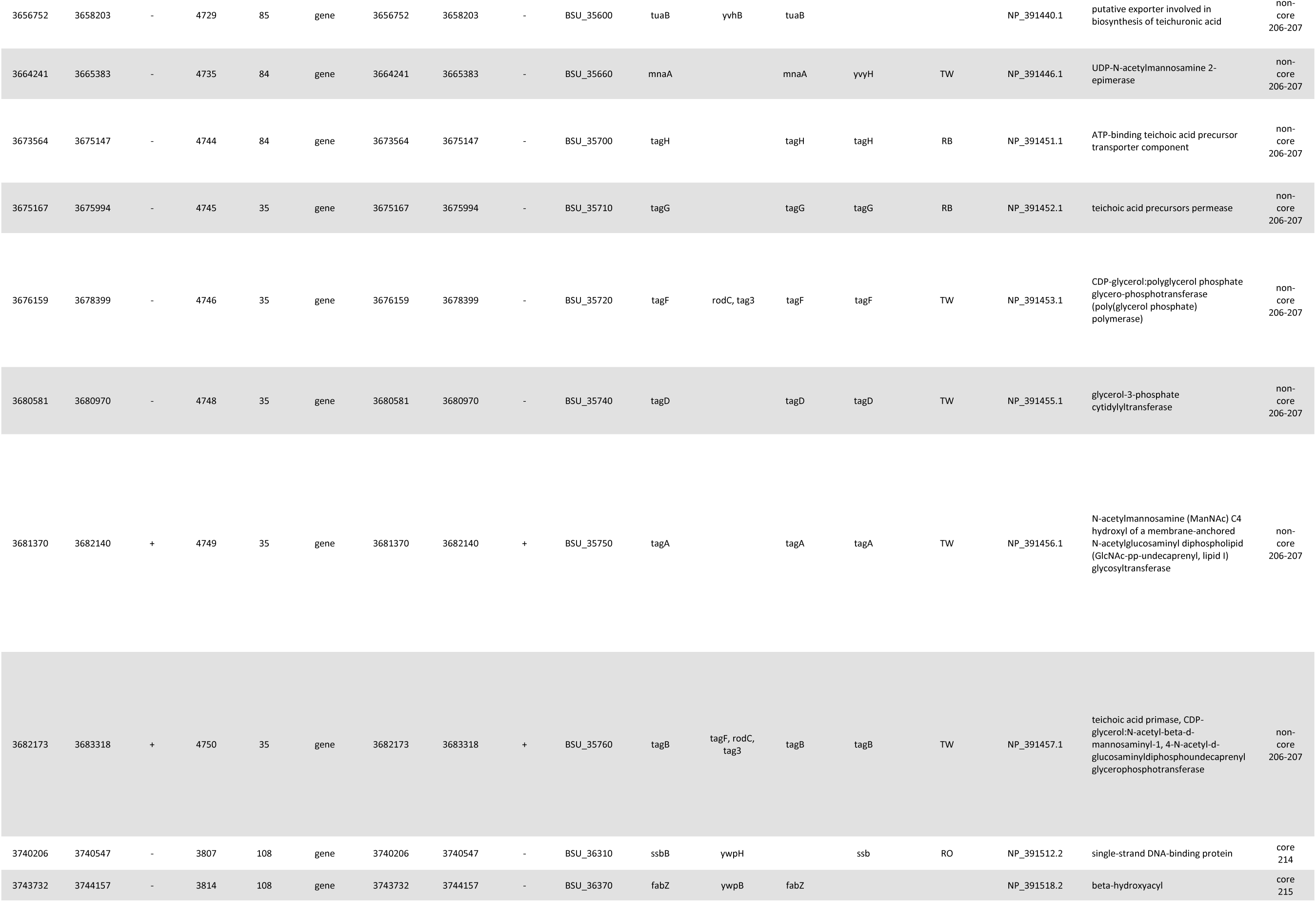
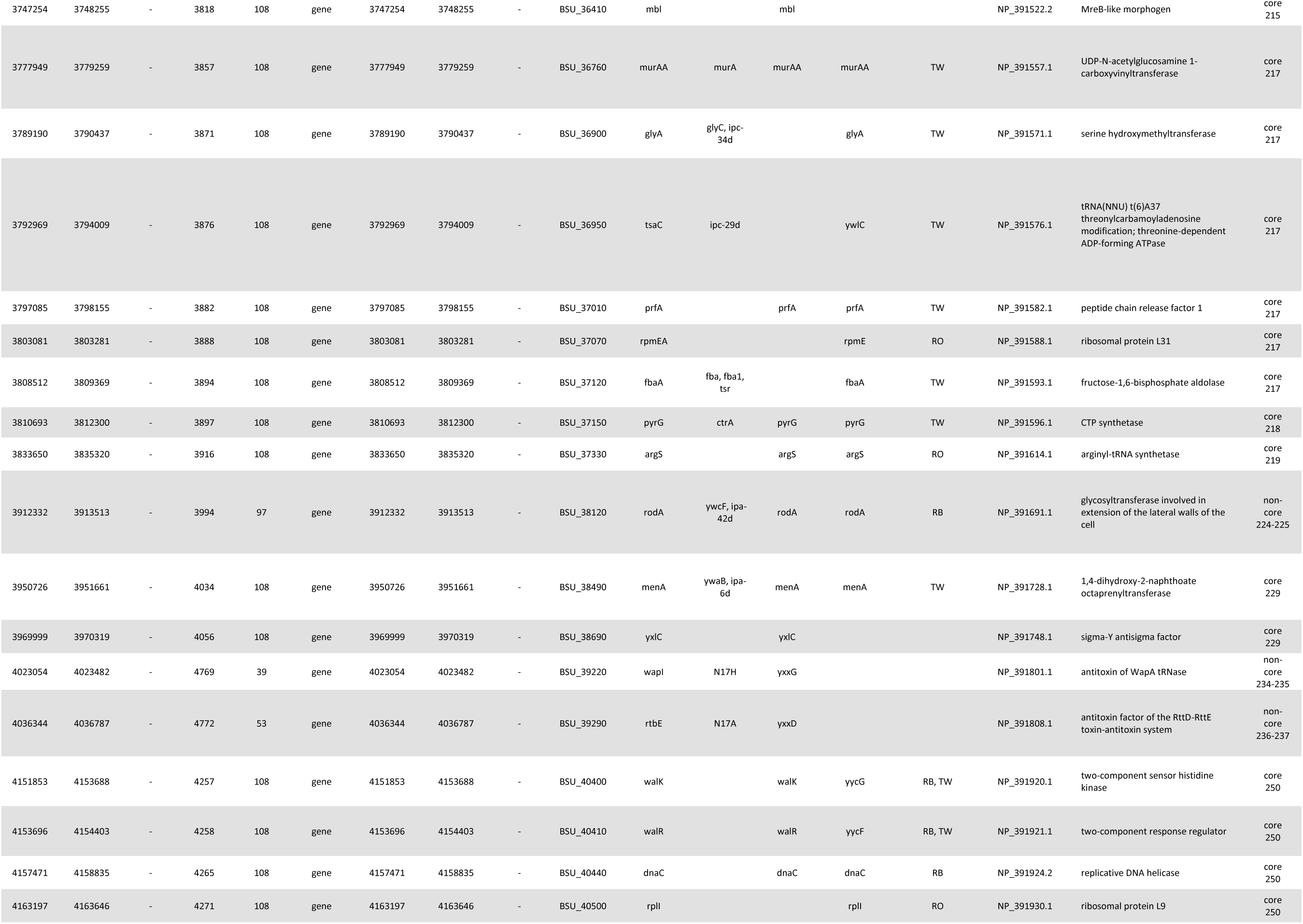
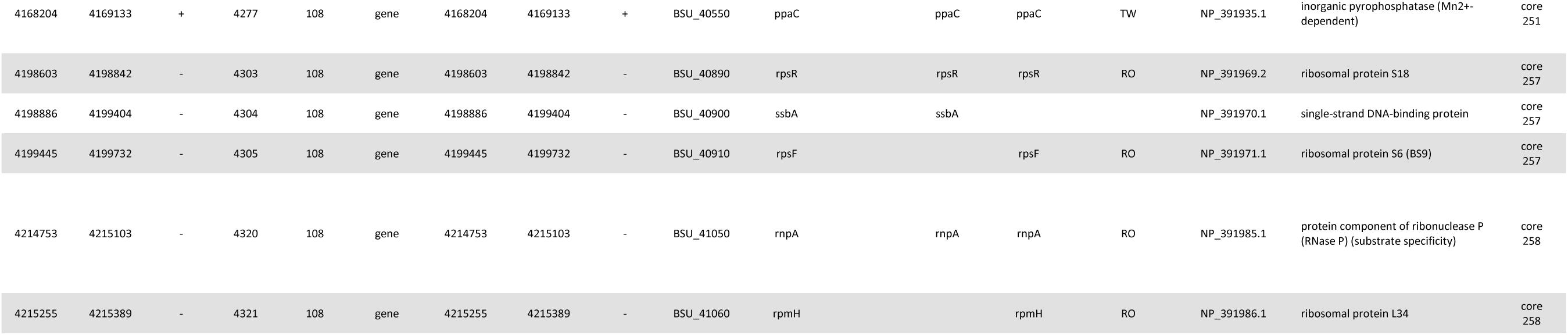
The 305 *B. subtilis* genes deemed essential by either Kobayashi et al. or Koo et al. These genes are compared to the PGG based annotation of core regions for the type strain genome used by Kobayashi et al. and Koo et al. (strain 168, GenBank sequence AL009126.3, BioSample SAMEA3138188, Assembly ASM904v1**/**GCF_000009045.1). Columns 1-5 are the start, stop, strand, cluster, and cluster size for the PGG annotation. Columns 6-11 are the gene type, start, stop, strand, locus tag, and gene symbol/name for the GenBank annotation. Column 12 is a list of synonyms for *B. subtilis* genes associated with the Koo and Kobayashi genes. Column 13 is the Koo et al. gene symbol/name. Columns 14-15 are the Kobayashi et al. gene symbol/name and evidence type (from Supporting Table 4 “RB, reference to study with *Bacillus subtilis*; RO, reference to study with other bacteria; TW, this work; TW*, inactivation failed but IPTG mutant could not be made”). Columns 16-17 are the GenBank protein product accession and name. Column 18 is the PGG core or non-core region the gene is contained in.

Another 8 genes are involved in wall teichoic acid (WTA) biosynthesis: Genes *tuaB* (cluster 4729 present in 85 of 108 genomes), *mnaA/yvyH* (cluster 4735 present in 84 of 108 genomes), *tagH* (cluster 4744 present in 84 of 108 genomes), *tagG* (cluster 4745 present in 35 of 108 genomes), *tagF* (cluster 4746 present in 35 of 108 genomes), *tagD* (cluster 4748 present in 35 of 108 genomes), *tagA* (cluster 4749 present in 35 of 108 genomes) and *tagB* (cluster 4750 present in 35 of 108 genomes). The WTA genes are involved in production of anionic glycopolymers required for consistent cell shape and division (13). The WTA genes are part of a 31 gene region which has been shown to be dispensable (14) but results in misformed cells with poor growth properties. Gene *rodA* (cluster 3994 present in 97 of 108 genomes) appears to be the exception as it is asserted to be essential for maintaining a rod shape and preventing spherical cells which lyse (15). Kobayashi et al. (2) stated: “Ten essential genes are involved in cell shape and division. Septum formation requires seven (ftsA, L, W, and Z, divIB and C, and pbpB; ref. 21), whereas cell shape requires three (rodA, and mreB and C).” Interestingly, genes *ftsZ* (cluster 1675 present in 105 of 108 genomes) and *pbpB* (cluster 1662 present in 107 of 108 genomes) while considered core, using our 95% of genomes definition are the only core genes not present in all 108 genomes. We investigated these 11 genes further to understand why essential genes did not appear to be core genes. By examining the PGG we discovered that alternate genes with homology to the essential genes had replaced the essential genes. Gene *pbpB* (cluster 1662 in 107 genomes) is replaced in the one remaining genome by cluster 7120 which is also annotated as *pbpB*. Gene *ftsZ* (cluster 1675 in 105 genomes) is replaced in three genones by a four gene insertion of clusters 8068, 8300, 8069, and 8070 where both 8068 and 8070 are annotated as *ftsZ*. Gene *rodA* (cluster 3994 in 97 genomes) is replaced by either: cluster 8718 (2 genomes) or clusters 10492, 6436, and 6437 (1 genome) or clusters 6436 and 6437 (8 genomes) where 8718 and 6436 are annotated as *rodA*. For *B. subtilis* spp. *spizizenii* strain W23, poly(ribitol phosphate) is the main teichoic acid (16) and this was thought to distinguish spp. *spizizenii* from spp. *subtilis* whose type strain 168 has poly(glycerol phosphate) as the main teichoic acid. Further study found that the ribitol/glycerol distinction does not distinguish between *spizizenii* and *subtilis* subspecies (17) but rather either subspecies can contain one or the other. Our PGG confirms this and in fact finds six distinct vairieties of the WTA region. For example the *tagD* gene (cluster 4748 in 35 genomes) has been replaced by mutlitple orthologs with the same annotation: cluster 3746 (23 genomes), cluster 5431 (43 genomes), cluster 6915 (2 genomes), cluster 7624 (3 genomes), and cluster 8731 (1 genome).

This shows that most essential genes in *B. subtilis* ssp. *subtilis* are encompassed by core genes/regions. There are 258 core regions for *B. subtilis* (Table 2). The 289 essential genes which are core genes are contained in only 63 of these regions. These 289 essential genes are not evenly distributed in these 63 regions (e.g., 46 are in core region 3). Similarly, the 16 essential genes in non-core regions (the regions between core regions) are contained in only 7 non-core regions with 8 WTA genes in the non-core region between core regions 206 and 207. A table of all *B. subtilis* genes is provided in Supplementary Table 1.

**Table 2.** The 258 *B. subtilis* core regions identified through the pan-genome graph.

| Core Region | Start | Stop | Length (bp) |
| --- | --- | --- | --- |
| 1 | 410 | 92089 | 91680 |
| 2 | 92253 | 95352 | 3100 |
| 3 | 98109 | 165706 | 67598 |
| 4 | 165830 | 165902 | 73 |
| 5 | 166253 | 166328 | 76 |
| 6 | 166501 | 168053 | 1553 |
| 7 | 177083 | 202079 | 24997 |
| 8 | 205409 | 210946 | 5538 |
| 9 | 222971 | 227954 | 4984 |
| 10 | 228066 | 238129 | 10064 |
| 11 | 241917 | 243661 | 1745 |
| 12 | 243892 | 250053 | 6162 |
| 13 | 252514 | 282897 | 30384 |
| 14 | 283003 | 317603 | 34601 |
| 15 | 317725 | 318927 | 1203 |
| 16 | 319180 | 320352 | 1173 |
| 17 | 320421 | 327377 | 6957 |
| 18 | 327618 | 332396 | 4779 |
| 19 | 335329 | 340585 | 5257 |
| 20 | 340613 | 364142 | 23530 |
| 21 | 365170 | 365754 | 585 |
| 22 | 365841 | 369217 | 3377 |
| 23 | 369773 | 370105 | 333 |
| 24 | 370259 | 406007 | 35749 |
| 25 | 406131 | 435989 | 29859 |
| 26 | 436036 | 455738 | 19703 |
| 27 | 455771 | 478741 | 22971 |
| 28 | 488314 | 490554 | 2241 |
| 29 | 490777 | 514764 | 23988 |
| 30 | 515016 | 524249 | 9234 |
| 31 | 525743 | 529419 | 3677 |
| 32 | 559264 | 560612 | 1349 |
| 33 | 561514 | 566101 | 4588 |
| 34 | 568345 | 569949 | 1605 |
| 35 | 570371 | 575562 | 5192 |
| 36 | 579541 | 582479 | 2939 |
| 37 | 582536 | 585037 | 2502 |
| 38 | 585155 | 587336 | 2182 |
| 39 | 587744 | 591891 | 4148 |
| 40 | 593407 | 599067 | 5661 |
| 41 | 599107 | 600105 | 999 |
| 42 | 601019 | 602427 | 1409 |
| 43 | 603012 | 604576 | 1565 |
| 44 | 604736 | 631811 | 27076 |
| 45 | 632774 | 648704 | 15931 |
| 46 | 648742 | 651868 | 3127 |
| 47 | 664775 | 680873 | 16099 |
| 48 | 680907 | 694281 | 13375 |
| 49 | 694425 | 696998 | 2574 |
| 50 | 697157 | 724825 | 27669 |
| 51 | 724987 | 737347 | 12361 |
| 52 | 737603 | 738982 | 1380 |
| 53 | 749562 | 749763 | 202 |
| 54 | 750959 | 753131 | 2173 |
| 55 | 753817 | 754401 | 585 |
| 56 | 755907 | 764765 | 8859 |
| 57 | 764781 | 770113 | 5333 |
| 58 | 772142 | 788636 | 16495 |
| 59 | 791462 | 795867 | 4406 |
| 60 | 796001 | 813818 | 17818 |
| 61 | 814384 | 817243 | 2860 |
| 62 | 827993 | 844645 | 16653 |
| 63 | 850367 | 860263 | 9897 |
| 64 | 860303 | 875055 | 14753 |
| 65 | 875178 | 878051 | 2874 |
| 66 | 879002 | 886173 | 7172 |
| 67 | 888143 | 893798 | 5656 |
| 68 | 893904 | 898867 | 4964 |
| 69 | 905010 | 909125 | 4116 |
| 70 | 909198 | 911928 | 2731 |
| 71 | 913924 | 928658 | 14735 |
| 72 | 928803 | 953294 | 24492 |
| 73 | 953373 | 957667 | 4295 |
| 74 | 959311 | 988377 | 29067 |
| 75 | 988417 | 988872 | 456 |
| 76 | 989712 | 1013851 | 24140 |
| 77 | 1013958 | 1050234 | 36277 |
| 78 | 1050443 | 1092728 | 42286 |
| 79 | 1092770 | 1108704 | 15935 |
| 80 | 1108736 | 1125153 | 16418 |
| 81 | 1125123 | 1135225 | 10103 |
| 82 | 1135255 | 1149935 | 14681 |
| 83 | 1150206 | 1156473 | 6268 |
| 84 | 1156504 | 1159081 | 2578 |
| 85 | 1160776 | 1169849 | 9074 |
| 86 | 1178218 | 1182395 | 4178 |
| 87 | 1187700 | 1204420 | 16721 |
| 88 | 1204506 | 1213449 | 8944 |
| 89 | 1217326 | 1250504 | 33179 |
| 90 | 1252177 | 1262861 | 10685 |
| 91 | 1268275 | 1268592 | 318 |
| 92 | 1270426 | 1276291 | 5866 |
| 93 | 1276337 | 1289101 | 12765 |
| 94 | 1290018 | 1313615 | 23598 |
| 95 | 1313840 | 1314304 | 465 |
| 96 | 1315869 | 1334247 | 18379 |
| 97 | 1339632 | 1343227 | 3596 |
| 98 | 1345631 | 1382430 | 36800 |
| 99 | 1382677 | 1383147 | 471 |
| 100 | 1383320 | 1417416 | 34097 |
| 101 | 1417561 | 1441788 | 24228 |
| 102 | 1447251 | 1466599 | 19349 |
| 103 | 1466638 | 1478836 | 12199 |
| 104 | 1482248 | 1494362 | 12115 |
| 105 | 1494403 | 1514963 | 20561 |
| 106 | 1514997 | 1534239 | 19243 |
| 107 | 1534279 | 1538692 | 4414 |
| 108 | 1538770 | 1557631 | 18862 |
| 109 | 1558034 | 1573790 | 15757 |
| 110 | 1575051 | 1616122 | 41072 |
| 111 | 1616267 | 1671129 | 54863 |
| 112 | 1671166 | 1731446 | 60281 |
| 113 | 1731457 | 1780220 | 48764 |
| 114 | 1780404 | 1782523 | 2120 |
| 115 | 1860014 | 1879759 | 19746 |
| 116 | 1879787 | 1880377 | 591 |
| 117 | 1887352 | 1888743 | 1392 |
| 118 | 1888774 | 1895165 | 6392 |
| 119 | 1896424 | 1899125 | 2702 |
| 120 | 1899589 | 1900514 | 926 |
| 121 | 1901117 | 1901377 | 261 |
| 122 | 1901429 | 1903058 | 1630 |
| 123 | 1903511 | 1904749 | 1239 |
| 124 | 1905370 | 1906207 | 838 |
| 125 | 1906272 | 1906706 | 435 |
| 126 | 1907013 | 1907201 | 189 |
| 127 | 1910167 | 1924962 | 14796 |
| 128 | 1924993 | 1942455 | 17463 |
| 129 | 1942714 | 1957360 | 14647 |
| 130 | 1958027 | 1960087 | 2061 |
| 131 | 1998340 | 2003182 | 4843 |
| 132 | 2003276 | 2015681 | 12406 |
| 133 | 2017886 | 2021144 | 3259 |
| 134 | 2021223 | 2022467 | 1245 |
| 135 | 2073658 | 2075779 | 2122 |
| 136 | 2079096 | 2085980 | 6885 |
| 137 | 2086070 | 2128421 | 42352 |
| 138 | 2128464 | 2139457 | 10994 |
| 139 | 2140898 | 2151256 | 10359 |
| 140 | 2287097 | 2297109 | 10013 |
| 141 | 2297984 | 2330022 | 32039 |
| 142 | 2330075 | 2393559 | 63485 |
| 143 | 2393602 | 2433522 | 39921 |
| 144 | 2435012 | 2435224 | 213 |
| 145 | 2435833 | 2476859 | 41027 |
| 146 | 2476969 | 2532197 | 55229 |
| 147 | 2532353 | 2564883 | 32531 |
| 148 | 2564923 | 2581617 | 16695 |
| 149 | 2584035 | 2587573 | 3539 |
| 150 | 2588701 | 2653340 | 64640 |
| 151 | 2701362 | 2701754 | 393 |
| 152 | 2716035 | 2718802 | 2768 |
| 153 | 2718959 | 2724827 | 5869 |
| 154 | 2727160 | 2730226 | 3067 |
| 155 | 2732747 | 2732881 | 135 |
| 156 | 2732980 | 2739105 | 6126 |
| 157 | 2739486 | 2740193 | 708 |
| 158 | 2741357 | 2743889 | 2533 |
| 159 | 2747984 | 2751651 | 3668 |
| 160 | 2752167 | 2773222 | 21056 |
| 161 | 2773783 | 2784652 | 10870 |
| 162 | 2784688 | 2805313 | 20626 |
| 163 | 2805348 | 2832367 | 27020 |
| 164 | 2832424 | 2840925 | 8502 |
| 165 | 2841307 | 2897432 | 56126 |
| 166 | 2898292 | 2898929 | 638 |
| 167 | 2898931 | 2899752 | 822 |
| 168 | 2899816 | 2982269 | 82454 |
| 169 | 2982603 | 2998620 | 16018 |
| 170 | 3009915 | 3106995 | 97081 |
| 171 | 3107232 | 3109360 | 2129 |
| 172 | 3109397 | 3115974 | 6578 |
| 173 | 3117976 | 3183195 | 65220 |
| 174 | 3183201 | 3188048 | 4848 |
| 175 | 3188173 | 3208154 | 19982 |
| 176 | 3210445 | 3213829 | 3385 |
| 177 | 3214372 | 3253448 | 39077 |
| 178 | 3257092 | 3266655 | 9564 |
| 179 | 3269914 | 3305261 | 35348 |
| 180 | 3305599 | 3308231 | 2633 |
| 181 | 3308368 | 3318002 | 9635 |
| 182 | 3318029 | 3334990 | 16962 |
| 183 | 3335414 | 3344979 | 9566 |
| 184 | 3345013 | 3354487 | 9475 |
| 185 | 3355593 | 3382499 | 26907 |
| 186 | 3382633 | 3419421 | 36789 |
| 187 | 3419656 | 3436651 | 16996 |
| 188 | 3436849 | 3440961 | 4113 |
| 189 | 3441121 | 3450788 | 9668 |
| 190 | 3450709 | 3457941 | 7233 |
| 191 | 3459848 | 3464067 | 4220 |
| 192 | 3467564 | 3474070 | 6507 |
| 193 | 3476555 | 3488042 | 11488 |
| 194 | 3487974 | 3495701 | 7728 |
| 195 | 3495876 | 3499257 | 3382 |
| 196 | 3499541 | 3535473 | 35933 |
| 197 | 3536012 | 3542691 | 6680 |
| 198 | 3545889 | 3568135 | 22247 |
| 199 | 3568527 | 3580038 | 11512 |
| 200 | 3582936 | 3591268 | 8333 |
| 201 | 3591288 | 3601909 | 10622 |
| 202 | 3606762 | 3609221 | 2460 |
| 203 | 3610064 | 3630835 | 20772 |
| 204 | 3631003 | 3634754 | 3752 |
| 205 | 3636046 | 3647406 | 11361 |
| 206 | 3648654 | 3651068 | 2415 |
| 207 | 3684826 | 3688547 | 3722 |
| 208 | 3691372 | 3693906 | 2535 |
| 209 | 3695363 | 3696223 | 861 |
| 210 | 3696257 | 3714679 | 18423 |
| 211 | 3716009 | 3722541 | 6533 |
| 212 | 3724420 | 3725136 | 717 |
| 213 | 3727427 | 3727687 | 261 |
| 214 | 3728511 | 3741592 | 13082 |
| 215 | 3742351 | 3750677 | 8327 |
| 216 | 3750768 | 3761700 | 10933 |
| 217 | 3761987 | 3810611 | 48625 |
| 218 | 3810693 | 3819178 | 8486 |
| 219 | 3833650 | 3845998 | 12349 |
| 220 | 3847853 | 3851893 | 4041 |
| 221 | 3852186 | 3856447 | 4262 |
| 222 | 3856639 | 3874180 | 17542 |
| 223 | 3874332 | 3899182 | 24851 |
| 224 | 3900963 | 3912226 | 11264 |
| 225 | 3914009 | 3926399 | 12391 |
| 226 | 3926682 | 3937515 | 10834 |
| 227 | 3938310 | 3949922 | 11613 |
| 228 | 3949952 | 3950584 | 633 |
| 229 | 3950726 | 3973278 | 22553 |
| 230 | 3973364 | 3982915 | 9552 |
| 231 | 3983187 | 3984026 | 840 |
| 232 | 3984132 | 3989084 | 4953 |
| 233 | 3989948 | 3990967 | 1020 |
| 234 | 3992671 | 4018656 | 25986 |
| 235 | 4030710 | 4031645 | 936 |
| 236 | 4031797 | 4033755 | 1959 |
| 237 | 4038513 | 4039159 | 647 |
| 238 | 4039466 | 4040875 | 1410 |
| 239 | 4049009 | 4062143 | 13135 |
| 240 | 4062543 | 4084383 | 21841 |
| 241 | 4084799 | 4086540 | 1742 |
| 242 | 4088002 | 4092666 | 4665 |
| 243 | 4106245 | 4121056 | 14812 |
| 244 | 4121166 | 4123903 | 2738 |
| 245 | 4128119 | 4130044 | 1926 |
| 246 | 4134436 | 4139537 | 5102 |
| 247 | 4140260 | 4141474 | 1215 |
| 248 | 4141711 | 4145506 | 3796 |
| 249 | 4145747 | 4147132 | 1386 |
| 250 | 4147419 | 4165622 | 18204 |
| 251 | 4166815 | 4171359 | 4545 |
| 252 | 4176900 | 4178145 | 1246 |
| 253 | 4183445 | 4185681 | 2237 |
| 254 | 4186092 | 4187174 | 1083 |
| 255 | 4191198 | 4195744 | 4547 |
| 256 | 4196350 | 4196730 | 381 |
| 257 | 4197780 | 4205517 | 7738 |
| 258 | 4205556 | 4215389 | 9834 |

For *E. coli*, Goodall et al. (4) determined *E. coli* essential genes using an analysis of transposon insertion events (TraDIS). The results of their study and two other studies, the Keio collection (5) and the Profiling of the *E. coli* Chromosome (PEC) (6) were captured in Table S2 of Goodall et al. Of the 414 genes with overlap between these studies, the 248 essential genes in common for all three studies are all core genes (Table 3). This set of 248 essential genes should be the highest quality predictions as determined by all three studies and confirms our assertion that essential genes should almost always be core genes. The next highest quality set of essential gene predictions is the 45 essential genes where two of the three studies agree which 41 are core genes: for Keio-PEC 15 of 16 are core genes, for TraDIS-Keio 8 of 11 are core genes, and for TraDIS-PEC 18 of 18 are core genes (Table 3). The lowest quality set of essential gene predictions is the 121 essential genes where only one study agrees which 89 are core genes: for Keio only 12 of 22 are core genes, for PEC only 18 of 18 are core genes, and for TraDIS only 59 of 81 are core genes (Table 3). One of the noncore essential genes present in two studies (TraDIS-Keio), *racR,* is probably a toxin suppressor which is not essential in the absence of the toxins. Bindal et al. (18) noted, “We further show that both YdaS and YdaT can act independently as toxins and that RacR serves to counteract the toxicity by tightly downregulating the expression of these toxins.” The *racR* gene is found in only 106 of the 971 genomes in the *E. coli* PGG, whereas ydaS and ydaT are found in 106 and 150 genomes respectively, perhaps arguing that *ydaS* is the key toxin gene. This recapitulates the pattern we observed in *B. subtilis* where toxin suppressor genes are only essential in the presence of toxin genes. Similarly the *dicA* gene (TraDIS-Keio) can be deleted if the *dicB* gene is also deleted. Kato et al. (19) noted: “The *dicA* gene encoding a repressor of a cell division inhibitor was deleted in our study with the *dicB*, the inhibitor gene.” There are 521 core regions for *E. coli* (Table 4). The 378 essential genes which are core genes are contained in only 133 of these regions. These 378 essential genes are not evenly distributed in these 133 regions (e.g., 27 are in core region 362). Similarly, the 36 essential genes in non-core regions (the regions between core regions) are contained in only 23 non-core regions with 4 in the non-core region between core regions 152 and 153. A table of all *E. coli* genes is provided in Supplementary Table 3.

**Table 3.**
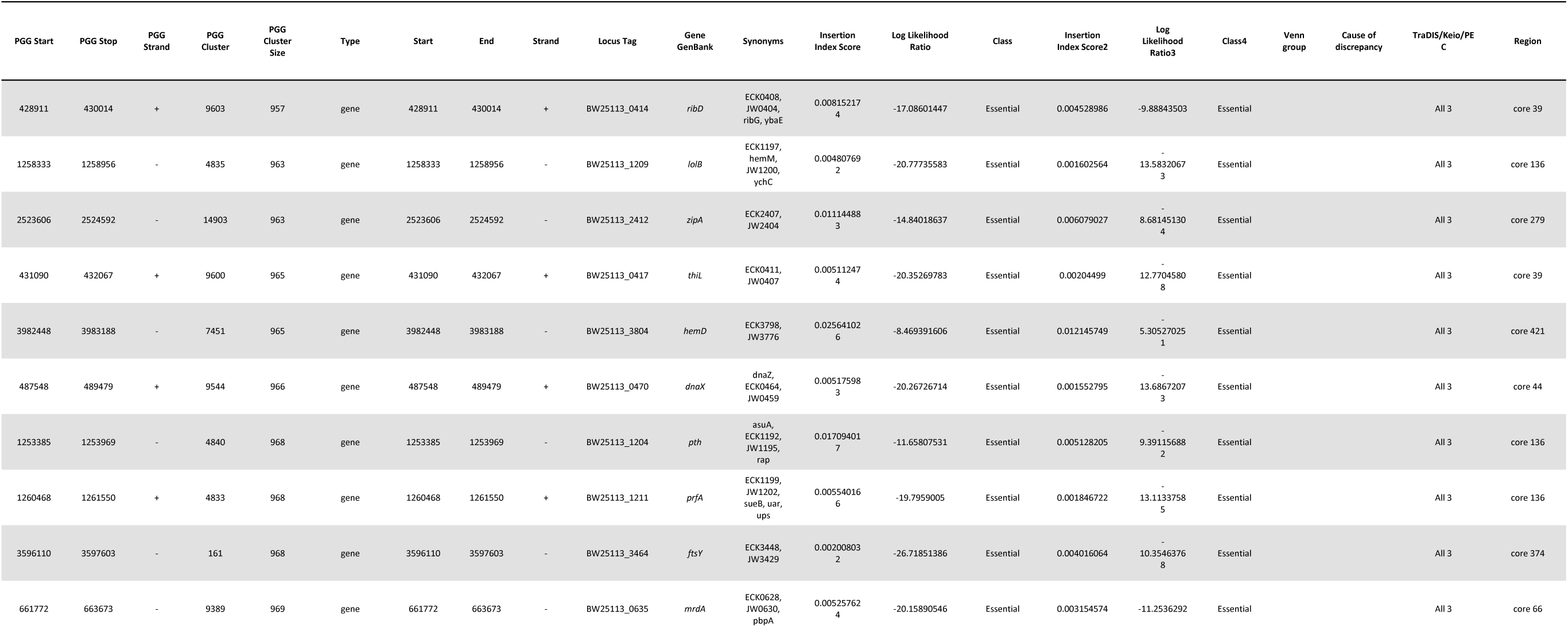
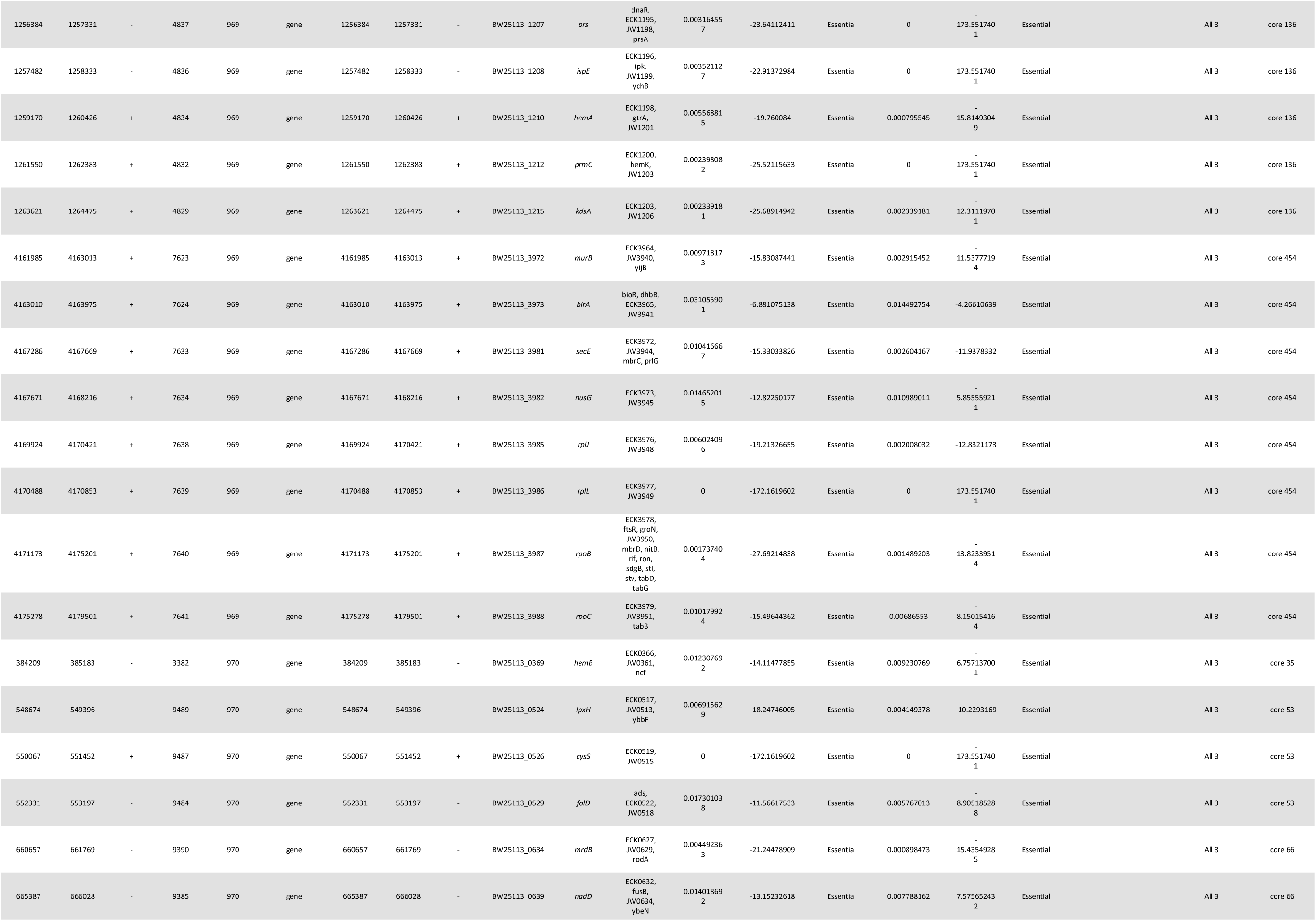
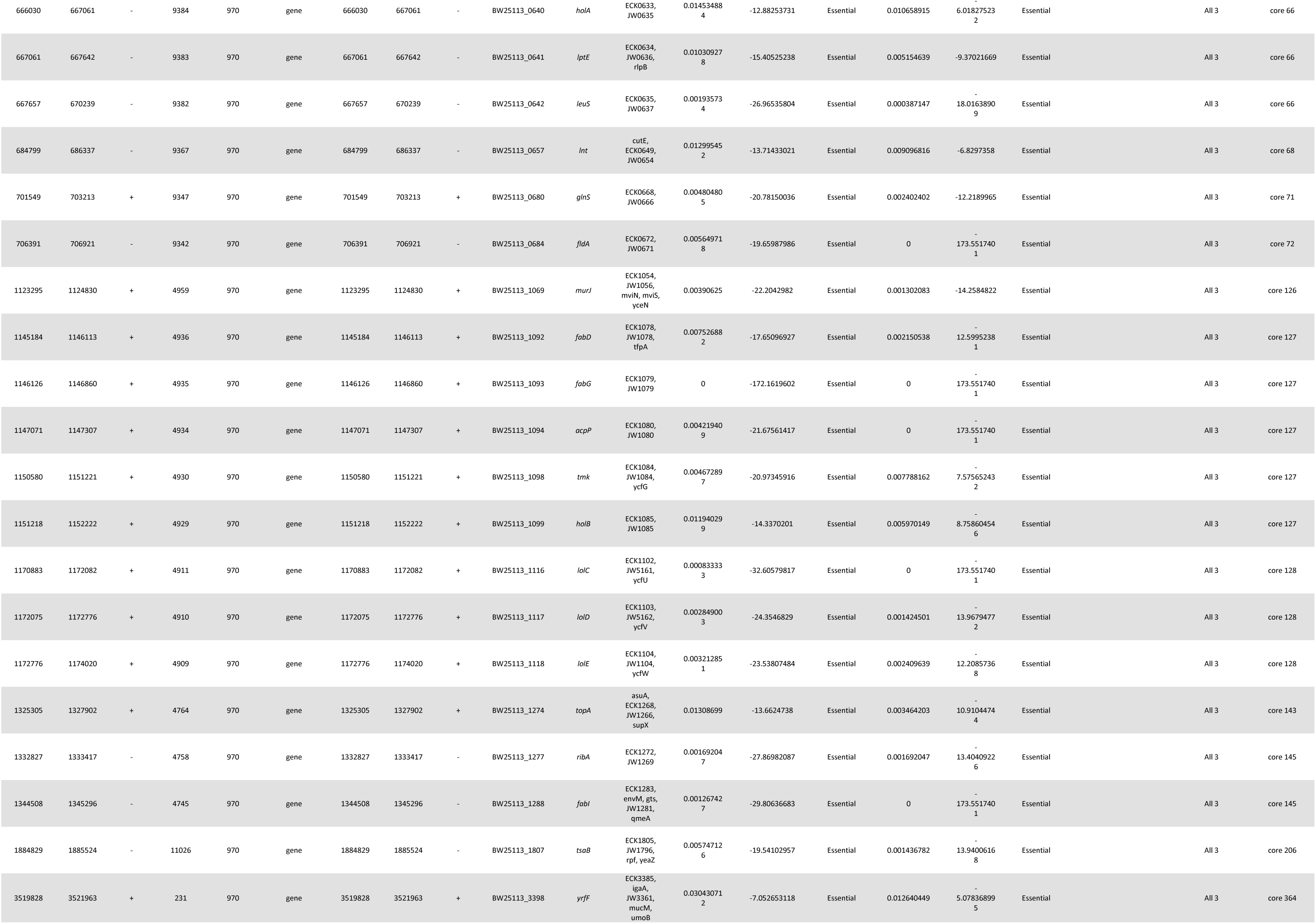
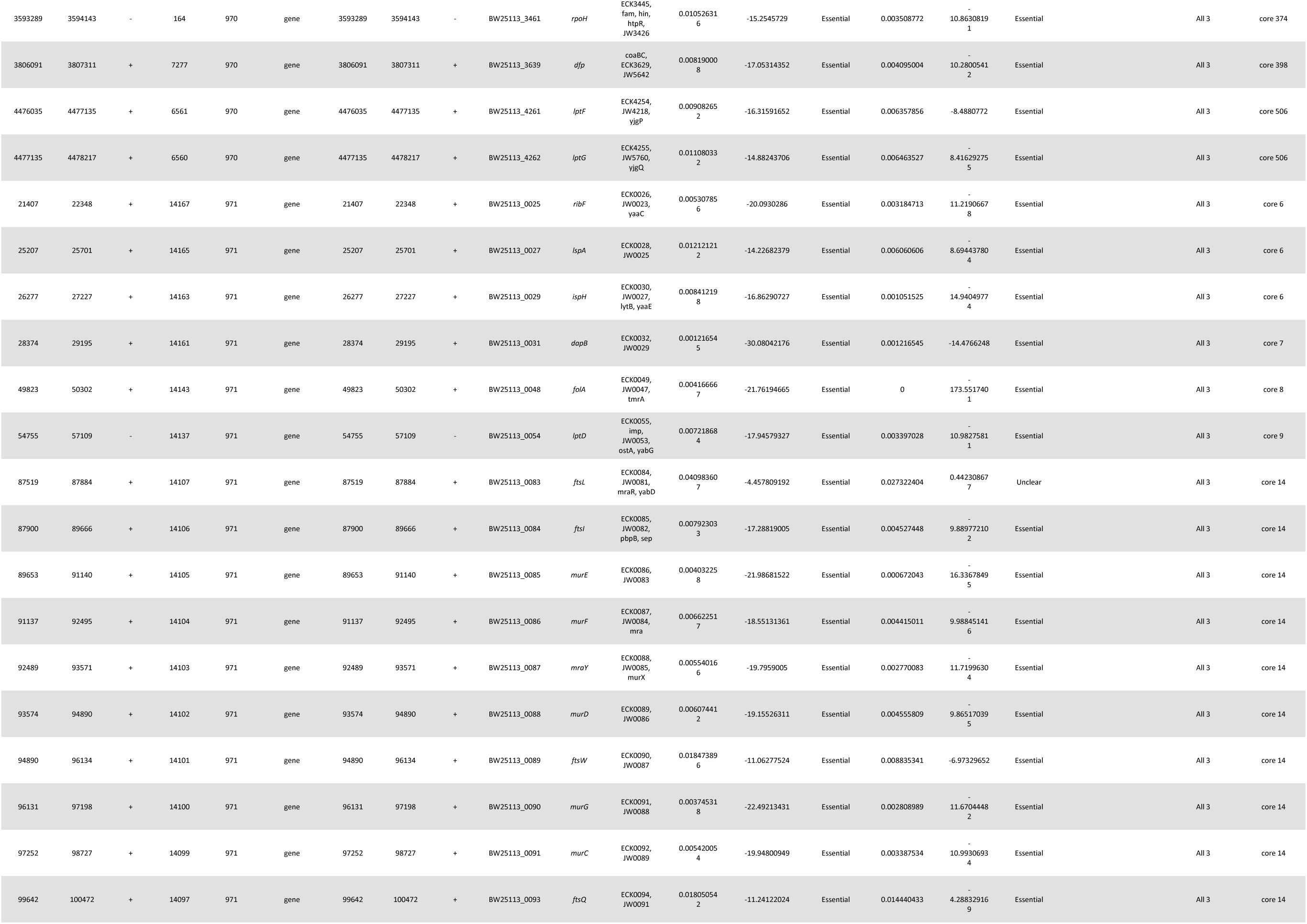
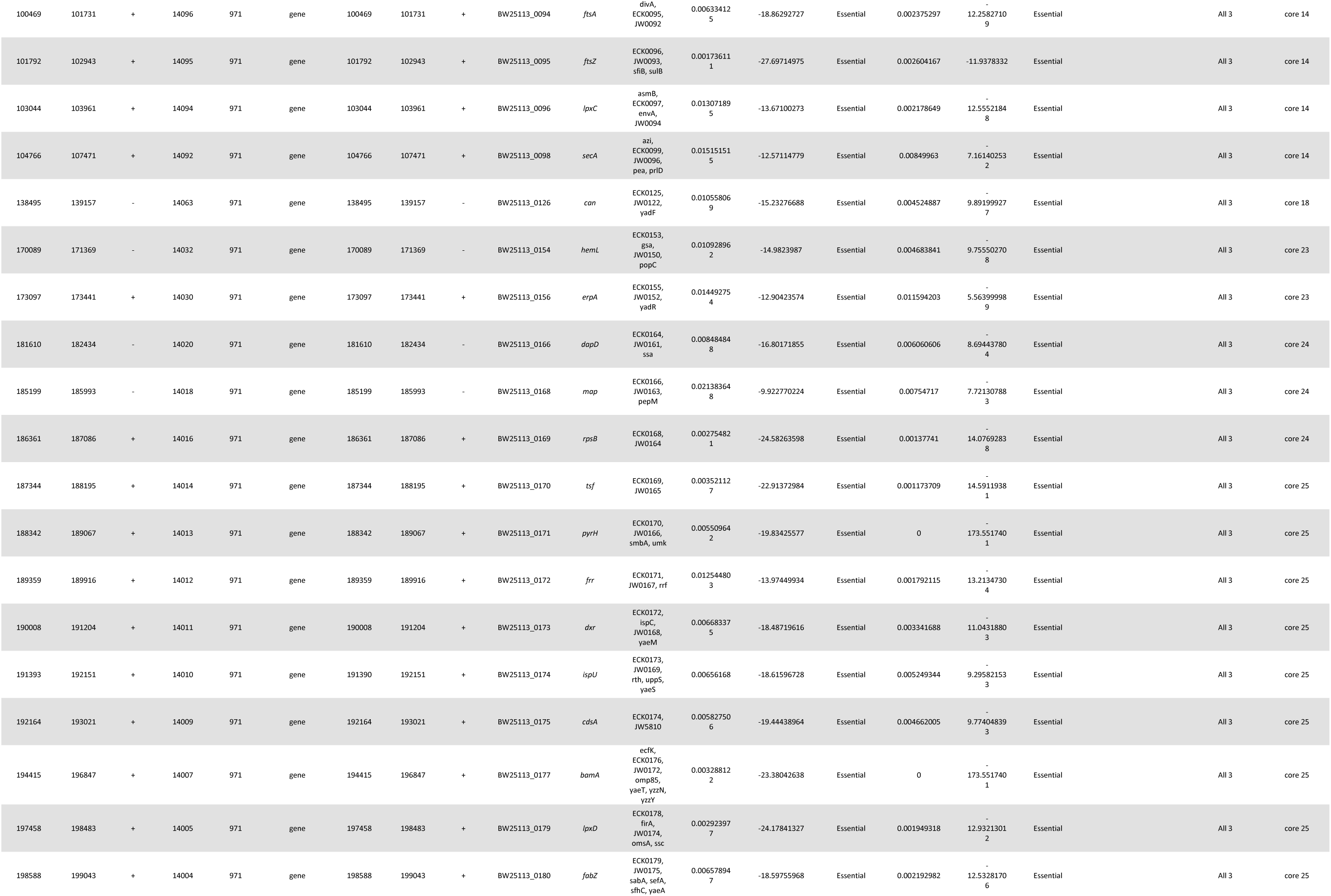
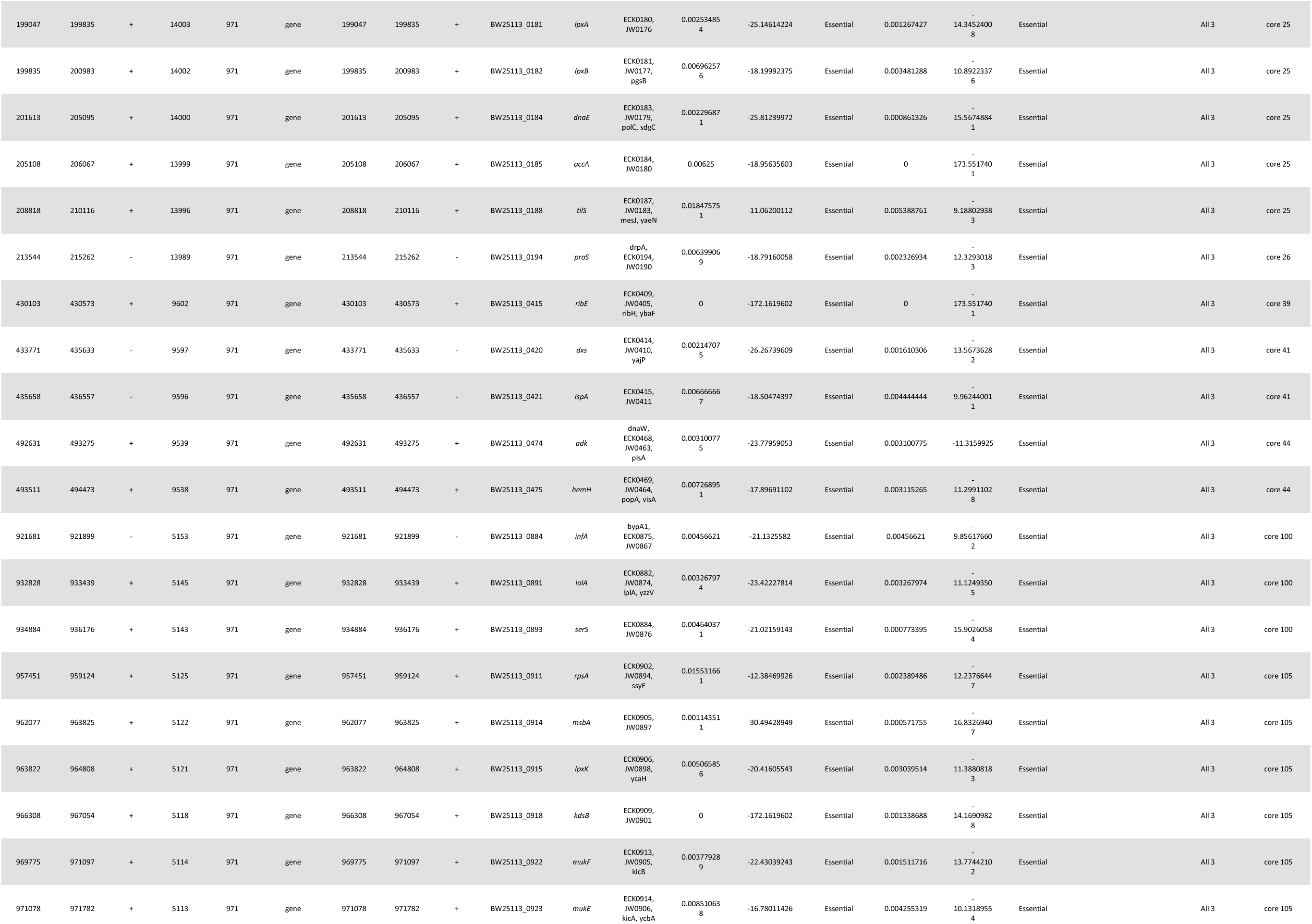
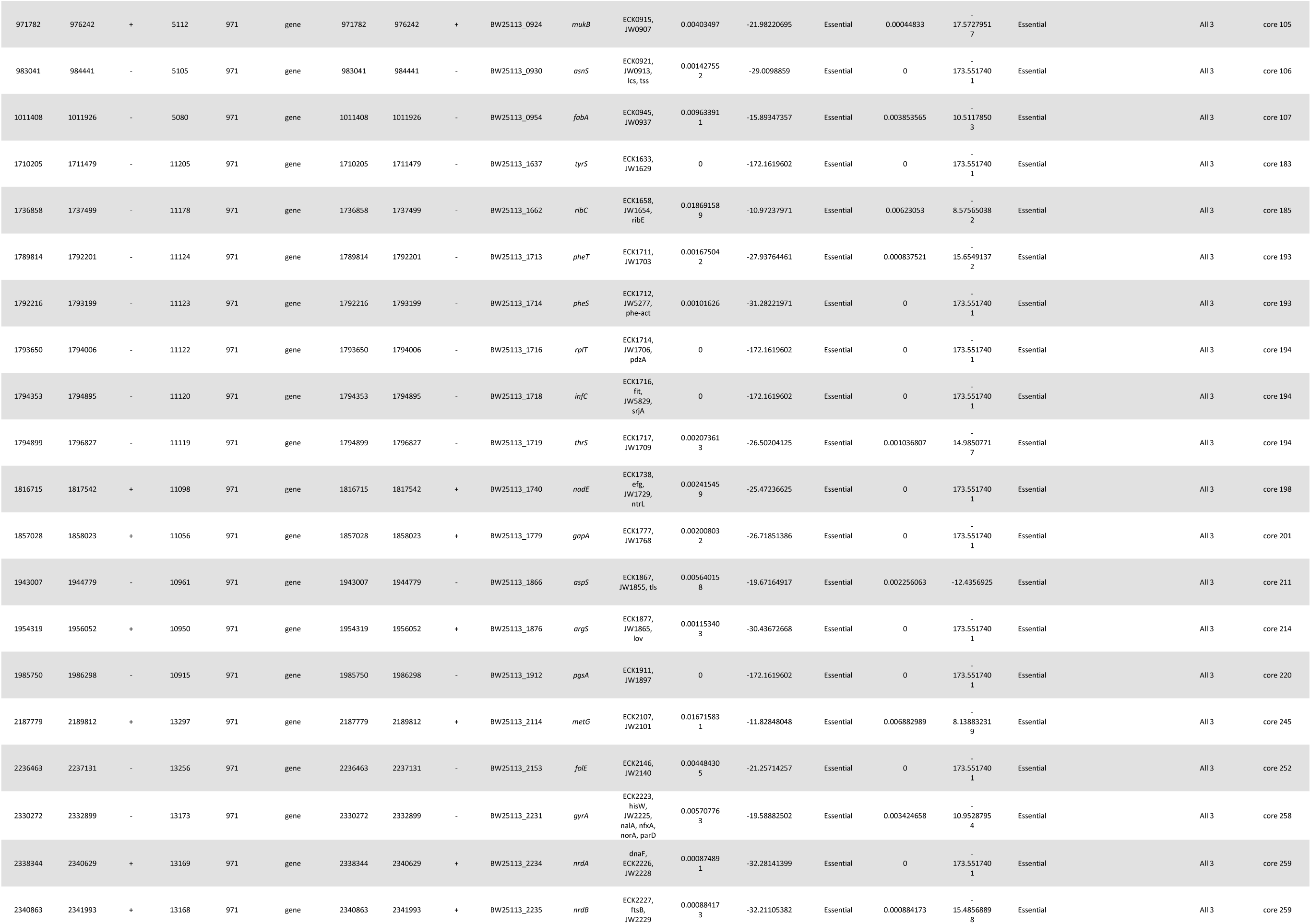
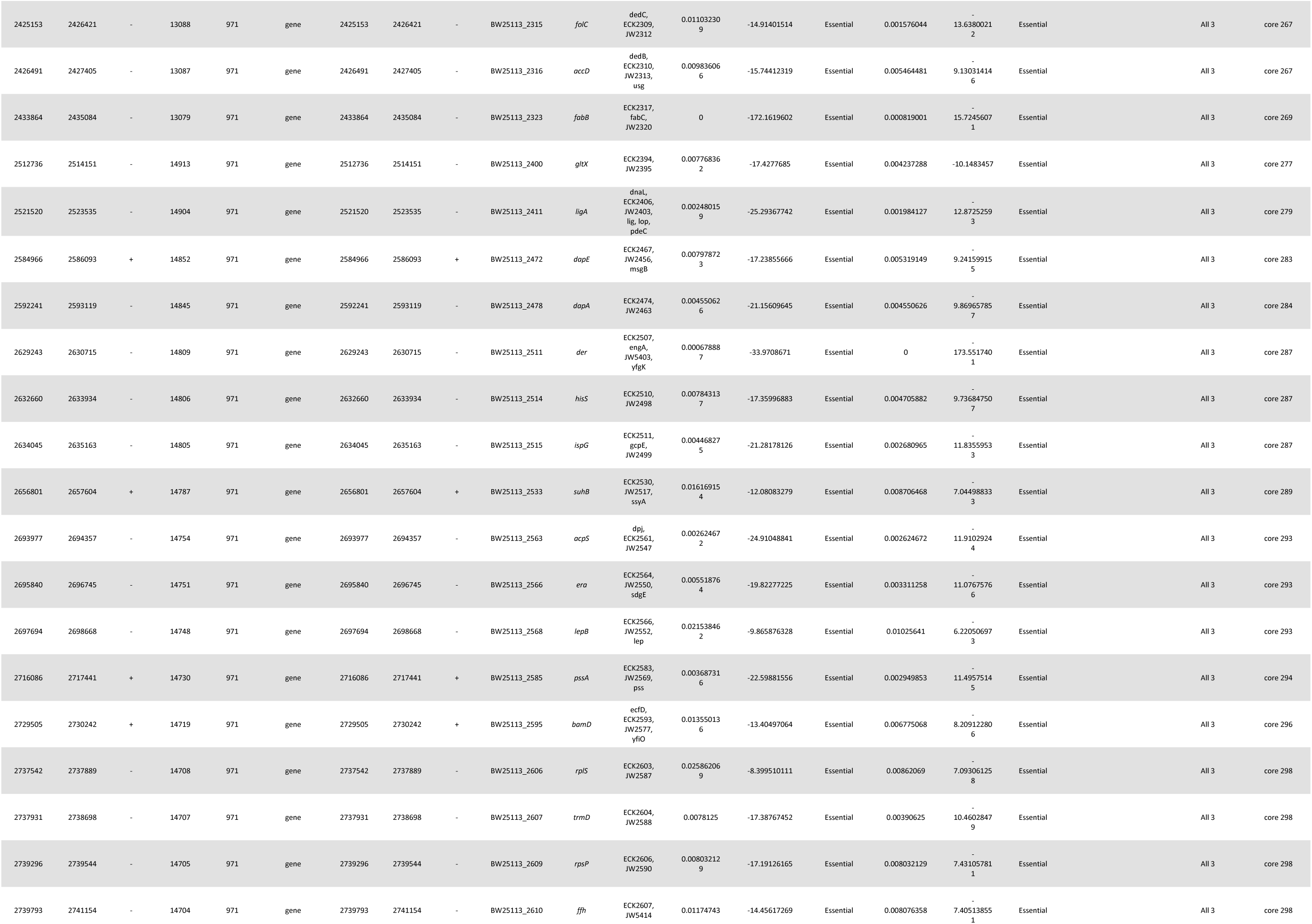
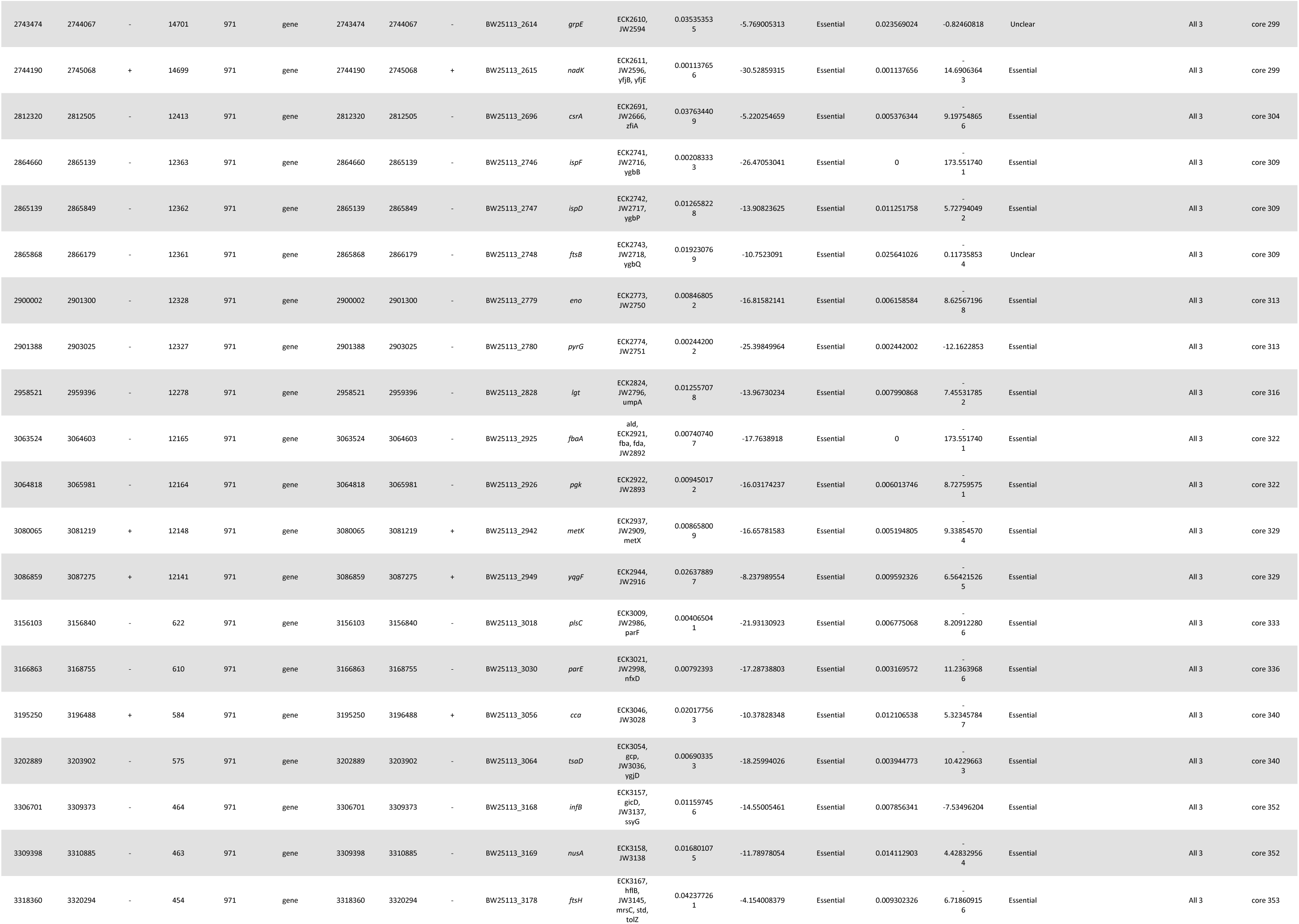
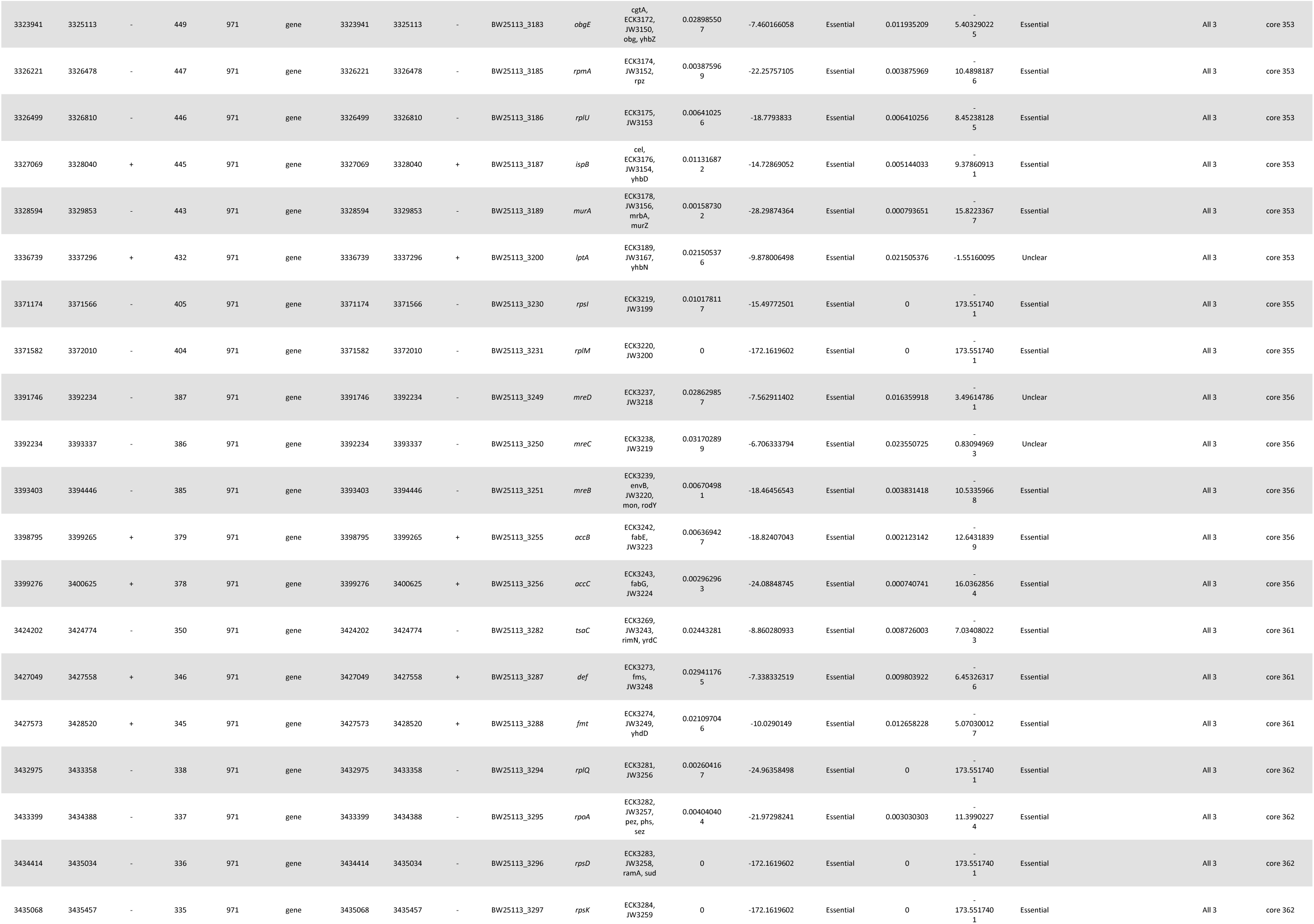
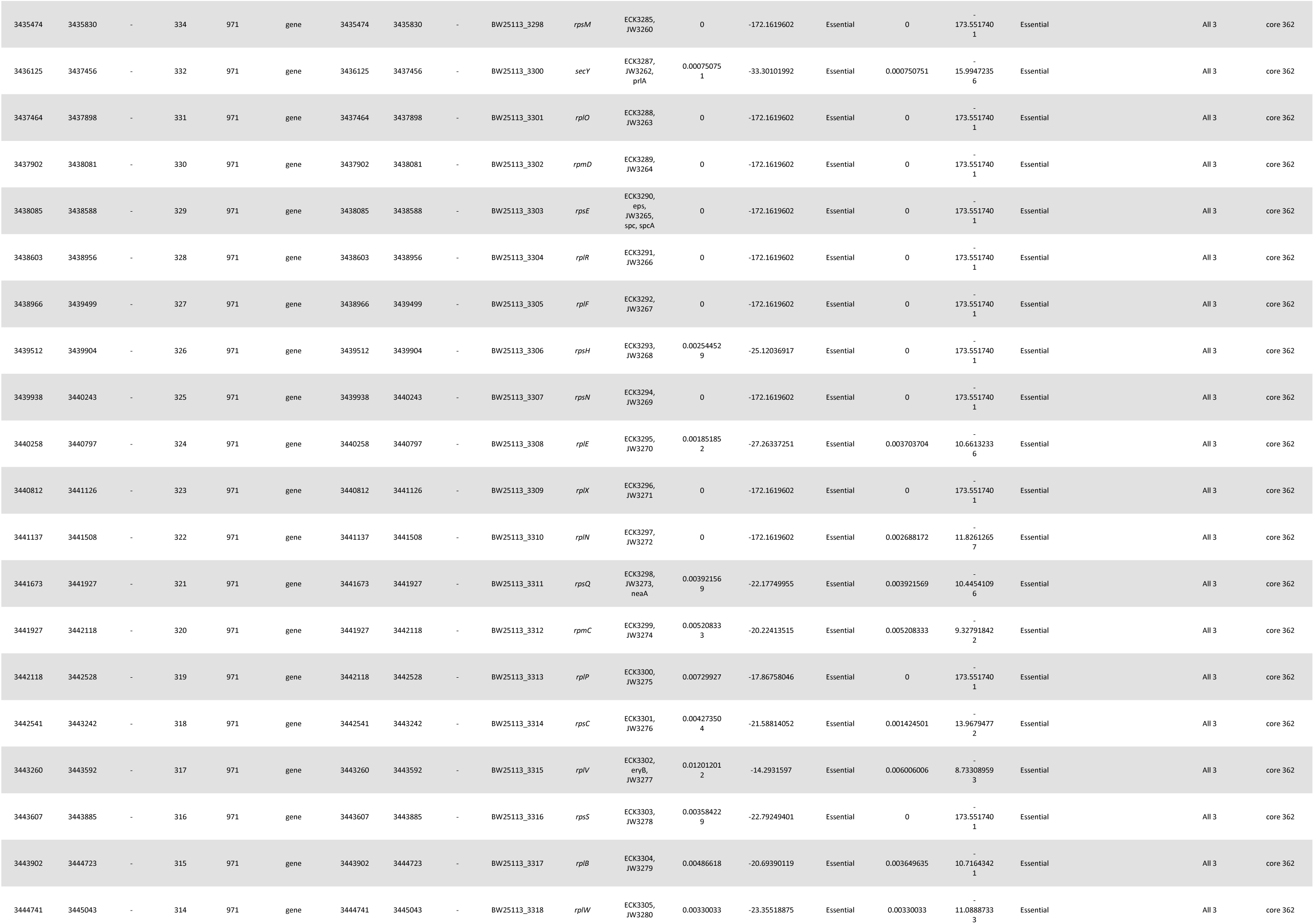
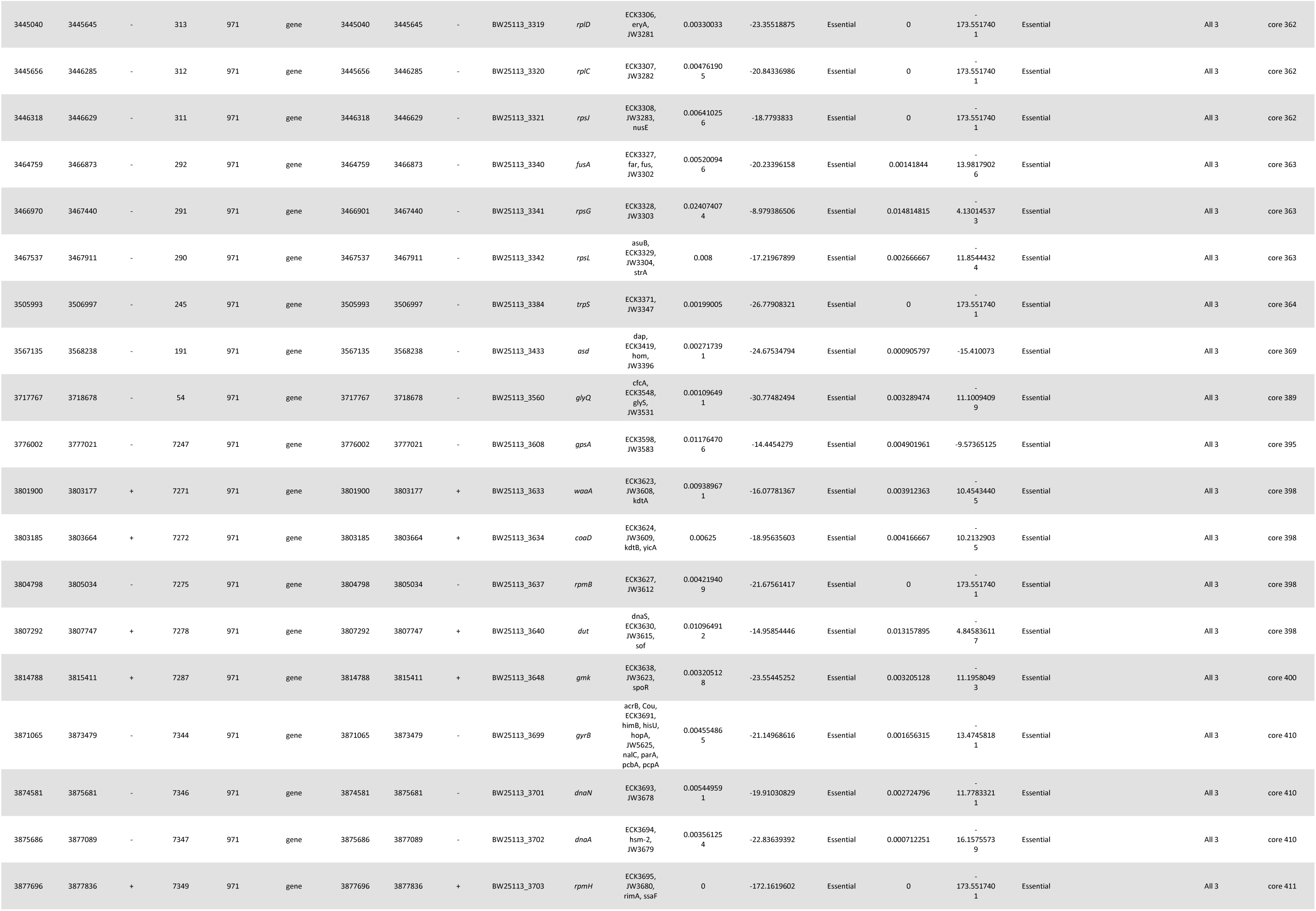
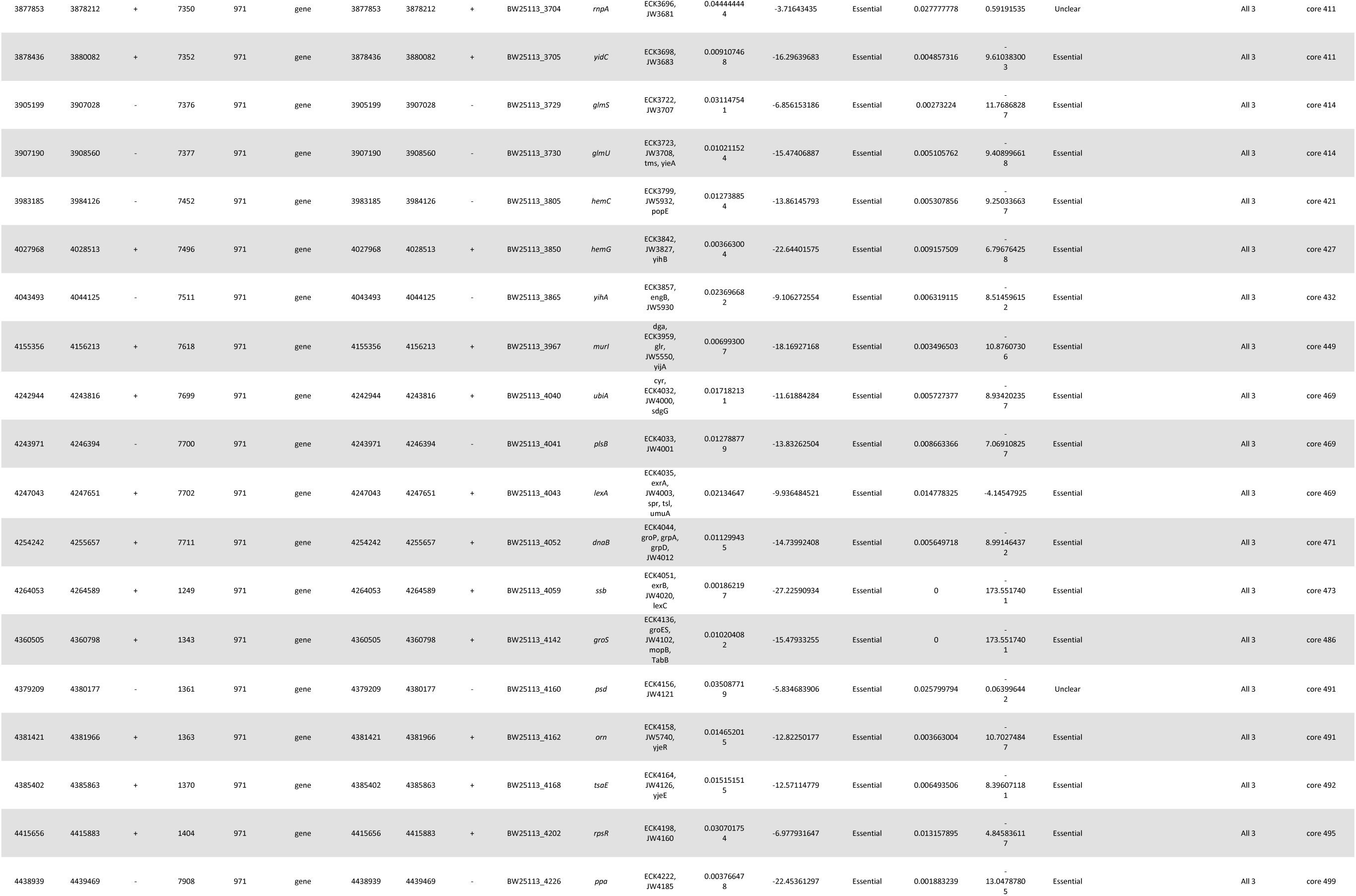
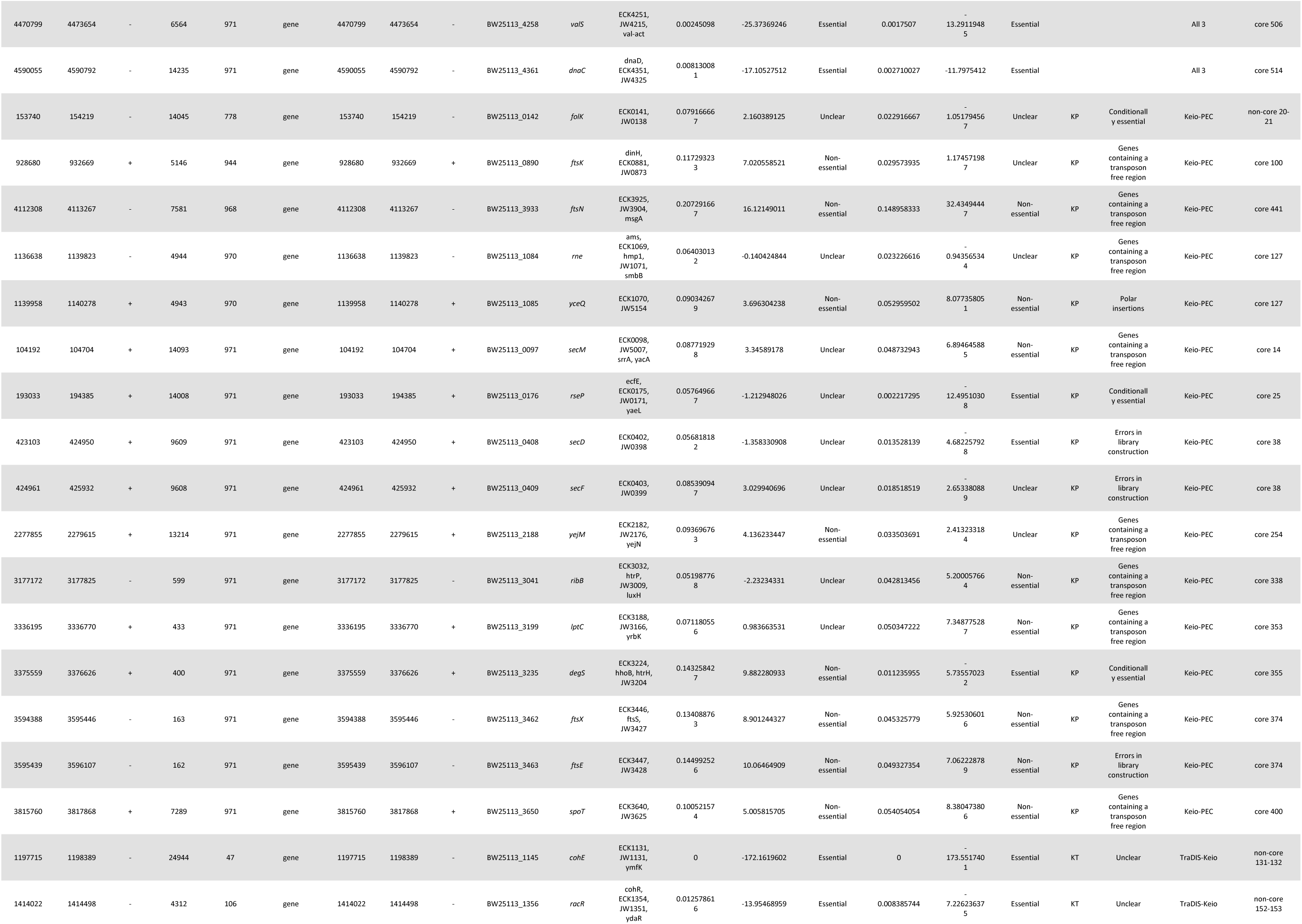
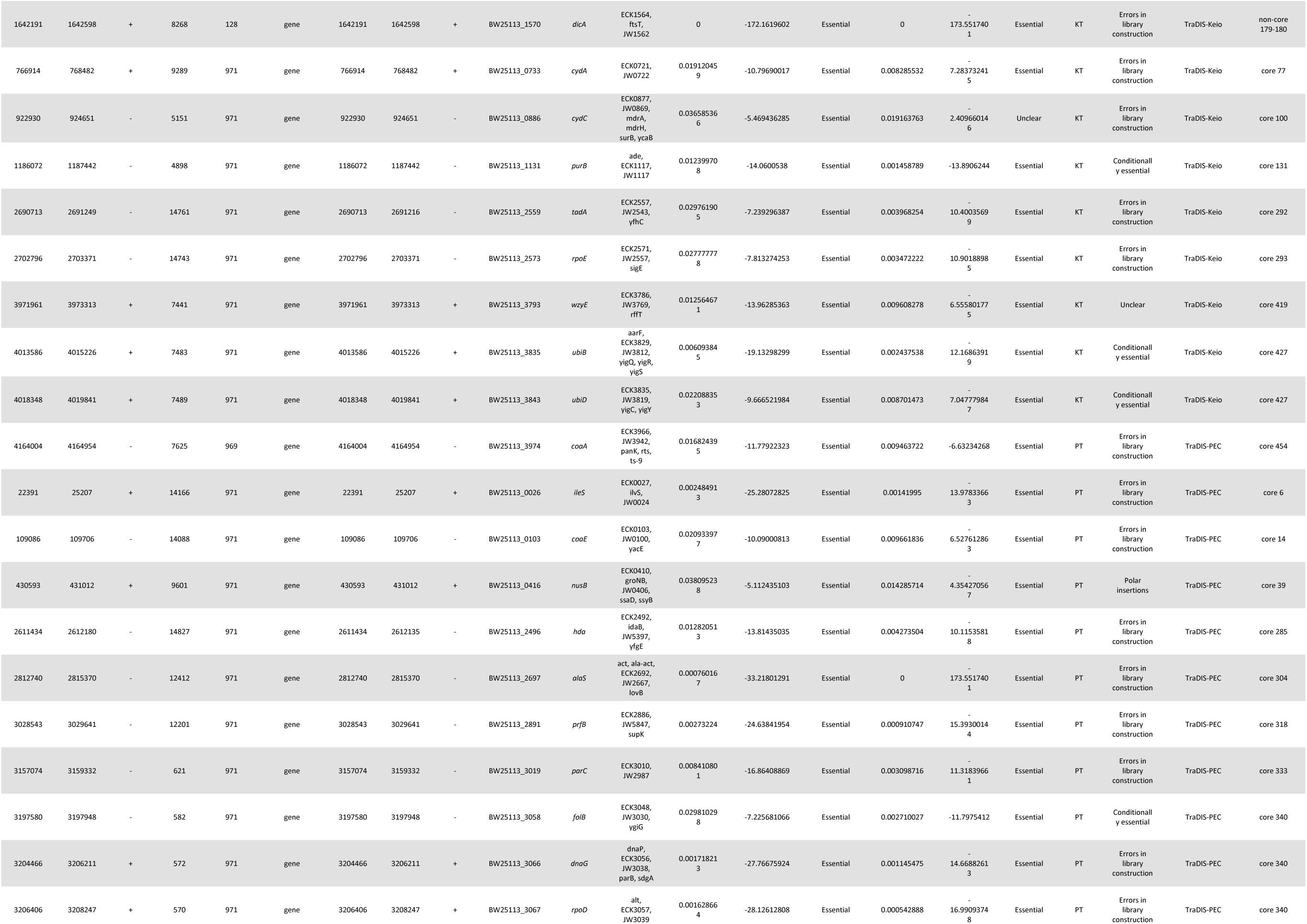
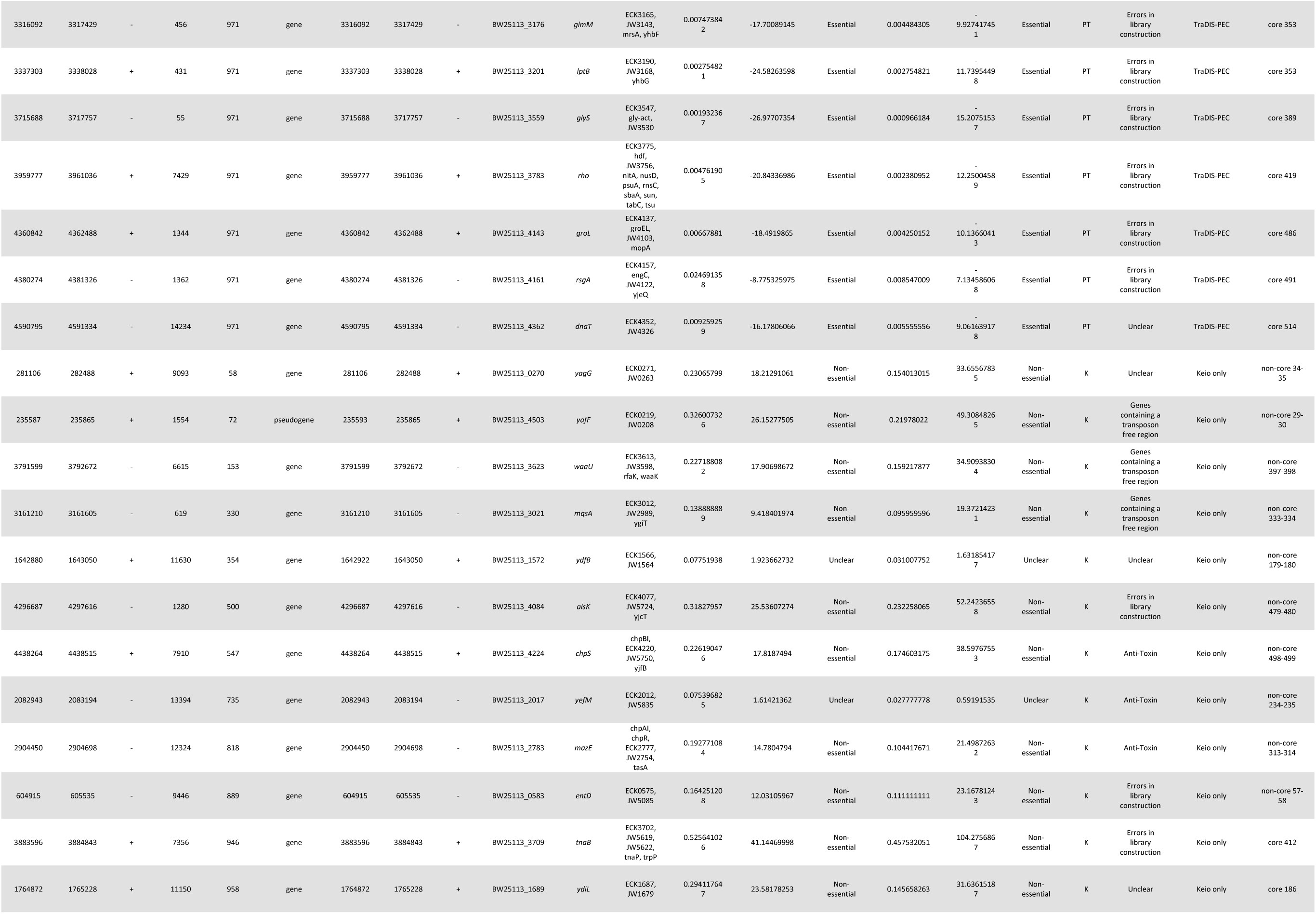
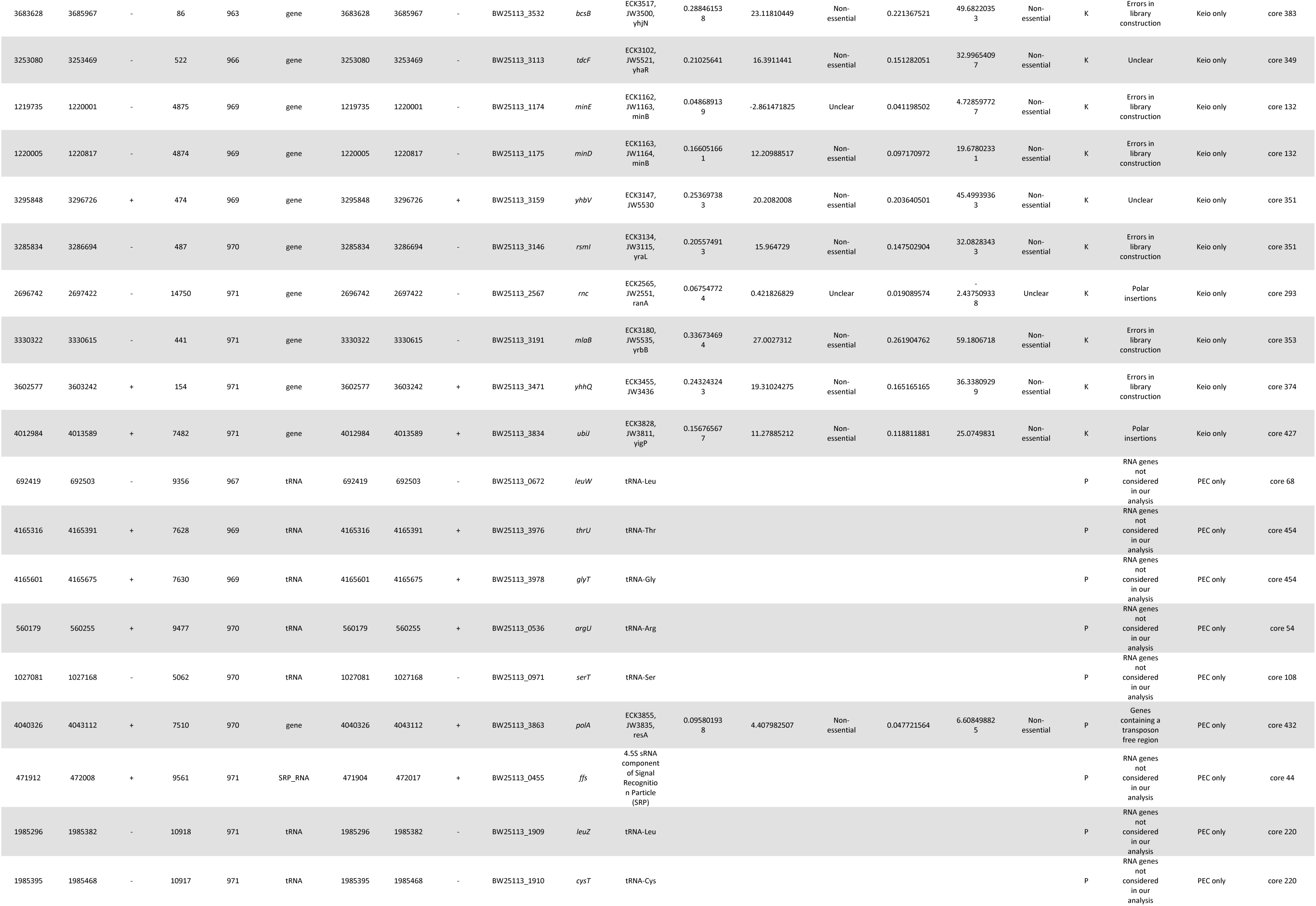
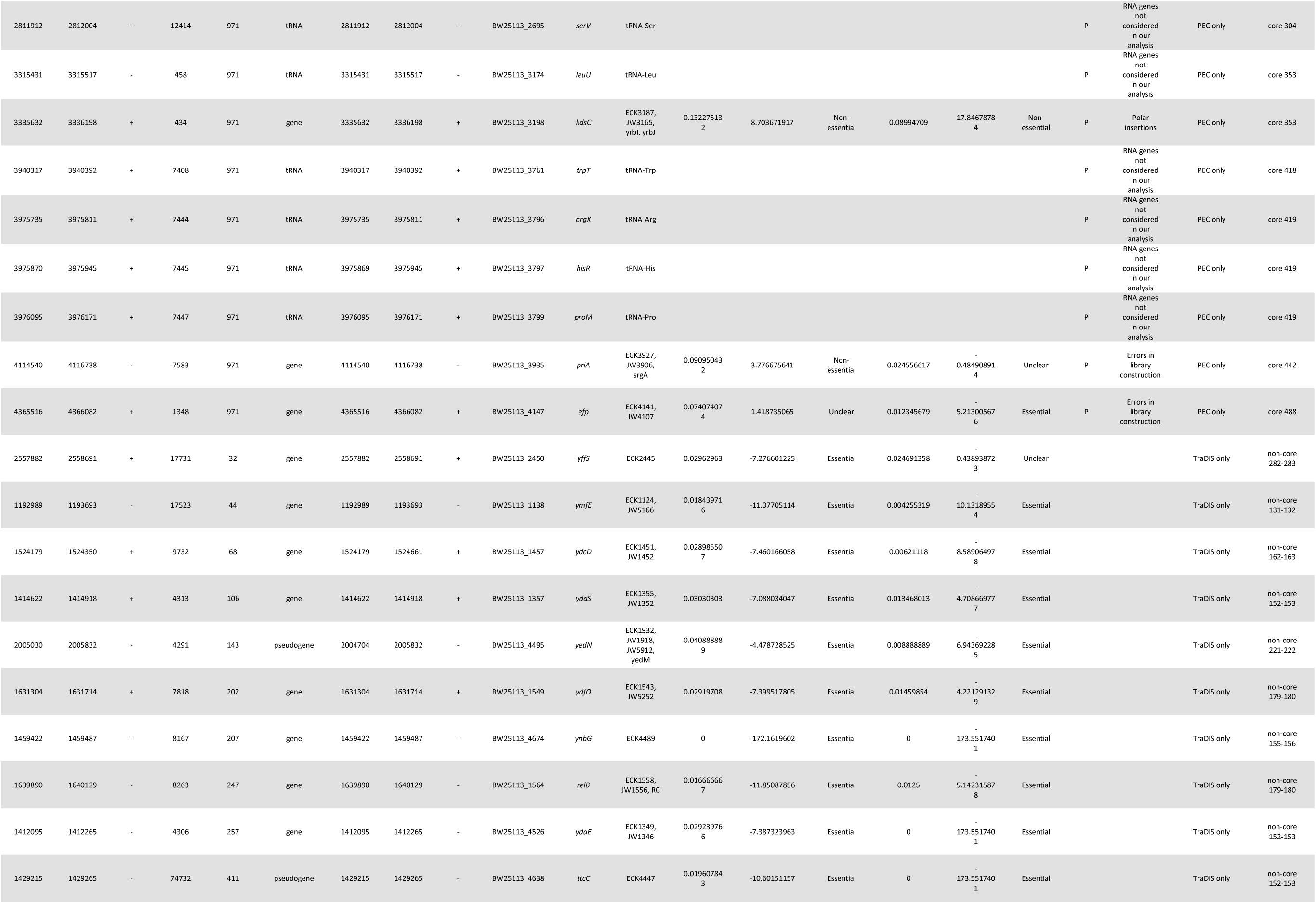
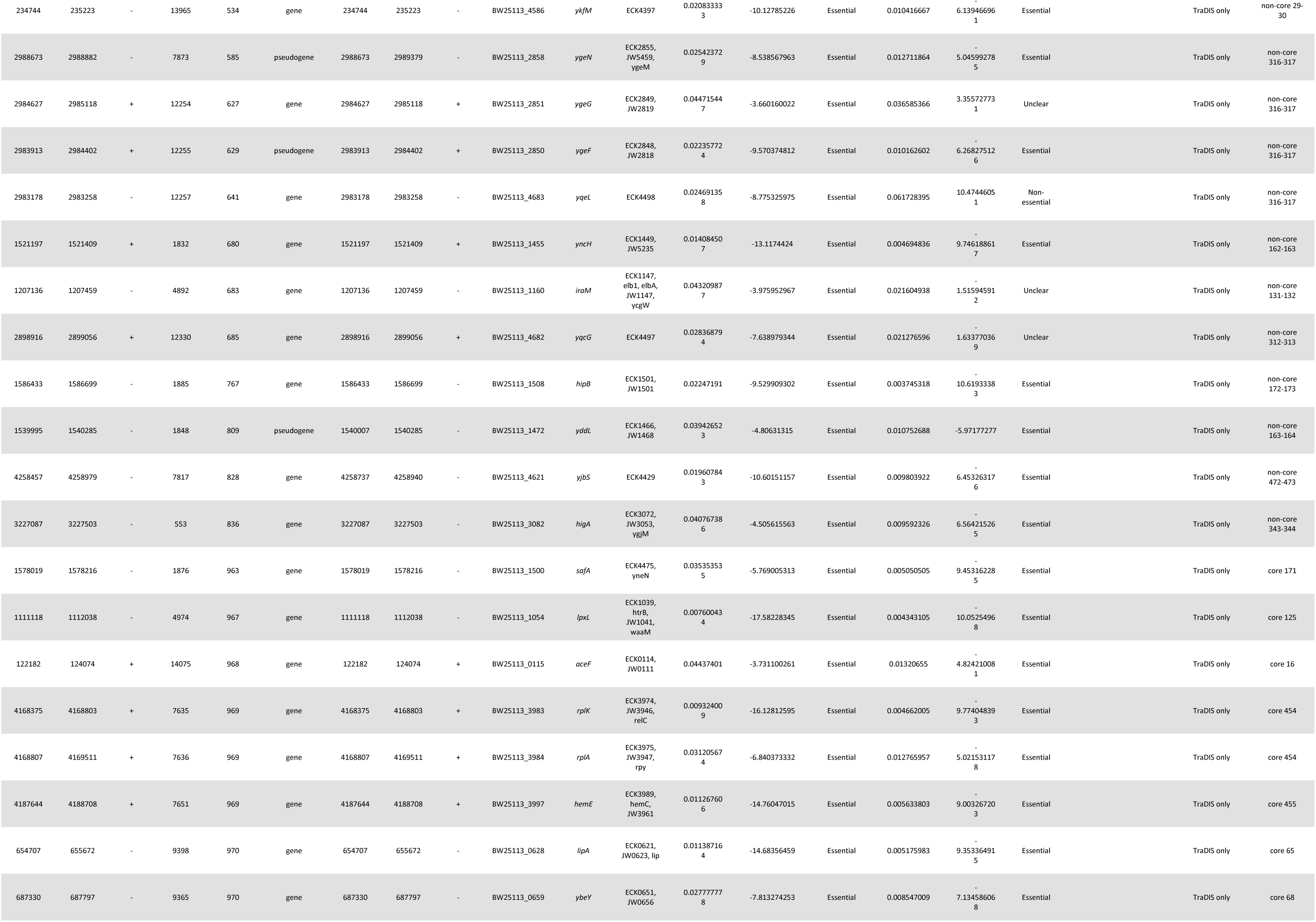
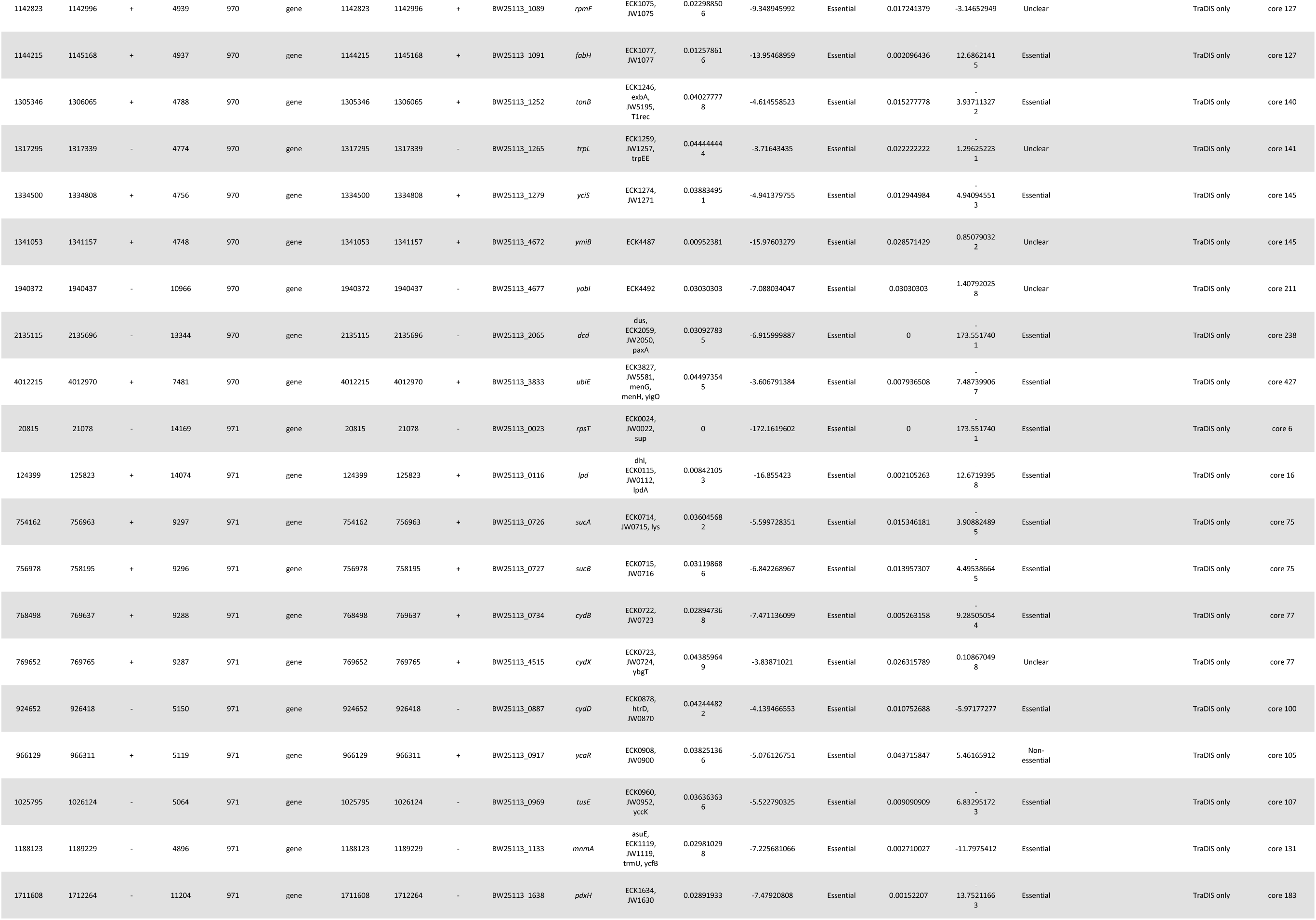
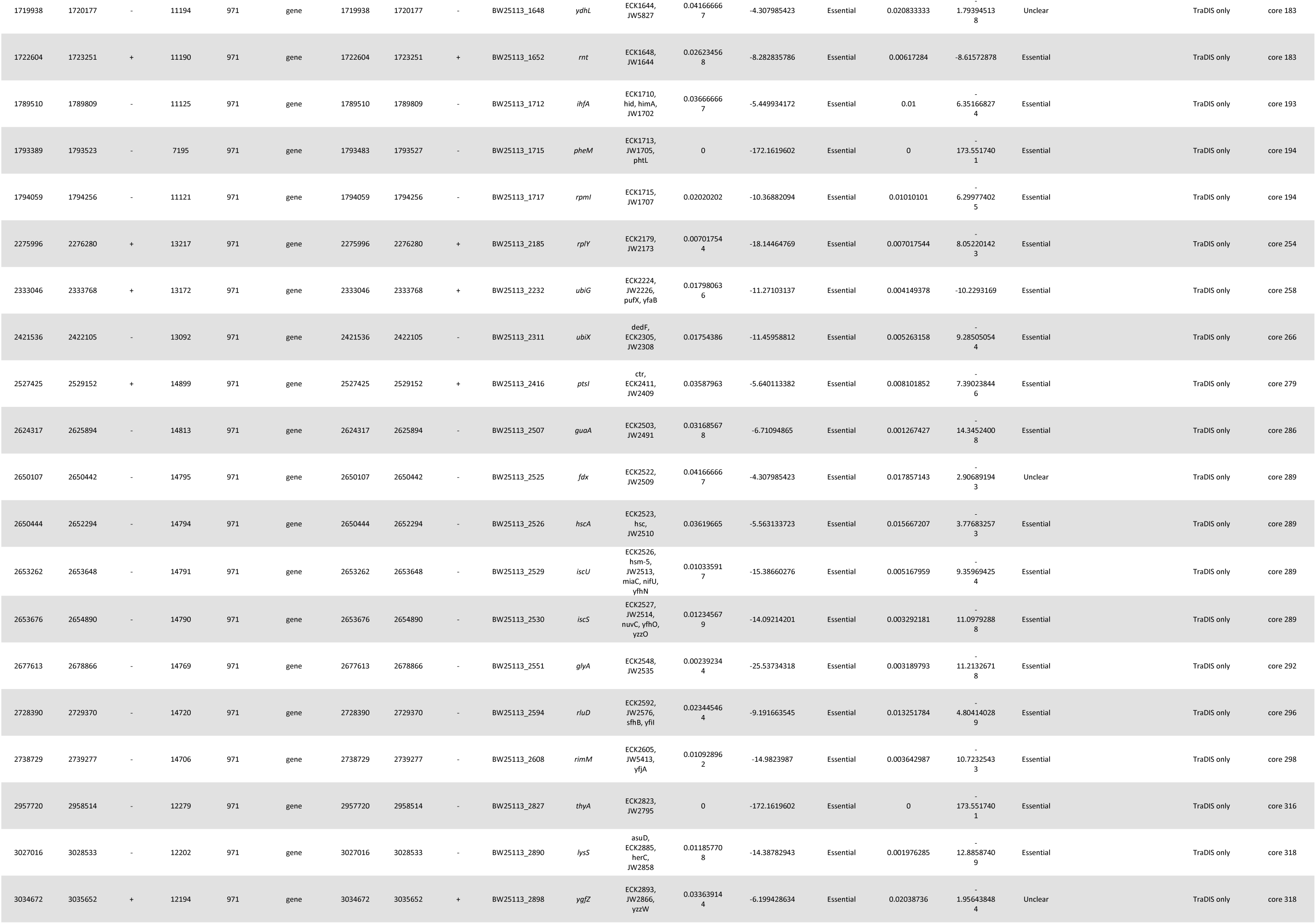
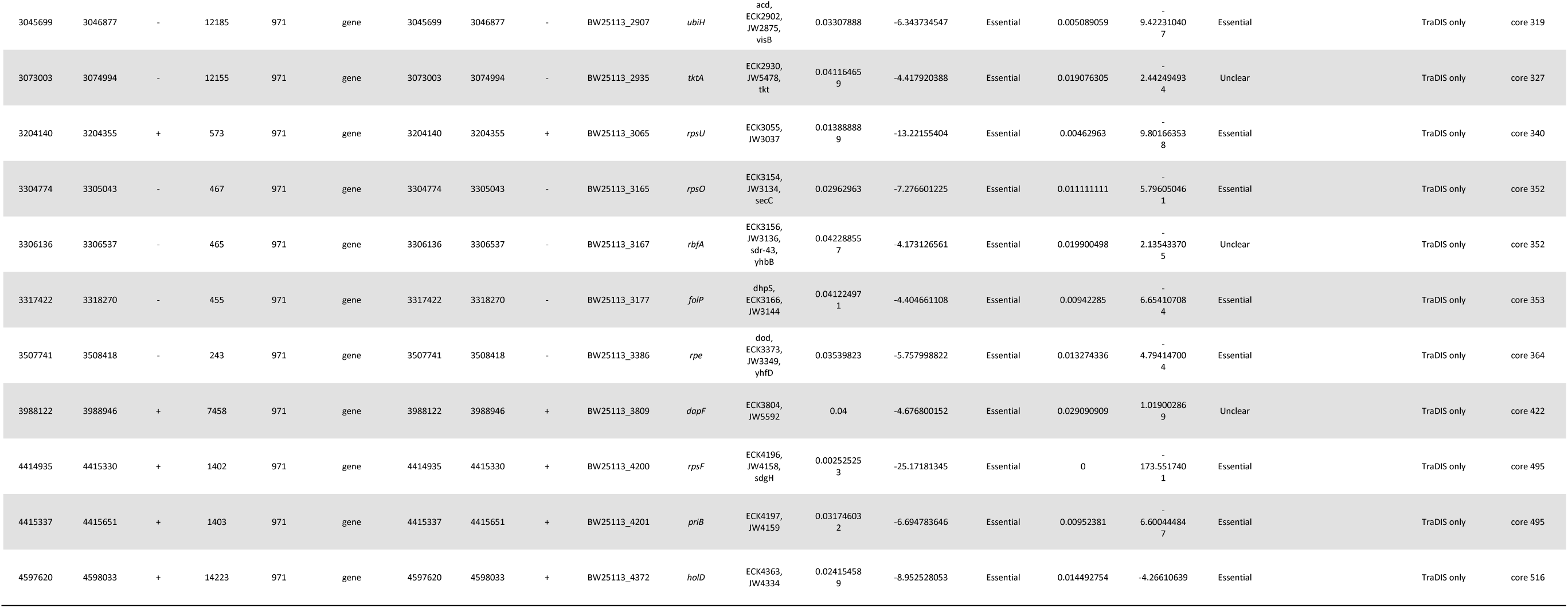
The 414 *E. coli* genes deemed essential by Goodall et al., Baba et al., or Yamazaki et al. These genes are compared to the PGG based annotation of core regions for the K-12 BW25113 strain used by Goodall (GenBank sequence CP009273.1, Assembly ASM75055v1/GCA_000750555.1, BioSample SAMN03013572). Columns 1-5 are the start, stop, strand, cluster, and cluster size for the PGG annotation. Columns 6-11 are the gene type, start, stop, strand, locus tag, and gene symbol/name for the GenBank annotation. Column 12 is a list of gene synonyms for the gene from GenBank. Columns 13-21 are from Goodall et al.: 13-15 from Table S1 (normal essentiality), 16-18 from Table S4 (essentiality after outgrowth), 19-20 from Table S3 (outlier discrepancies), and 21 from Table S2 (comparison of data sets). Column 22 is the PGG core or non-core region the gene is contained in.

**Table 4.** The 521 *E. coli* core regions identified through the pan-genome graph.

| Core Region | Start | Stop | Length (bp) |
| --- | --- | --- | --- |
| 1 | 337 | 5237 | 4901 |
| 2 | 5683 | 9893 | 4211 |
| 3 | 9928 | 15298 | 5371 |
| 4 | 16751 | 16960 | 210 |
| 5 | 17325 | 18655 | 1331 |
| 6 | 20815 | 28207 | 7393 |
| 7 | 28374 | 39115 | 10742 |
| 8 | 39244 | 50302 | 11059 |
| 9 | 50380 | 58179 | 7800 |
| 10 | 59687 | 66546 | 6860 |
| 11 | 66477 | 67752 | 1276 |
| 12 | 67838 | 72097 | 4260 |
| 13 | 72131 | 73786 | 1656 |
| 14 | 75335 | 113586 | 38252 |
| 15 | 113596 | 118038 | 4443 |
| 16 | 118579 | 125823 | 7245 |
| 17 | 127869 | 130699 | 2831 |
| 18 | 130875 | 143343 | 12469 |
| 19 | 143455 | 144357 | 903 |
| 20 | 144431 | 146088 | 1658 |
| 21 | 154216 | 161021 | 6806 |
| 22 | 161217 | 163751 | 2535 |
| 23 | 166265 | 180582 | 14318 |
| 24 | 181610 | 187086 | 5477 |
| 25 | 187344 | 212466 | 25123 |
| 26 | 212666 | 220198 | 7533 |
| 27 | 220258 | 222062 | 1805 |
| 28 | 222237 | 225166 | 2930 |
| 29 | 225245 | 233494 | 8250 |
| 30 | 235906 | 242292 | 6387 |
| 31 | 244845 | 246584 | 1740 |
| 32 | 247385 | 248440 | 1056 |
| 33 | 249196 | 249648 | 453 |
| 34 | 250746 | 258657 | 7912 |
| 35 | 380688 | 385435 | 4748 |
| 36 | 390586 | 403917 | 13332 |
| 37 | 404403 | 419788 | 15386 |
| 38 | 419793 | 425932 | 6140 |
| 39 | 426061 | 432563 | 6503 |
| 40 | 432617 | 433591 | 975 |
| 41 | 433771 | 442122 | 8352 |
| 42 | 442271 | 447066 | 4796 |
| 43 | 447110 | 447302 | 193 |
| 44 | 447526 | 502022 | 54497 |
| 45 | 502059 | 503536 | 1478 |
| 46 | 504331 | 509324 | 4994 |
| 47 | 509449 | 509856 | 408 |
| 48 | 509857 | 518286 | 8430 |
| 49 | 525588 | 531153 | 5566 |
| 50 | 543813 | 545072 | 1260 |
| 51 | 545083 | 546788 | 1706 |
| 52 | 546983 | 548643 | 1661 |
| 53 | 548674 | 553197 | 4524 |
| 54 | 560179 | 560255 | 77 |
| 55 | 588766 | 592919 | 4154 |
| 56 | 594170 | 600880 | 6711 |
| 57 | 601721 | 602839 | 1119 |
| 58 | 605480 | 613494 | 8015 |
| 59 | 613710 | 619966 | 6257 |
| 60 | 620341 | 630832 | 10492 |
| 61 | 630805 | 632025 | 1221 |
| 62 | 633160 | 637323 | 4164 |
| 63 | 639013 | 647312 | 8300 |
| 64 | 650039 | 651792 | 1754 |
| 65 | 652013 | 655884 | 3872 |
| 66 | 657093 | 670956 | 13864 |
| 67 | 678933 | 683203 | 4271 |
| 68 | 684799 | 692826 | 8028 |
| 69 | 692969 | 694633 | 1665 |
| 70 | 695030 | 701346 | 6317 |
| 71 | 701549 | 703213 | 1665 |
| 72 | 703790 | 710654 | 6865 |
| 73 | 716512 | 724796 | 8285 |
| 74 | 734457 | 743225 | 8769 |
| 75 | 748641 | 758195 | 9555 |
| 76 | 758470 | 760505 | 2036 |
| 77 | 766914 | 776374 | 9461 |
| 78 | 777033 | 777108 | 76 |
| 79 | 777224 | 793887 | 16664 |
| 80 | 794042 | 796031 | 1990 |
| 81 | 798959 | 802737 | 3779 |
| 82 | 802795 | 803365 | 571 |
| 83 | 803424 | 811003 | 7580 |
| 84 | 811195 | 816044 | 4850 |
| 85 | 816998 | 827692 | 10695 |
| 86 | 828526 | 832892 | 4367 |
| 87 | 833121 | 836987 | 3867 |
| 88 | 837006 | 853011 | 16006 |
| 89 | 854669 | 868257 | 13589 |
| 90 | 868435 | 870783 | 2349 |
| 91 | 870791 | 872119 | 1329 |
| 92 | 872166 | 883415 | 11250 |
| 93 | 883435 | 895101 | 11667 |
| 94 | 895300 | 899190 | 3891 |
| 95 | 900049 | 910426 | 10378 |
| 96 | 910808 | 911503 | 696 |
| 97 | 911716 | 918270 | 6555 |
| 98 | 918369 | 920996 | 2628 |
| 99 | 921274 | 921358 | 85 |
| 100 | 921340 | 936176 | 14837 |
| 101 | 936415 | 940352 | 3938 |
| 102 | 940387 | 942475 | 2089 |
| 103 | 945796 | 946536 | 741 |
| 104 | 946728 | 955551 | 8824 |
| 105 | 955720 | 979753 | 24034 |
| 106 | 979975 | 992968 | 12994 |
| 107 | 1000224 | 1026874 | 26651 |
| 108 | 1027081 | 1037371 | 10291 |
| 109 | 1046419 | 1046631 | 213 |
| 110 | 1046917 | 1051634 | 4718 |
| 111 | 1051717 | 1059231 | 7515 |
| 112 | 1061041 | 1063164 | 2124 |
| 113 | 1063368 | 1063604 | 237 |
| 114 | 1065316 | 1067615 | 2300 |
| 115 | 1067627 | 1074602 | 6976 |
| 116 | 1074761 | 1076269 | 1509 |
| 117 | 1076812 | 1080103 | 3292 |
| 118 | 1080448 | 1081512 | 1065 |
| 119 | 1093021 | 1096243 | 3223 |
| 120 | 1096307 | 1097140 | 834 |
| 121 | 1097167 | 1098652 | 1486 |
| 122 | 1098868 | 1099304 | 437 |
| 123 | 1099407 | 1100358 | 952 |
| 124 | 1100417 | 1108862 | 8446 |
| 125 | 1110976 | 1114502 | 3527 |
| 126 | 1114763 | 1136442 | 21680 |
| 127 | 1136638 | 1162845 | 26208 |
| 128 | 1163055 | 1175797 | 12743 |
| 129 | 1177161 | 1179076 | 1916 |
| 130 | 1179073 | 1183696 | 4624 |
| 131 | 1183772 | 1191829 | 8058 |
| 132 | 1219409 | 1222928 | 3520 |
| 133 | 1223170 | 1224985 | 1816 |
| 134 | 1226052 | 1235405 | 9354 |
| 135 | 1235791 | 1240438 | 4648 |
| 136 | 1252177 | 1264475 | 12299 |
| 137 | 1265159 | 1281982 | 16824 |
| 138 | 1282994 | 1289600 | 6607 |
| 139 | 1290902 | 1305357 | 14456 |
| 140 | 1305346 | 1308037 | 2692 |
| 141 | 1311479 | 1318975 | 7497 |
| 142 | 1321109 | 1321984 | 876 |
| 143 | 1322024 | 1329086 | 7063 |
| 144 | 1329381 | 1329715 | 335 |
| 145 | 1330088 | 1345296 | 15209 |
| 146 | 1346085 | 1351925 | 5841 |
| 147 | 1361192 | 1364260 | 3069 |
| 148 | 1377220 | 1384983 | 7764 |
| 149 | 1386248 | 1387147 | 900 |
| 150 | 1387484 | 1390179 | 2696 |
| 151 | 1391622 | 1394493 | 2872 |
| 152 | 1400236 | 1406205 | 5970 |
| 153 | 1429442 | 1431150 | 1709 |
| 154 | 1431517 | 1436000 | 4484 |
| 155 | 1436111 | 1439947 | 3837 |
| 156 | 1476512 | 1481220 | 4709 |
| 157 | 1483970 | 1486107 | 2138 |
| 158 | 1488405 | 1497382 | 8978 |
| 159 | 1509019 | 1511259 | 2241 |
| 160 | 1511356 | 1513284 | 1929 |
| 161 | 1515220 | 1517322 | 2103 |
| 162 | 1517356 | 1520237 | 2882 |
| 163 | 1533107 | 1538317 | 5211 |
| 164 | 1540545 | 1546248 | 5704 |
| 165 | 1546655 | 1546939 | 285 |
| 166 | 1547085 | 1550017 | 2933 |
| 167 | 1550882 | 1551313 | 432 |
| 168 | 1561761 | 1566678 | 4918 |
| 169 | 1573890 | 1575047 | 1158 |
| 170 | 1575099 | 1576781 | 1683 |
| 171 | 1577183 | 1580743 | 3561 |
| 172 | 1581077 | 1581991 | 915 |
| 173 | 1603229 | 1604937 | 1709 |
| 174 | 1605164 | 1606111 | 948 |
| 175 | 1606223 | 1608960 | 2738 |
| 176 | 1611285 | 1613165 | 1881 |
| 177 | 1613377 | 1614464 | 1088 |
| 178 | 1616903 | 1617171 | 269 |
| 179 | 1619514 | 1625225 | 5712 |
| 180 | 1650441 | 1651001 | 561 |
| 181 | 1651004 | 1652127 | 1124 |
| 182 | 1657247 | 1669204 | 11958 |
| 183 | 1669229 | 1723251 | 54023 |
| 184 | 1728011 | 1728358 | 348 |
| 185 | 1728692 | 1762942 | 34251 |
| 186 | 1763331 | 1766542 | 3212 |
| 187 | 1766769 | 1769701 | 2933 |
| 188 | 1772647 | 1773558 | 912 |
| 189 | 1773661 | 1774638 | 978 |
| 190 | 1774658 | 1778934 | 4277 |
| 191 | 1778991 | 1783738 | 4748 |
| 192 | 1783870 | 1789409 | 5540 |
| 193 | 1789510 | 1793199 | 3690 |
| 194 | 1793389 | 1796827 | 3439 |
| 195 | 1799219 | 1806582 | 7364 |
| 196 | 1807678 | 1810385 | 2708 |
| 197 | 1810643 | 1812757 | 2115 |
| 198 | 1812862 | 1837971 | 25110 |
| 199 | 1839256 | 1844950 | 5695 |
| 200 | 1845117 | 1846785 | 1669 |
| 201 | 1855959 | 1863099 | 7141 |
| 202 | 1863212 | 1864495 | 1284 |
| 203 | 1866118 | 1866285 | 168 |
| 204 | 1867831 | 1873512 | 5682 |
| 205 | 1873846 | 1874205 | 360 |
| 206 | 1881121 | 1888139 | 7019 |
| 207 | 1888330 | 1916569 | 28240 |
| 208 | 1916570 | 1917480 | 911 |
| 209 | 1917622 | 1933348 | 15727 |
| 210 | 1933479 | 1938456 | 4978 |
| 211 | 1938603 | 1944779 | 6177 |
| 212 | 1946523 | 1948670 | 2148 |
| 213 | 1948835 | 1951264 | 2430 |
| 214 | 1951289 | 1956052 | 4764 |
| 215 | 1956837 | 1960448 | 3612 |
| 216 | 1960650 | 1963621 | 2972 |
| 217 | 1965287 | 1972101 | 6815 |
| 218 | 1973234 | 1975093 | 1860 |
| 219 | 1975068 | 1985100 | 10033 |
| 220 | 1985296 | 1995270 | 9975 |
| 221 | 1998784 | 2003347 | 4564 |
| 222 | 2006181 | 2006734 | 554 |
| 223 | 2016374 | 2017159 | 786 |
| 224 | 2017449 | 2025798 | 8350 |
| 225 | 2029316 | 2034599 | 5284 |
| 226 | 2034856 | 2035506 | 651 |
| 227 | 2036949 | 2038105 | 1157 |
| 228 | 2048386 | 2051468 | 3083 |
| 229 | 2051508 | 2055816 | 4309 |
| 230 | 2055872 | 2059245 | 3374 |
| 231 | 2072513 | 2072842 | 330 |
| 232 | 2074862 | 2076028 | 1167 |
| 233 | 2076237 | 2077664 | 1428 |
| 234 | 2079185 | 2082609 | 3425 |
| 235 | 2083280 | 2083462 | 183 |
| 236 | 2083543 | 2090706 | 7164 |
| 237 | 2131383 | 2132966 | 1584 |
| 238 | 2133240 | 2144466 | 11227 |
| 239 | 2145192 | 2146609 | 1418 |
| 240 | 2147497 | 2159002 | 11506 |
| 241 | 2159149 | 2160678 | 1530 |
| 242 | 2161470 | 2163092 | 1623 |
| 243 | 2170991 | 2178911 | 7921 |
| 244 | 2179003 | 2180777 | 1775 |
| 245 | 2186272 | 2189812 | 3541 |
| 246 | 2199174 | 2204579 | 5406 |
| 247 | 2204704 | 2208123 | 3420 |
| 248 | 2209136 | 2212036 | 2901 |
| 249 | 2213171 | 2222544 | 9374 |
| 250 | 2222917 | 2223864 | 948 |
| 251 | 2224103 | 2227318 | 3216 |
| 252 | 2227512 | 2245176 | 17665 |
| 253 | 2247724 | 2249493 | 1770 |
| 254 | 2253198 | 2279766 | 26569 |
| 255 | 2283690 | 2305013 | 21324 |
| 256 | 2306509 | 2313355 | 6847 |
| 257 | 2320846 | 2327881 | 7036 |
| 258 | 2328435 | 2333768 | 5334 |
| 259 | 2338344 | 2342951 | 4608 |
| 260 | 2343414 | 2351282 | 7869 |
| 261 | 2355910 | 2367017 | 11108 |
| 262 | 2367127 | 2374506 | 7380 |
| 263 | 2375087 | 2376004 | 918 |
| 264 | 2380413 | 2380916 | 504 |
| 265 | 2383527 | 2416080 | 32554 |
| 266 | 2417215 | 2424242 | 7028 |
| 267 | 2424501 | 2427405 | 2905 |
| 268 | 2428303 | 2431330 | 3028 |
| 269 | 2431310 | 2437249 | 5940 |
| 270 | 2437370 | 2441919 | 4550 |
| 271 | 2449806 | 2454749 | 4944 |
| 272 | 2457731 | 2459712 | 1982 |
| 273 | 2459788 | 2459862 | 75 |
| 274 | 2472681 | 2491774 | 19094 |
| 275 | 2492150 | 2507723 | 15574 |
| 276 | 2509122 | 2512290 | 3169 |
| 277 | 2512736 | 2514485 | 1750 |
| 278 | 2514612 | 2514687 | 76 |
| 279 | 2519289 | 2531070 | 11782 |
| 280 | 2532031 | 2537982 | 5952 |
| 281 | 2539132 | 2541456 | 2325 |
| 282 | 2543005 | 2550656 | 7652 |
| 283 | 2568337 | 2590096 | 21760 |
| 284 | 2590264 | 2594307 | 4044 |
| 285 | 2608179 | 2624224 | 16046 |
| 286 | 2624317 | 2628961 | 4645 |
| 287 | 2629243 | 2638223 | 8981 |
| 288 | 2638372 | 2646698 | 8327 |
| 289 | 2647215 | 2661205 | 13991 |
| 290 | 2666627 | 2666720 | 94 |
| 291 | 2666705 | 2667127 | 423 |
| 292 | 2667175 | 2692966 | 25792 |
| 293 | 2693022 | 2707589 | 14568 |
| 294 | 2707798 | 2719199 | 11402 |
| 295 | 2719430 | 2724917 | 5488 |
| 296 | 2724959 | 2730242 | 5284 |
| 297 | 2730272 | 2735752 | 5481 |
| 298 | 2735742 | 2743419 | 7678 |
| 299 | 2743474 | 2749314 | 5841 |
| 300 | 2782344 | 2793834 | 11491 |
| 301 | 2793879 | 2807513 | 13635 |
| 302 | 2807577 | 2810862 | 3286 |
| 303 | 2811143 | 2811219 | 77 |
| 304 | 2811832 | 2815998 | 4167 |
| 305 | 2816067 | 2817705 | 1639 |
| 306 | 2817850 | 2831464 | 13615 |
| 307 | 2835932 | 2849775 | 13844 |
| 308 | 2849812 | 2853013 | 3202 |
| 309 | 2859918 | 2866388 | 6471 |
| 310 | 2866746 | 2870977 | 4232 |
| 311 | 2880669 | 2885938 | 5270 |
| 312 | 2898106 | 2898777 | 672 |
| 313 | 2900002 | 2904044 | 4043 |
| 314 | 2904776 | 2927047 | 22272 |
| 315 | 2928943 | 2940932 | 11990 |
| 316 | 2941116 | 2980435 | 39320 |
| 317 | 2992343 | 2997327 | 4985 |
| 318 | 3024593 | 3038460 | 13868 |
| 319 | 3039527 | 3050308 | 10782 |
| 320 | 3050321 | 3052684 | 2364 |
| 321 | 3053112 | 3054216 | 1105 |
| 322 | 3059636 | 3067105 | 7470 |
| 323 | 3068045 | 3068554 | 510 |
| 324 | 3069538 | 3070815 | 1278 |
| 325 | 3070830 | 3072218 | 1389 |
| 326 | 3072246 | 3072689 | 444 |
| 327 | 3073003 | 3076030 | 3028 |
| 328 | 3076236 | 3077156 | 921 |
| 329 | 3077294 | 3091762 | 14469 |
| 330 | 3091917 | 3103800 | 11884 |
| 331 | 3129667 | 3132952 | 3286 |
| 332 | 3133075 | 3154505 | 21431 |
| 333 | 3154616 | 3159519 | 4904 |
| 334 | 3162108 | 3165192 | 3085 |
| 335 | 3165889 | 3166815 | 927 |
| 336 | 3166863 | 3174940 | 8078 |
| 337 | 3174978 | 3176682 | 1705 |
| 338 | 3177172 | 3178077 | 906 |
| 339 | 3185098 | 3185562 | 465 |
| 340 | 3188323 | 3209032 | 20710 |
| 341 | 3209086 | 3209850 | 765 |
| 342 | 3210138 | 3219530 | 9393 |
| 343 | 3219593 | 3227042 | 7450 |
| 344 | 3227976 | 3232904 | 4929 |
| 345 | 3233303 | 3239881 | 6579 |
| 346 | 3240011 | 3246028 | 6018 |
| 347 | 3246677 | 3250010 | 3334 |
| 348 | 3250038 | 3251369 | 1332 |
| 349 | 3251405 | 3260957 | 9553 |
| 350 | 3263575 | 3270212 | 6638 |
| 351 | 3285834 | 3297814 | 11981 |
| 352 | 3297932 | 3313339 | 15408 |
| 353 | 3315431 | 3353975 | 38545 |
| 354 | 3362373 | 3367848 | 5476 |
| 355 | 3369638 | 3385387 | 15750 |
| 356 | 3385817 | 3400625 | 14809 |
| 357 | 3400734 | 3406159 | 5426 |
| 358 | 3412196 | 3413425 | 1230 |
| 359 | 3413493 | 3416553 | 3061 |
| 360 | 3416784 | 3417144 | 361 |
| 361 | 3420568 | 3432489 | 11922 |
| 362 | 3432975 | 3446629 | 13655 |
| 363 | 3463504 | 3492974 | 29471 |
| 364 | 3505993 | 3524011 | 18019 |
| 365 | 3524074 | 3527799 | 3726 |
| 366 | 3527875 | 3536553 | 8679 |
| 367 | 3537433 | 3549149 | 11717 |
| 368 | 3553207 | 3556878 | 3672 |
| 369 | 3557494 | 3569024 | 11531 |
| 370 | 3569081 | 3572086 | 3006 |
| 371 | 3572310 | 3574986 | 2677 |
| 372 | 3578441 | 3585685 | 7245 |
| 373 | 3586084 | 3591727 | 5644 |
| 374 | 3591915 | 3606274 | 14360 |
| 375 | 3606329 | 3612349 | 6021 |
| 376 | 3619039 | 3623962 | 4924 |
| 377 | 3629568 | 3641148 | 11581 |
| 378 | 3643597 | 3644022 | 426 |
| 379 | 3647165 | 3648573 | 1409 |
| 380 | 3648615 | 3658328 | 9714 |
| 381 | 3659540 | 3662548 | 3009 |
| 382 | 3662952 | 3678978 | 16027 |
| 383 | 3682515 | 3689545 | 7031 |
| 384 | 3689818 | 3693447 | 3630 |
| 385 | 3693456 | 3699150 | 5695 |
| 386 | 3699458 | 3701677 | 2220 |
| 387 | 3701808 | 3703835 | 2028 |
| 388 | 3704159 | 3713960 | 9802 |
| 389 | 3715688 | 3720721 | 5034 |
| 390 | 3720767 | 3724125 | 3359 |
| 391 | 3724491 | 3734942 | 10452 |
| 392 | 3750036 | 3755315 | 5280 |
| 393 | 3763603 | 3764739 | 1137 |
| 394 | 3764742 | 3770387 | 5646 |
| 395 | 3770759 | 3782420 | 11662 |
| 396 | 3782407 | 3785911 | 3505 |
| 397 | 3787347 | 3789335 | 1989 |
| 398 | 3801900 | 3808450 | 6551 |
| 399 | 3808487 | 3810899 | 2413 |
| 400 | 3812234 | 3820651 | 8418 |
| 401 | 3820820 | 3825526 | 4707 |
| 402 | 3825579 | 3829676 | 4098 |
| 403 | 3833909 | 3835099 | 1191 |
| 404 | 3835310 | 3844086 | 8777 |
| 405 | 3844162 | 3846348 | 2187 |
| 406 | 3847282 | 3849667 | 2386 |
| 407 | 3857972 | 3863801 | 5830 |
| 408 | 3868681 | 3868776 | 96 |
| 409 | 3869500 | 3870834 | 1335 |
| 410 | 3871065 | 3877089 | 6025 |
| 411 | 3877383 | 3881552 | 4170 |
| 412 | 3881795 | 3890186 | 8392 |
| 413 | 3900213 | 3904885 | 4673 |
| 414 | 3905199 | 3918993 | 13795 |
| 415 | 3919372 | 3927130 | 7759 |
| 416 | 3927138 | 3934687 | 7550 |
| 417 | 3937055 | 3939984 | 2930 |
| 418 | 3940062 | 3953173 | 13112 |
| 419 | 3954037 | 3977554 | 23518 |
| 420 | 3979792 | 3979963 | 172 |
| 421 | 3980046 | 3987419 | 7374 |
| 422 | 3987882 | 3993505 | 5624 |
| 423 | 3993652 | 3995736 | 2085 |
| 424 | 3996648 | 4001737 | 5090 |
| 425 | 4001799 | 4008674 | 6876 |
| 426 | 4008714 | 4010552 | 1839 |
| 427 | 4010452 | 4028513 | 18062 |
| 428 | 4028891 | 4030577 | 1687 |
| 429 | 4030870 | 4033800 | 2931 |
| 430 | 4033879 | 4033994 | 116 |
| 431 | 4034266 | 4037404 | 3139 |
| 432 | 4037559 | 4049699 | 12141 |
| 433 | 4049985 | 4053590 | 3606 |
| 434 | 4068913 | 4071798 | 2886 |
| 435 | 4072657 | 4072869 | 213 |
| 436 | 4073659 | 4080209 | 6551 |
| 437 | 4086484 | 4086798 | 315 |
| 438 | 4087279 | 4096315 | 9037 |
| 439 | 4096397 | 4098442 | 2046 |
| 440 | 4098597 | 4108688 | 10092 |
| 441 | 4108773 | 4114384 | 5612 |
| 442 | 4114540 | 4117153 | 2614 |
| 443 | 4117214 | 4117822 | 609 |
| 444 | 4118006 | 4122195 | 4190 |
| 445 | 4122223 | 4143026 | 20804 |
| 446 | 4143624 | 4148151 | 4528 |
| 447 | 4148201 | 4150718 | 2518 |
| 448 | 4151052 | 4153502 | 2451 |
| 449 | 4155356 | 4156527 | 1172 |
| 450 | 4156587 | 4158140 | 1554 |
| 451 | 4158300 | 4158375 | 76 |
| 452 | 4158560 | 4161489 | 2930 |
| 453 | 4161567 | 4161682 | 116 |
| 454 | 4161985 | 4179501 | 17517 |
| 455 | 4180663 | 4188708 | 8046 |
| 456 | 4188718 | 4191747 | 3030 |
| 457 | 4193248 | 4197599 | 4352 |
| 458 | 4199702 | 4199777 | 76 |
| 459 | 4199962 | 4202892 | 2931 |
| 460 | 4202970 | 4203085 | 116 |
| 461 | 4203608 | 4208341 | 4734 |
| 462 | 4208524 | 4210260 | 1737 |
| 463 | 4212732 | 4217439 | 4708 |
| 464 | 4217659 | 4220070 | 2412 |
| 465 | 4220282 | 4221559 | 1278 |
| 466 | 4221812 | 4230127 | 8316 |
| 467 | 4230253 | 4230663 | 411 |
| 468 | 4232554 | 4240402 | 7849 |
| 469 | 4242328 | 4249049 | 6722 |
| 470 | 4251597 | 4254159 | 2563 |
| 471 | 4254242 | 4256789 | 2548 |
| 472 | 4257042 | 4258235 | 1194 |
| 473 | 4259342 | 4264589 | 5248 |
| 474 | 4264688 | 4266985 | 2298 |
| 475 | 4266988 | 4271557 | 4570 |
| 476 | 4273181 | 4275141 | 1961 |
| 477 | 4275341 | 4285722 | 10382 |
| 478 | 4286253 | 4286942 | 690 |
| 479 | 4287036 | 4294220 | 7185 |
| 480 | 4311514 | 4312239 | 726 |
| 481 | 4313153 | 4315558 | 2406 |
| 482 | 4316216 | 4323767 | 7552 |
| 483 | 4325511 | 4331445 | 5935 |
| 484 | 4331728 | 4333083 | 1356 |
| 485 | 4334746 | 4341479 | 6734 |
| 486 | 4352368 | 4362979 | 10612 |
| 487 | 4363182 | 4364179 | 998 |
| 488 | 4364446 | 4366516 | 2071 |
| 489 | 4366692 | 4373437 | 6746 |
| 490 | 4373656 | 4375158 | 1503 |
| 491 | 4375864 | 4382252 | 6389 |
| 492 | 4382400 | 4399823 | 17424 |
| 493 | 4399794 | 4399868 | 75 |
| 494 | 4402204 | 4414204 | 12001 |
| 495 | 4414333 | 4416374 | 2042 |
| 496 | 4418752 | 4421093 | 2342 |
| 497 | 4421138 | 4427312 | 6175 |
| 498 | 4428525 | 4438253 | 9729 |
| 499 | 4438939 | 4440735 | 1797 |
| 500 | 4440875 | 4449128 | 8254 |
| 501 | 4449285 | 4449672 | 388 |
| 502 | 4449717 | 4450181 | 465 |
| 503 | 4450339 | 4460138 | 9800 |
| 504 | 4460344 | 4462350 | 2007 |
| 505 | 4468290 | 4468706 | 417 |
| 506 | 4470799 | 4479880 | 9082 |
| 507 | 4485007 | 4486306 | 1300 |
| 508 | 4541193 | 4545166 | 3974 |
| 509 | 4547195 | 4549343 | 2149 |
| 510 | 4549814 | 4550497 | 684 |
| 511 | 4550527 | 4551301 | 775 |
| 512 | 4577726 | 4581096 | 3371 |
| 513 | 4581474 | 4583129 | 1656 |
| 514 | 4586967 | 4592675 | 5709 |
| 515 | 4593294 | 4595857 | 2564 |
| 516 | 4596132 | 4599254 | 3123 |
| 517 | 4599231 | 4604077 | 4847 |
| 518 | 4604497 | 4611419 | 6923 |
| 519 | 4612918 | 4618364 | 5447 |
| 520 | 4618672 | 4620339 | 1668 |
| 521 | 4620550 | 4631445 | 10896 |

## Discussion

For the purpose of synthetic engineering, determining the set of core regions for a given species is critical as changes to these regions should be expected to reduce fitness. Core regions indicate parts of the genome that are conserved across evolution within a species. These regions are not necessarily required for survival but presumably define the characteristic core genotype which produces the core phenotype (lifestyle). Since most essential gene studies are carried out under specific laboratory growth conditions, genes which would normally be essential for a species across a diverse set of environmental conditions might not be discovered (e.g., due to fluctuating temperatures). Correspondingly, genes required to out compete rival organisms through increased fitness or to evade immune responses might not be found under laboratory conditions. Core regions, therefore, should be a superset of essential genes in most cases but exceptions might occur for genes which are not needed in a species’ natural niche but are required in a laboratory setting. Another exception would be for genes which are essential for a particular strain but not for other strains due to the presence of compensating non-core genes.

Verification of core genes or regions requires experimental studies on specific genomes. However, it is cost prohibitive to do knockout studies on all strains of a pan-genome. One has to carefully choose a single genome as a representative of the entire pan-genome for the purpose of verifying the essentiality of core regions and/or the non-essentiality of noncore regions by experimental validation. However, given the diversity of most bacterial species it is unlikely that any one strain completely captures the capabilities of the species in all environmental conditions. Further, while there are clearly core genes/regions associated with viability for a species, other core regions probably contribute to a lesser degree to cell viability. For example, for the purpose of synthetic engineering, changes in these locations may reduce fitness by slowing cell growth.

The use of a PGG for identifying core regions of a bacterium is an automatable, low-cost, rapid, and effective way to evaluate both gram-negative and gram-positive bacteria. This method compliments and expands upon the experimental knockout approach by including environmental diversity as a measure of what regions and genes are conserved across the species. The approach also overcomes the limitations of knockout studies that are specific to the strains and conditions used among other issues. Core region analysis should inform the design of synthetic engineering.

The *B. subtilis* WTA region provides a cautionary note for relying entirely upon core regions to determine what is safe to remove. While most non-core regions involve cassettes of genes which are entirely absent from som strains such as phage regions, sometimes orthologous replacement possibly due to homologous recombination can have functionally equivalent genes appearing to be non-core. A closer examination of the PGG can determine if a region is simply missing from some strains versus being replaced in which case further study may be needed before removal of the region. Of course in some cases the orthologous replacement does not need to occur at the same location in the genome but that was the case for all instances in *B. subtilis*.

## Materials and Methods

For *B. subtilis* ssp. *subtilis* and *E. coli* we selected strains with complete genomes in RefSeq (20). We restricted our analysis to complete genomes to ensure that missing genes due to incomplete genome sequencing/assembly did not affect the approach or results. We limited our choice to RefSeq for two reasons: RefSeq performs a series of quality checks to remove dubious genome assemblies, and the initial pan-genome construction depends upon reasonably consistent annotation which RefSeq provides. We extracted the genomes based on organism name: *Bacillus subtilis* (we did not specify subspecies, since for many RefSeq genomes a subspecies is not given) and *Esherichia coli* (we also specified *Shigella* since all *Shigella* species are actually considered to be the same species as *Escherichia coli*) (21, 22). For each pan-genome we then compared the genomes using a fast Average Nucleotide Identity (ANI) estimate generated using MASH (23). We used type strains and ANI to determine which of these genomes were actually the desired organism. We also used ANI to remove very closely related strains to reduce oversampling bias (for example for the *B. subtilis* type strain, 168, has at least 8 genomes in RefSeq). We used GGRASP (24) to choose a single medoid sequence from any complete linkage ANI cluster with a threshold of 0.01% or 1/10,000 base pair difference. The strain 168 medoid genome is the Entrez reference genome for the *B. subtilis* type strain (GenBank sequence AL009126.3, BioSample SAMEA3138188, Assembly ASM904v1**/**GCA_000009045.1) which can be used to map the Kobayashi and Koo results.

Using this approach, for *B. subtilis* 143 genomes were downloaded from RefSeq. Of these 132 genomes were determined to be *B. subtilis* spp. *subtilis* based on type strains and ANI. The minimum ANI between any pair of the 132 *B. subtilis* spp. *subtilis* genomes was 97.28% whereas the maximum ANI of any of the 11 other genomes to the 132 genomes was 95.73%, providing good separation between the other subspecies. The 132 genomes were reduced to 109 genomes after removing redundant strains. Finally we removed strain delta6 (BioSample SAMN05150066) because it is known to have been engineered to remove multiple genes. Thus we were left with 108 *B. subtilis* genomes (Supplemental Table 2). For *E. coli* (and *Shigella*) we downloaded 1097 complete genomes from RefSeq. Of these, 1096 were determined using ANI to actually be *E. coli*. The non *E. coli* genome was clearly mislabeled as its maximum ANI to any other genome was 82.27%. The minimum pairwise ANI of any of the 1096 genomes was 95.53% which is not as tight as for *B. subtilis* spp. *subtilis* which is to be expected given that *E. coli* is a species grouping not a subspecies grouping. One could arbitrarily try to choose a tighter grouping around the K-12 reference genome but the pairwise ANI values of the other genomes compared to the K-12 reference genome vary continuously from 96.22% to 100% with no punctate break in the values. After removing redundancy 969 *E. coli* genomes remained. We added back in two redundant genomes: The K-12 Entrez *E. coli* reference strain MG1655 (BioSample SAMN02604091) and the K-12 strain BW25113 (GenBank sequence accession CP009273.1, GenBank Assembly accession ASM75055v1/GCA_000750555.1, GenBank BioSample accession SAMN03013572) used by Goodall. By using a 95% threshold for the number of genomes a gene must be in to be considered core, some small number of the 971 genomes (Supplemental Table 4) could be engineered to remove what are normally core genes and not affect the assignment of core genes.

For *B. subtilis* ssp. *subtilis* and *E. coli* initial pan-genomes were based on the RefSeq annotation of these genomes. The pan-genome was generated using the pan-genome pipeline at the J. Craig Venter Institute (JCVI) at the nucleotide level using default parameters (25). This produced ortholog clusters using gene context (26) as well as a Pan-Genome Graph (PGG) (7). The PGG has two main components: nodes representing genes, and edges representing the sequence between genes and the order and orientation of the genes in the genomes. We updated the code repository for the JCVI pan-genome pipeline with a script: iterate_pgg_graph.pl, which calls pgg_annotate.pl for the genomes in the existing PGG in order to ensure consistent annotation of the genomes and iterates until the PGG stabilizes. The script pgg_annotate.pl uses an existing PGG to assign regions of a genome to nodes of the graph. This is done by blasting the medoid sequence for the gene cluster the node represents against the genome and then uses Needleman-Wunsch (27) to extend the alignment if needed. If there are conflicting blast matches then the matches are resolved based on which matches are consistent with the structure of the PGG which encapsulates gene context across the entire pan-genome. Once the nodes of the PGG are mapped to each of the genomes in the pan-genome a new version of the PGG is intrinsic and then explicitly extracted. This process is iterated to stability. This ensures that each genome is consistently annotated so that genes missing from the original annotation of some genomes will be consistently annotated across all genomes.

The original and refined PGG statistics for *B. subtilis* and *E. coli* are provided in Table 5. The major goal of this reannotation and iteration until stabilization was to achieve consistent annotation across all genomes in the PGG leading to a more comprehensive and cohesive PGG. While the RefSeq annotations of these genomes tends to be highly consistent, many small genes are often arbitrarily called from genome to genome and even some common longer genes can occasionally be missed. There are three obvious points of improvement in the refined PGG for both the cluster and edge stats: the number of size 1 clusters/edges significantly decreased due to some dubious RefSeq gene calls being eliminated and some becoming shared with other genomes; the number of core clusters/edges significantly increased showing an improvement in the consistency of annotation across all genomes; and the number of genes/edges in clusters/edge instances greatly increased again indicating a much more consistent annotation.

**Table 5.** Pan-genome graph statistics for *B. subtilis* and *E. coli*.

| **PGG Statistic** | ***B. subtilis***<br>**original PGG** | ***B. subtilis***<br>**refined PGG** | ***E. coli***<br>**original PGG** | ***E. coli***<br>**refined PGG** |
| --- | --- | --- | --- | --- |
| Size 1 clusters | 4434 | 3231 | 87423 | 27273 |
| Shared (size>1) clusters | 8174 | 8204 | 68970 | 48129 |
| # of genes in shared clusters | 463311 | 487562 | 5039502 | 5610683 |
| Core clusters | 3558 | 3778 | 2968 | 3631 |
| Size 1 edges | 7282 | 5479 | 153199 | 67566 |
| Shared edges | 9755 | 9433 | 99284 | 67248 |
| Edge instances in shared edges | 460452 | 485177 | 4970823 | 5567497 |
| Core edges | 3230 | 3520 | 2218 | 3124 |

Core regions were determined based on the PGG. Nodes in the PGG were orthologous clusters of genes. Edges in the PGG represented adjacency of genes (contained in the clusters) in the underlying genomes. The definition of which genes were or were not considered “core” was determined relative to a threshold criterion. We used a criterion for core such that 95% or more of the underlying genome had to contain the cluster or edge. Considering that we used only complete genomes it might have been possible to use a 100% threshold. However, we opted for a 95% threshold based on prior experience and an abundance of caution to not under call core genes/edges. Each core region began with a core cluster followed by a core edge (if possible – otherwise the core region comprises a single cluster) to another core cluster and so on until a core edge cannot be found to continue the core region. A core region is just a path in the PGGwhich was then mapped onto any particular genome to determine the core region coordinates. When the core threshold was below 100% any particular genome may be missing a cluster (gene) or edge along this path which results in the path being broken into its remaining constituent parts.

In order to compare core regions to experimentally determined essential genes we needed a common base of reference. For each of the experimental studies, the genes are specified based on a reference strain that was used for the experiments and has a complete genome in RefSeq. For Kobayashi et al., only gene symbols/names were given which we mapped to Entrez GeneIDs using Entrez search. GeneIDs with no matches were manually curated to estimate the best matching gene symbol listed in the literature. For Koo, locus IDs were provided giving direct access to the gene coordinates for RefSeq accession NC_000964.3 (BioSample SAMEA3138188, Assembly GCF_000009045.1). For Goodall, we used the data from three studies in Table S2 from Goodall et al. Gene symbols/names again were all that was available but these were consistent with the GenBank annotation downloadable in gff format for the K-12 BW25113 reference genome (GenBank accession CP009273.1) used by Goodall (BioSample SAMN03013572). This gave us coordinates for all essential genes on RefSeq genomes which were annotated with a PGG which produces a file with coordinates for clusters and edges mapped to the genome. These coordinates allow us to affiliate essential genes to gene clusters.

## Acknowledgements

This research is based upon work supported [in part] by the Office of the Director of National Intelligence (ODNI), Intelligence Advanced Research Projects Activity (IARPA) under Finding Engineering Linked Indicators (FELIX) program contract #N6600118C-4506. The views and conclusions contained herein are those of the authors and should not be interpreted as necessarily representing the official policies, either expressed or implied, of ODNI, IARPA, or the U.S. Government. The U.S. Government is authorized to reproduce and distribute reprints for governmental purposes notwithstanding any copyright annotation therein.

The authors would like to thank Derren Barken for his assistance in table generation.

---

**Supplementary Table 1**. All *B. subtilis* genes compared to the PGG based annotation of core regions for the type strain genome used by Kobayashi et al and Koo et al. (strain 168, GenBank sequence AL009126.3, BioSample SAMEA3138188, Assembly ASM904v1**/**GCF_000009045.1). Columns 1-5 are the start, stop, strand, cluster, and cluster size for the PGG annotation. Columns 6-11 are the gene type, start, stop, strand, locus tag, and gene symbol/name for the GenBank annotation. Column 12 is the Koo et al. gene symbol/name. Columns 13-14 are the Kobayashi et al. gene symbol/name and evidence type (from Supporting Table 4 “RB, reference to study with Bacillus subtilis ; RO, reference to study with other bacteria; TW, this work; TW*, inactivation failed but IPTG mutant could not be made”). Columns 15-16 are the GenBank protein product accession and name. Column 17 is the PGG core or non-core region the gene is contained in.

**Supplementary Table 2.** The 108 *B. subtilis* genomes used in the study. Data is from GenBank RefSeq: BioSample ID, Assembly ID, GenBank Species, GenBank Strain, Genome SIaze, and whether the genome is a type strain.

**Supplementary Table 3.** All *E. coli* genes compared to the PGG based annotation of core regions for the K-12 BW25113 strain used by Goodall (GenBank sequence CP009273.1, Assembly ASM75055v1/GCA_000750555.1, BioSample SAMN03013572). Columns 1-5 are the start, stop, strand, cluster, and cluster size for the PGG annotation. Columns 6-11 are the gene type, start, stop, strand, locus tag, and gene symbol/name for the GenBank annotation. Column 12 is a list of gene synonyms for the gene from GenBank. Columns 13-21 are from Goodall et al.: 13-15 from Table S1 (normal essentiality), 16-18 from Table S4 (essentiality after outgrowth), 19-20 from Table S3 (outlier discrepancies), and 21 from Table S2 (comparison of data sets). Column 22 is the PGG core or non-core region the gene is contained in.

**Supplementary Table 4.** The 971 *E. coli* genomes used in the study. Data is from GenBank RefSeq: BioSample ID, Assembly ID, GenBank Species, GenBank Strain, Genome SIaze, and whether the genome is a type strain.

## References

1. Hutchison CA, Chuang RY, Noskov VN, Assad-Garcia N, Deerinck TJ, Ellisman MH, Gill J, Kannan K, Karas BJ, Ma L, Pelletier JF, Qi ZQ, Richter RA, Strychalski EA, Sun L, Suzuki Y, Tsvetanova B, Wise KS, Smith HO, Glass JI, Merryman C, Gibson DG, Venter JC. 2016. Design and synthesis of a minimal bacterial genome. Science 351: aad6253.

2. Kobayashi K, Ehrlich SD, Albertini A, Amati G, Andersen KK, Arnaud M, Asai K, Ashikaga S, Aymerich S, Bessieres P, Boland F, Brignell SC, Bron S, Bunai K, Chapuis J, Christiansen LC, Danchin A, Débarbouille M, Dervyn E, Deuerling E, Devine K, Devine SK, Dreesen O, Errington J, Fillinger S, Foster SJ, Fujita Y, Galizzi A, Gardan R, Eschevins C, Fukushima T, Haga K, Harwood CR, Hecker M, Hosoya D, Hullo MF, Kakeshita H, Karamata D, Kasahara Y, Kawamura F, Koga K, Koski P, Kuwana R, Imamura D, Ishimaru M, Ishikawa S, Ishio I, Le Coq D, Masson A, Mauël C, Meima R, Mellado RP, Moir A, Moriya S, Nagakawa E, Nanamiya H, Nakai S, Nygaard P, Ogura M, Ohanan T, O’Reilly M, O’Rourke M, Pragai Z, Pooley HM, Rapoport G, Rawlins JP, Rivas LA, Rivolta C, Sadaie A, Sadaie Y, Sarvas M, Sato T, Saxild HH, Scanlan E, Schumann W, Seegers JF, Sekiguchi J, Sekowska A, Séror SJ, Simon M, Stragier P, Studer R, Takamatsu H, Tanaka T, Takeuchi M, Thomaides HB, Vagner V, van Dijl JM, Watabe K, Wipat A, Yamamoto H, Yamamoto M, Yamamoto Y, Yamane K, Yata K, Yoshida K, Yoshikawa H, Zuber U, Ogasawara N. 2003. Essential *Bacillus subtilis* genes. Proc Natl Acad Sci U S A. 100: 4678–83.

3. Koo BM, Kritikos G, Farelli JD, Todor H, Tong K, Kimsey H, Wapinski I, Galardini M, Cabal A, Peters JM, Hachmann AB, Rudner DZ, Allen KN, Typas A, Gross CA. 2017. Construction and Analysis of Two Genome-Scale Deletion Libraries for *Bacillus subtilis*. Cell Syst. 4:291–305.

4. Goodall ECA, Robinson A, Johnston IG, Jabbari S, Turner KA, Cunningham AF, Lund PA, Cole JA, Henderson IR. 2018. The Essential Genome of *Escherichia coli* K-12. mBio. 20: e02096–17.

5. Baba T, Ara T, Hasegawa M, Takai Y, Okumura Y, Baba M, Datsenko KA, Tomita M, Wanner BL, Mori H. 2006. Construction of *Escherichia coli* K-12 in-frame, single-gene knockout mutants: the Keio collection. Mol Syst Biol. 2:2006.0008.

6. Yamazaki Y, Niki H, Kato J. 2008. Profiling of *Escherichia coli* Chromosome database. Methods Mol Biol. 416:385–389.

7. Chan AP, Sutton G, DePew J, Krishnakumar R, Choi Y, Huang XZ, Beck E, Harkins DM, Kim M, Lesho EP, Nikolich MP, Fouts DE. 2015. A novel method of consensus pan-chromosome assembly and large-scale comparative analysis reveal the highly flexible pan-genome of *Acinetobacter baumannii*. Genome Biol. 16:143.

8. Koskiniemi S, Lamoureux JG, Nikolakakis KC, t’Kint de Roodenbeke C, Kaplan MD, Low DA, Hayes CS. 2013. Rhs proteins from diverse bacteria mediate intercellular competition. Proc Natl Acad Sci U S A. 110:7032–7.

9. Holberger LE, Garza-Sánchez F, Lamoureux J, Low DA, Hayes CS. 2012. A novel family of toxin/antitoxin proteins in *Bacillus* species. FEBS Lett. 586(2):132–6.

10. Brantl S, Müller P. 2019. Toxin-Antitoxin Systems in *Bacillus subtilis*. Toxins 11: pii: E262.

11. Westers H, Dorenbos R, van Dijl JM, Kabel J, Flanagan T, Devine KM, Jude F, Seror SJ, Beekman AC, Darmon E, Eschevins C, de Jong A, Bron S, Kuipers OP, Albertini AM, Antelmann H, Hecker M, Zamboni N, Sauer U, Bruand C, Ehrlich DS, Alonso JC, Salas M, Quax WJ. 2003. Genome engineering reveals large dispensable regions in *Bacillus subtilis*. Mol Biol Evol. 20:2076–90.

12. Ohshima H, Matsuoka S, Asai K, Sadaie Y. 2002. Molecular organization of intrinsic restriction and modification genes *BsuM* of *Bacillus subtilis* Marburg. J Bacteriol. 184:381–9.

13. Brown S, Santa Maria Jr, JP, Walker S. 2013. Wall teichoic acids of gram-positive bacteria. Annu Rev Microbiol. 67:313–36.

14. D’Elia MA, Millar KE, Beveridge TJ, Brown ED. 2006. Wall teichoic acid polymers are dispensable for cell viability in *Bacillus subtilis*. J Bacteriol. 188:8313–6.

15. Henriques AO, Glaser P, Piggot PJ, Moran CP Jr. 1998. Control of cell shape and elongation by the *rodA* gene in *Bacillus subtilis*. Mol Microbiol. 28:235–47.

16. Lazarevic V. Abellan F-X, Möller SB, Karamata D, Mauël C. 2002. Comparison of ribitol and glycerol teichoic acid genes in *Bacillius subtilis*W23 and 168: Identical function, similar divergent organization, but different regulation. Microbiology. 148:815–24.

17. Ahn S, Jun S, Ro H-J, Kim JH, Kim S. 2018. Complete genome of *Bacillus subtilis* subsp. *subtilis* KCTC 3135^T^ and variation in cell wall genes of *B. subtilis* strains. J Microbiol Biotechnol. 28:1760–68.

18. Bindal G, Krishnamurthi R, Seshasayee ASN, Rath D. 2017. CRISPR-Cas-mediated gene silencing reveals RacR to be a negative regulator of YdaS and YdaT toxins in *Escherichia coli* K-12. mSphere. 2:e00483–17.

19. Kato J, Hashimoto M. Construction of consecutive deletions of the *Escherichia coli* chromosome. 2007. Mol Syst Biol 3:132.

20. O’Leary NA, Wright MW, Brister JR, Ciufo S, Haddad D, McVeigh R, Rajput B, Robbertse B, Smith-White B, Ako-Adjei D, Astashyn A, Badretdin A, Bao Y, Blinkova O, Brover V, Chetvernin V, Choi J, Cox E, Ermolaeva O, Farrell CM, Goldfarb T, Gupta T, Haft D, Hatcher E, Hlavina W, Joardar VS, Kodali VK, Li W, Maglott D, Masterson P, McGarvey KM, Murphy MR, O’Neill K, Pujar S, Rangwala SH, Rausch D, Riddick LD, Schoch C, Shkeda A, Storz SS, Sun H, Thibaud-Nissen F, Tolstoy I, Tully RE, Vatsan AR, Wallin C, Webb D, Wu W, Landrum MJ, Kimchi A, Tatusova T, DiCuccio M, Kitts P, Murphy TD, Pruitt KD. 2016. Reference sequence (RefSeq) database at NCBI: current status, taxonomic expansion, and functional annotation. Nucleic Acids Res. 44:D733–45.

21. Lan R, Reeves PR. 2002. *Escherichia coli* in disguise: molecular origins of *Shigella*. Microbes and Infection. 4: 1125–1132.

22. Meier-Kolthoff JP, Hahnke RL, Petersen J, Scheuner C, Michael V, Fiebig A, Rohde C, Rohde M, Fartmann B, Goodwin LA, Chertkov O, Reddy T, Pati A, Ivanova NN, Markowitz V, Kyrpides NC, Woyke T, Göker M, Klenk HP. 2014. Complete genome sequence of DSM 30083(T), the type strain (U5/41(T)) of *Escherichia coli*, and a proposal for delineating subspecies in microbial taxonomy. Stand Genomic Sci. 8:9:2.

23. Ondov BD, Treangen TJ, Melsted P, Mallonee AB, Bergman NH, Koren S, Phillippy AM. 2016. Mash: fast genome and metagenome distance estimation using MinHash. Genome Biol. 17:132.

24. Clarke TH, Brinkac LM, Sutton G, Fouts DE. 2018. GGRaSP: a R-package for selecting representative genomes using Gaussian mixture models. Bioinformatics. 34:3032–3034.

25. Inman JM, Sutton GG, Beck E, Brinkac LM, Clarke TH, Fouts DE. 2019. Large-scale comparative analysis of microbial pan-genomes using PanOCT. Bioinformatics. 35:1049–1050.

26. Fouts DE, Brinkac L, Beck E, Inman J, Sutton G. 2012. PanOCT: automated clustering of orthologs using conserved gene neighborhood for pan-genomic analysis of bacterial strains and closely related species. Nucleic Acids Res. 40:e172.

27. Needleman SB, Wunsch CD. 1970. A general method applicable to the search for similarities in the amino acid sequence of two proteins. J Mol Biol. 48:443–53.

